# Guide-tree bias of whole genome alignment can mislead phylogenomic analyses

**DOI:** 10.64898/2026.07.06.736671

**Authors:** Qixin Tao, Stefan Grünewald

## Abstract

Whole-genome alignment (WGA) is widely used for genome-scale phylogenetic inference, and most scalable WGA pipelines rely on progressive alignment guided by a pre-specified tree. Among progressive whole-genome aligners, Progressive Cactus is a successful state-of-the-art method. However, analyses of real and simulated avian data indicate that guide-tree choice can influence downstream tree inference; star guide trees do not remove this effect and can exacerbate long-branch attraction artefacts. We have developed a consensus strategy based on the Progressive Cactus framework by generating a small set of alternative guide-tree alignments and retaining only homology relationships consistently recovered across all alignments. In simulation experiments, consensus alignments improve precision, bring inferred site-pattern frequency distributions closer to those of the true alignments, and recover more true splits than single guide-tree alignments. In a real landbird (Telluraves) dataset, we observe a strong bias towards single binary guide trees and long-branch attraction for less resolved trees. While the reconstructed tree still depends on the phylogenetic method and taxon sampling, our consensus alignment has no clear bias. We implemented a hierarchical consensus workflow that only locally resolves uncertainty in the guide tree. Therefore, the computational cost increases only moderately, for example by an estimated 68 percent for a recently published large-scale alignment of more than 300 modern birds (Neoaves) taxa.

## Introduction

As sequencing costs continue to decline and telomere-to-telomere (T2T) assemblies become more routine, whole-genome data acquisition has shifted from a few species-specific engineering efforts to large-scale, cross-clade practice (Chan and Ragan 2013; Kille et al. 2022). These genome resources are mainly generated to support comparative and functional studies, including applications in drug development (Abadio et al. 2011), molecular breeding (Song et al. 2024), and conservation (Zoonomia Consortium 2020). For downstream analyses that require site-resolved homology, whole-genome alignment (WGA) remains essential. At the same time, WGA faces two major challenges: rapidly increasing data scale and widespread structural rearrangements, such as inversions and translocations, which make homology relationships discontinuous and increase computational burden (Kille et al. 2022). Developing scalable and reliable WGA therefore remains an important problem.

A central challenge is how to perform large-scale WGA under realistic computational constraints. Progressive alignment has long been widely used for this purpose: it uses a pre-specified guide tree to define merge order and then builds a multiple alignment through iterative pairwise steps (Feng and Doolittle 1987; Chao et al. 2022). This paradigm underlies many widely used protein- or gene-scale aligners (Higgins and Sharp 1988; Katoh et al. 2002; Edgar 2004) and genome-scale alignment frameworks (Blanchette et al. 2004; Darling et al. 2010). However, benchmarking studies have shown that large-scale and reliable WGA remains challenging to be built, especially at deeper phylogenetic distances and under complex genome structures (Earl et al. 2014).

In progressive whole-genome alignment, Progressive Cactus is a widely used state-of-the-art method. It follows the core guide-tree-driven design and progressively merges genomes through ancestral reconstruction (Armstrong et al. 2020). Compared with traditional reference-based pipelines, it is reference-free, making regions that are absent from or only partially represented in a single reference genome less likely to be missed in the final alignment. It also uses cactus graphs (Paten et al. 2011) to represent complex rearrangements and homology relationships shaped by sequence duplication, and stores results in an HAL file (Hickey et al. 2013). These features have supported major comparative genomics projects, including the Zoonomia mammalian framework (Christmas et al. 2023) and dense avian sampling in the Bird 10,000 Genomes (B10K) Project (Feng et al. 2020). The method has also been applied to diverse downstream applications, including distant hybridization and reticulate evolution, as well as local phylogenetic analyses of hybridization and introgression based on Cactus whole-genome alignments (Ren et al. 2025; Wang et al. 2025), rapid structural divergence (Ivankovic et al. 2024), trait-associated evolution of noncoding regulation (Tamagawa et al. 2024; Shakya et al. 2025), bacterial anti-phage defense systems (Wu et al. 2024), and genome annotation workflows (Freedman and Sackton 2025). Derivative workflows such as ACMGA (Zhou et al. 2024) further illustrate how Progressive Cactus can be adapted into specialized variants for plant pan-genome analysis.

Nevertheless, Progressive Cactus cannot fully avoid a general risk inherent to progressive algorithms. Progressive alignment relies on a guide tree to determine the merge order, so the chosen guide-tree topology directly affects early alignment steps. Once local errors are introduced, later stages provide limited global correction, allowing such errors to accumulate and propagate (Feng and Doolittle 1987; Boyce et al. 2015). Lake (1991) argued that alignment order itself can systematically bias inferred tree topology. This influence is not necessarily obvious from pairwise agreement statistics. High overall agreement among alignments generated under different guide trees does not guarantee downstream topological robustness, because even a relatively small subset of alignment differences may still alter the recovery of some nodes. Nelesen et al. (2008) found that such alignments can remain globally similar yet still yield markedly different downstream phylogenetic inferences. In an avian test, approximately 98.5% of aligned pairs were identical across guide-tree variants (Armstrong et al. 2020). Even so, the authors noted that Progressive Cactus, although robust to small perturbations in the guide tree, still depends on a reasonably accurate phylogenetic tree (Hickey et al. 2024). Consistent with this, HAlign-G (Zhang, Zhou, et al. 2025) demonstrated that improving guide-tree quality can improve both Progressive Cactus alignment quality and downstream phylogenetic performance.

Several methods have been proposed to address alignment uncertainty. Among them, consensus-based ideas are most closely related to our study. Early work showed that comparing multiple alignments obtained under different parameter settings can help identify unstable sites (Gatesy et al. 1993; Wallace et al. 2006; Collingridge and Kelly 2012), but the perturbations they consider usually involve gap penalties, substitution costs, and alternative aligners rather than guide-tree topology itself as the primary source of bias. GUIDANCE (Penn et al. 2010) is also relevant because it generates alternative MSAs under guide-tree and related perturbations, but it begins from an initial alignment, so dependence on the initial conditions is not fully eliminated. SATé (Liu et al. 2009) addresses a different aspect of the problem by explicitly linking tree estimation and alignment refinement, using the current tree to update the alignment and the updated alignment to re-estimate the tree. However, these methods were developed mainly for gene- or protein-scale MSAs rather than whole-genome progressive alignment. At WGA scale, generating and processing large ensembles of alternative alignments is often computationally prohibitive, which limits the direct applicability of these methods to guide-tree-induced artefacts in progressive WGA. Consequently, there remains a substantial methodological gap for approaches that can identify homology relationships robust to guide-tree perturbation while remaining computationally practical at genome scale.

Against this background, we propose a whole-genome consensus alignment strategy within the Progressive Cactus WGA framework to mitigate the influence of guide-tree choice. Starting from a partially resolved input tree, some relationships among the input genomes may remain unresolved, either among individual taxa or among higher-level units containing already resolved subclades. We randomly generate a small set of alternative guide trees around these unresolved relationships, inducing clearly different alignment orders and thus producing several alignment versions. The alternative guide trees differ only in how the unresolved taxa or clusters are connected, while preserving resolved relationships in the input tree. We then retain the homologous relationships that are consistent across all alignments to construct a consensus alignment based on reference coordinates, yielding a final consensus alignment in which the reference sequence is gap-free. This rationale is consistent with Lake’s argument that the common subset should be less affected by order bias (Lake 1991). In our setting, the perturbation arises primarily from guide-tree topology rather than from parameter variation.

Our implementation targets whole-genome data, so we keep the number of parallel alignments to a small constant to avoid an unmanageable computational cost. For datasets with larger numbers of taxa, we further propose a hierarchical strategy: the program automatically identifies unresolved polytomies (interior nodes with more than two children) in the partially resolved input global-guide tree, then constructs alternative guide trees and extracts consensus alignments for these unresolved units at lower levels, infers consensus ancestors, and propagates those ancestors upward to higher alignment layers. Because consensus columns are defined with respect to reference chromosome coordinates, we use the reference as the outgroup in every round of multiple-guide-tree alignments. Groups with fixed topology need to be aligned only once, and their alignments can be reused in subsequent steps. The resulting sub-consensus alignments and fixed-topology sub-alignments are finally regrafted onto the root-level alignment to restore the full taxon set. This decomposition both breaks the problem into smaller alignment tasks and confines the extra computation to unresolved polytomies, rather than requiring a global rerun of the entire tree with full taxon set, thereby keeping the approach computationally practical. We have implemented this workflow in a program available on GitHub: given a partially resolved tree as input, it generates the working directories for each alignment level, the corresponding multiple guide trees, and an instruction file with commands which can be directly copied and run in parallel. In our 43-taxon simulation, this hierarchical design required 2.58 times as much CPU time as a single fully resolved alignment, indicating a moderate computational increase.

Different downstream analyses may depend on different aspects of an alignment. For phylogenomic inference, overall alignment length or recall may not be sufficient to assess alignment quality. Most aligned positions are invariant or singletons, whereas parsimony-informative site patterns are relatively few but especially important for tree reconstruction (Francis and Canfield 2020). Favorable overall alignment statistics do not necessarily indicate accurate phylogenetic signal. In simulations, where the true alignment is available, this can be evaluated by comparing site-pattern distributions between inferred and true alignments. Under this criterion, retaining more aligned sequence is not always more important than preserving phylogenetically relevant signal. In this study we therefore assess alignment quality in relation to its suitability for phylogenetic reconstruction, rather than by maximizing alignment coverage alone.

We evaluate the approach in simulations anchored on a classic avian reference topology (Jarvis et al. 2014). We compare single guide-tree WGAs, including fully resolved binary trees and star-like trees, to the consensus alignments. We quantify performance using precision, Jensen-Shannon divergence between inferred and truth site-pattern distributions, and recovery of splits in the true tree. Across 100 replicated experiments, compared with single guide tree WGAs, the resulting consensus alignments recover true splits more often, show reduced guide-tree-induced bias, and are closer to the truth in both precision and divergence, indicating better agreement with the true alignment. In contrast, less-resolved star-like guide trees did not improve alignment accuracy relative to single fully resolved guide trees and could instead induce long-branch attraction (LBA) artefacts in downstream tree inference. Finally, applying the consensus strategy to an empirical eight-taxon avian dataset confirms the topology for relationships within Afroaves published by Stiller et al. (2024).

## Results

We used avian genomes from Jarvis et al. (2014), which were later also used in the large-scale avian phylogenetic analysis by Stiller et al. (2024), for the real data experiments. Details (number of scaffolds/contigs, sequence length, and missing data proportion) of the sequences for the ten species used here can be found in Supplementary Table 1. We generated simulated genomes using Evolver (http://www.drive5.com/evolver/) with model parameters similar to Armstrong et al. (2020); see Subsection 2.1 of Supplementary Material. If not stated otherwise, the taxa and edge lengths for the underlying true trees of the simulations are extracted from the tree in Figure S7 of Jarvis et al. (2014), and all edge lengths are rescaled by 0.3, because for the original edge the simulator did not always terminate successfully. Mobile elements were disabled in Evolver because simulations with mobile elements also sometimes failed.

We first establish that Progressive Cactus has a guide-tree bias for 4-taxon real data. We then evaluate our consensus strategy on simulated datasets for true trees with various taxon numbers and edge-length distributions and compare the performance with alignments based on a single resolved or unresolved guide-tree. We next assess the computational cost for a simulated 43-taxon dataset. Finally, we apply the consensus method to a real 8-taxon landbird (Afroaves) dataset.

### Guide Tree Bias of Progressive Cactus for Bird Data

As an example of a difficult but solved problem of phylogenetic reconstruction, we use two sets of four bird taxa. Both contain chicken as the outgroup and turkey vulture. Then we added bar-tailed trogon and cuckoo-roller (both belonging to the Cavitaves clade) and seriema and zebra finch (both belonging to the Australaves clade), respectively. We ran Progressive Cactus on these taxon sets using all three possible rooted binary trees of the ingroups as guide trees, producing three alignments, and then inferred a tree from each alignment using IQ-TREE v1.6.12 (Nguyen et al. 2015).

The inferred tree topology was always the guide tree, with an interior edge length close to 0.005 (see Figure 1A for the first taxon set and Supplementary Figure 4 for the other one). Optimizing model-parameters and edge lengths for the alternative trees shows that the tree topology that coincides with the guide tree always has the highest log-likelihood score and the longest interior edge length (see Supplementary Tables 2, 3).

**Figure 1.**
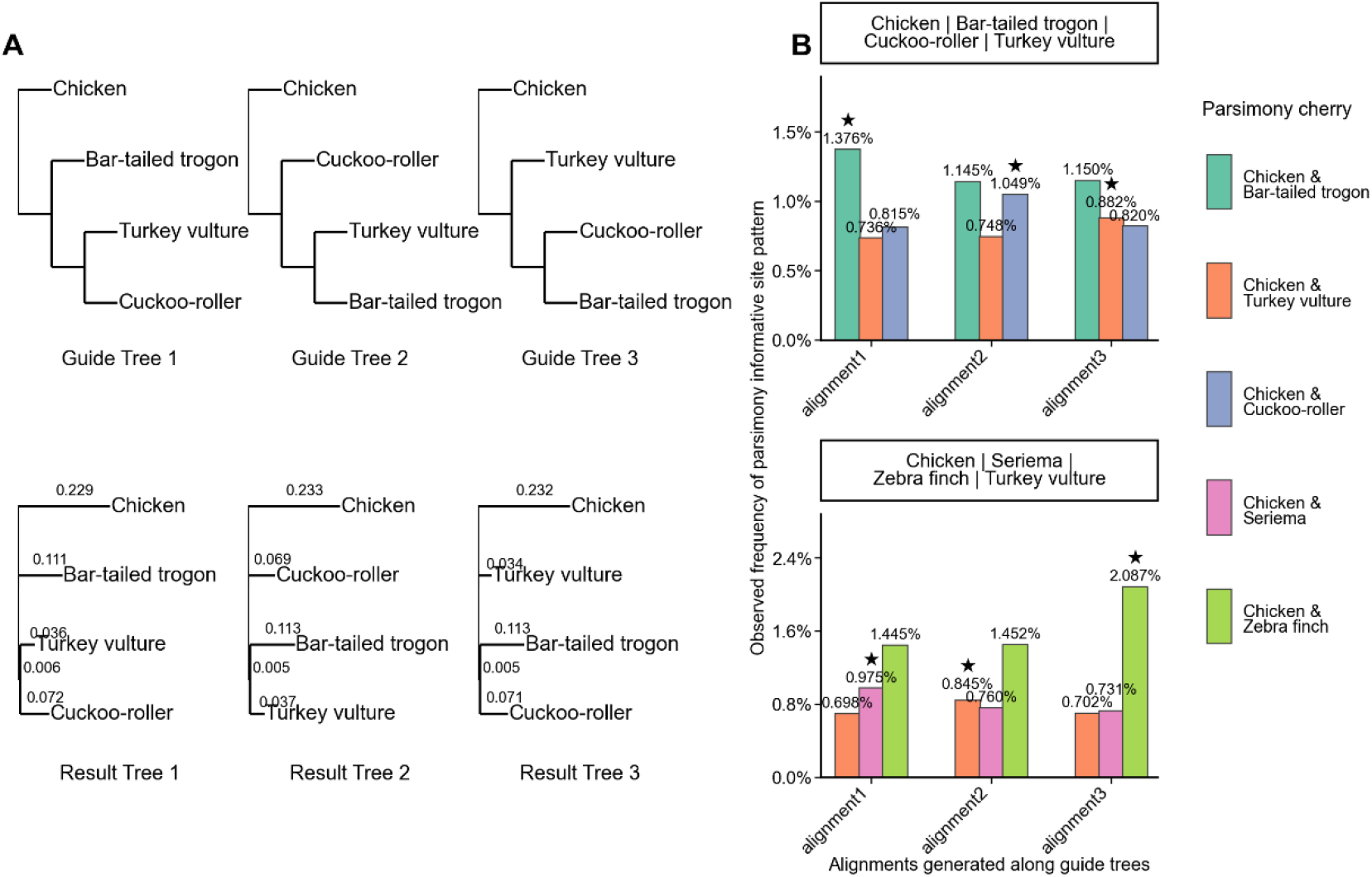
Evidence for guide-tree bias. **A**, Trees inferred from alignments generated under three binary guide trees for the quartet chicken, bar-tailed trogon, cuckoo-roller, and turkey vulture. **B**, Proportions of the three classes of parsimony-informative site patterns in each resulting quartet alignment.

Finally, we compared the proportions of three classes of parsimony-informative site patterns supporting the three topologies across the three alignments: for each class, its proportion is clearly higher in the alignment generated using the corresponding topology as the guide tree than in the other two alignments (Figure 1B).

Together, these results show that using a guide tree during alignment generation shifts the parsimony-informative site-pattern composition and makes downstream inference tend to recover the guide-tree topology.

### Performance comparison on simulation

#### Four-taxon datasets

We generate simulated genomes on a four-taxon set with four true-tree edge-length distributions. For four taxa, tree reconstruction is most challenging, if two of the pendant edges and the interior edge are short and the other two pendant edges are long. If there is one short and one long pendant edge on each side of the interior edge, then the tree is said to be in the Felsenstein zone (Felsenstein 1978) and there is a risk of long-branch attraction (LBA). If both short pendant edges are separated from the long ones by the interior edge, the tree is in the Farris zone, and long-branch repulsion (LBR) might happen. We consider a realistic scenario which was derived from a subtree of the main Jarvis tree (showing a slight LBA tendency). The other three scenarios are a tree deep in the Felsenstein zone, a tree in the Farris zone, and a clock-like tree (Figure 2A). For each scenario, we generated alignments using three fully resolved binary guide trees and a star guide tree, which are drawn in Supplementary Figure 5, and we further built consensus alignments: consensus3 is the consensus of the three binary-guide-tree alignments, and consensus4 adds the star-tree alignment into the consensus (i.e., a consensus across all four alignments).

**Figure 2.**
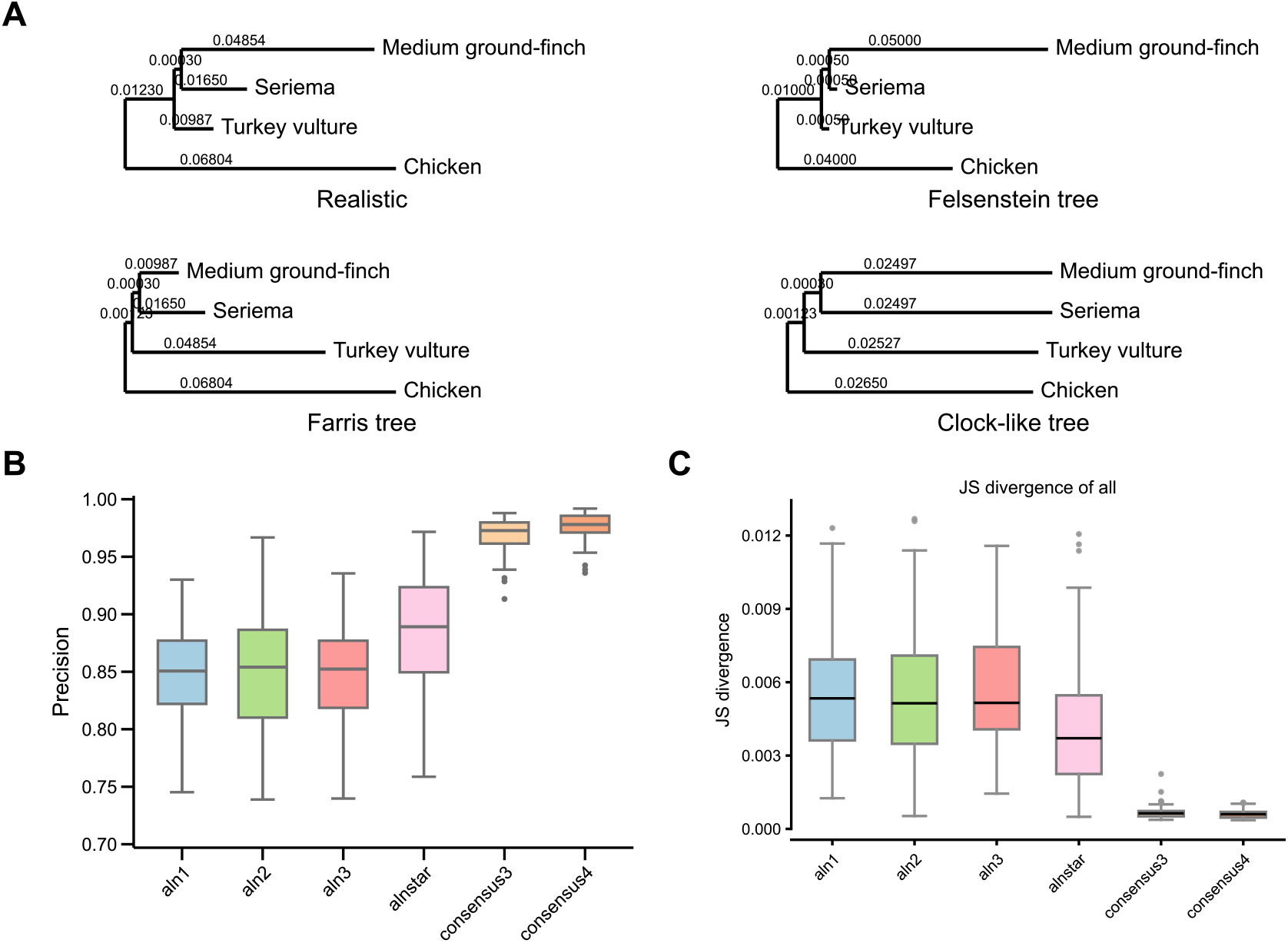
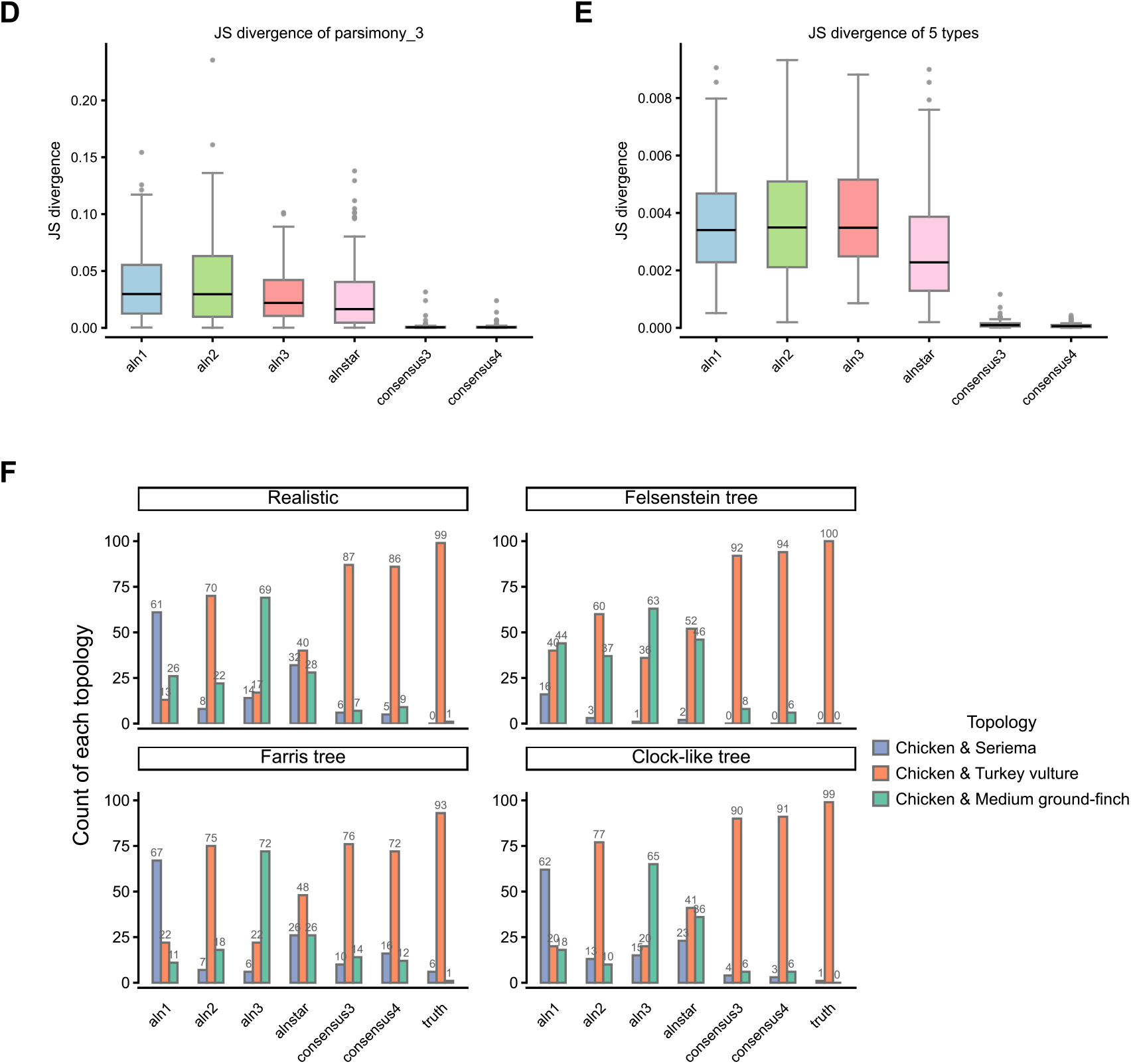
Simulation-based evaluation using four-taxon datasets. **A**, Simulation trees used by Evolver for the four evolutionary scenarios. **B**, Precision of guide-tree-based alignments and consensus alignments in the realistic scenario. **C-E**, Jensen-Shannon (JS) divergence calculated from all site patterns, the three classes of parsimony-informative site patterns, and all site patterns divided into five categories. **F**, Counts of inferred trees corresponding to the three alternative topologies for alignments generated under the four scenarios.

To quantify alignment quality, we computed alignment precision using the Alignathon definition (Earl et al. 2014) and Jensen-Shannon (JS) divergence measuring the distance between site-pattern frequency distributions from a reconstructed alignment and the true alignment. Because the four scenarios show similar trends for both precision and JS divergence, we report the realistic scenario in the main text and provide the other scenarios in Supplementary Figures 8-16. Consensus alignments achieve the highest precision (Figure 2B), indicating that they retain the largest fraction of true homologous pairwise matches with the true alignment. We further computed JS divergence at multiple levels, including individual site patterns, three parsimony-informative site-pattern classes supporting the three candidate topologies and five broader site-pattern types: invariant (all four taxa share the same nucleotide), singleton (3-1, e.g., AAAT), parsimony-informative (2-2, e.g., AATT), three-nucleotide 2-1-1 patterns (e.g., TTAG), and four-nucleotide 1-1-1-1 patterns (e.g., ATGC). Across these comparisons, consensus alignments consistently show the lowest JS divergence (Figures 2C-E, Supplementary Figure 7), meaning their site-pattern frequency distributions are closest to the true alignment across different ways of partitioning site patterns. In addition, using a star tree as the guide tree does not improve alignment quality: the star-tree-based alignment is slightly better than alignments generated from binary resolved guide trees in terms of precision or JS divergence, but the difference was very small compared to those for consensus alignments. As for all our experiments, the recall for consensus alignments is lower than for single-guide tree alignments (Supplementary Figure 6). Precision measures the fraction of correctly aligned pairs among all aligned pairs in the inferred alignment, whereas recall measures the fraction of aligned pairs in the true alignment that are correctly recovered (Earl et al. 2014).

Finally, we inferred phylogenetic trees using IQ-TREE from concatenated alignments and counted the frequencies of inferred topologies across 100 replicates for each guide-tree method and each consensus method (Figure 2F). Across all four scenarios, except for the true alignment, consensus alignments recover the correct topology most frequently, even more often than alignments generated using the true tree topology as the guide tree. In contrast, star-tree-based alignments tend to produce a Farris tree: the frequency of the Farris tree inferred from star-tree alignments is close to the frequency of the correct topology. Only when the Farris tree is the true tree, the star-tree alignment yields the correct topology for a clear majority of the samples, but it still does not outperform the consensus alignments.

#### Twelve-taxon simulation indicates using four guide trees as default

With more taxa, it becomes infeasible to enumerate all possible binary guide trees to generate many alignments and then take a consensus, due to time and computational cost. Therefore, we tested how many guide trees are needed to obtain a stable and high-quality consensus alignment in a multi-taxon simulation. Suh (2016) suggested a 9-way hard polytomy at the root of modern birds (Neoaves) evolution where two of the children of the root divide into pairs of other clades. For our 12-taxon experiment, we used the 11 categories defined by the Suh tree to select one representative taxon from each category, and added chicken. The simulated true tree for these taxa was then taken from the corresponding Jarvis TENT subtree (Figure 3E). The two clades that are fixed a priori are Telluraves (seriema and turkey vulture) and Phaethoquornithes (sunbittern and red-throated loon).

**Figure 3.**
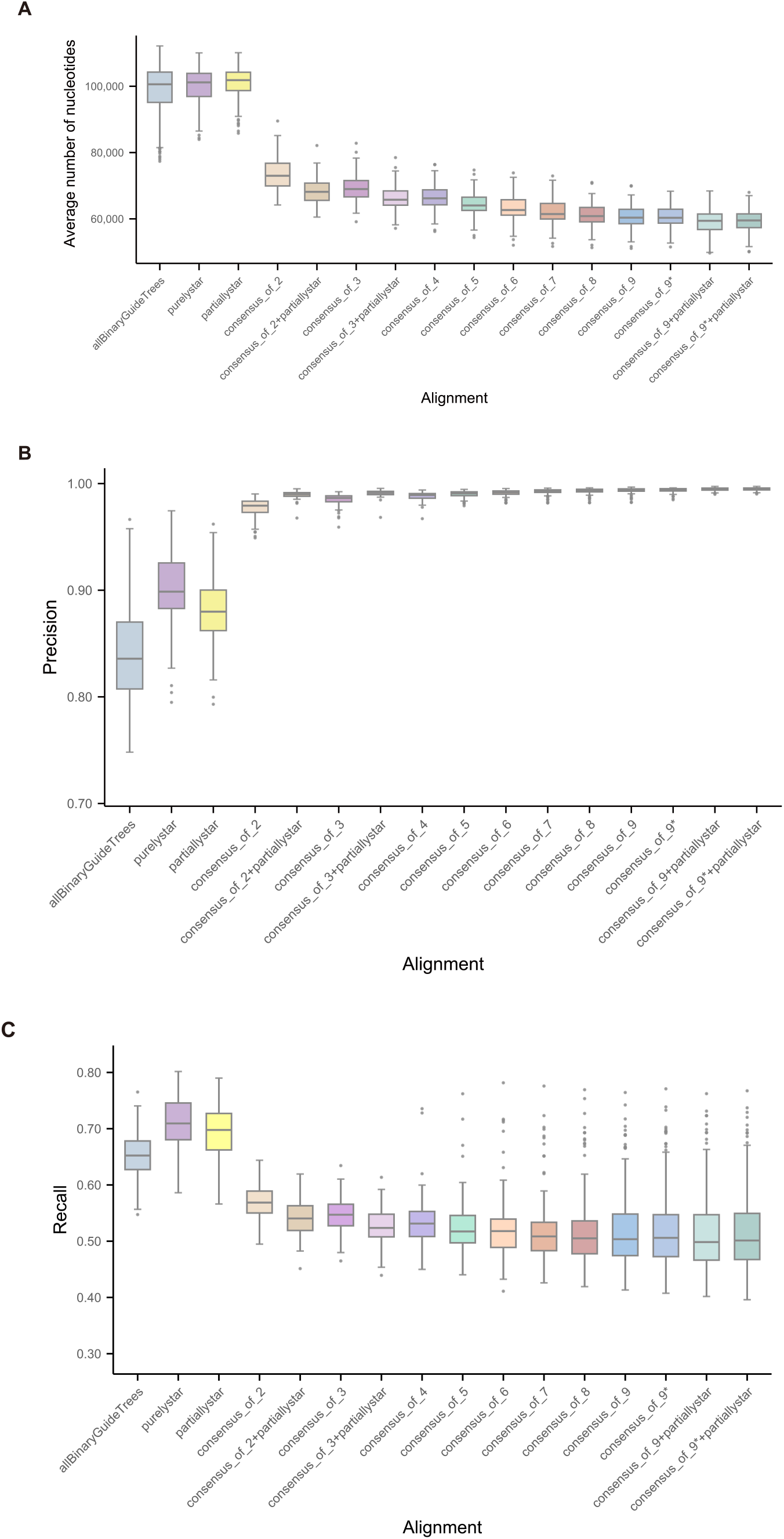
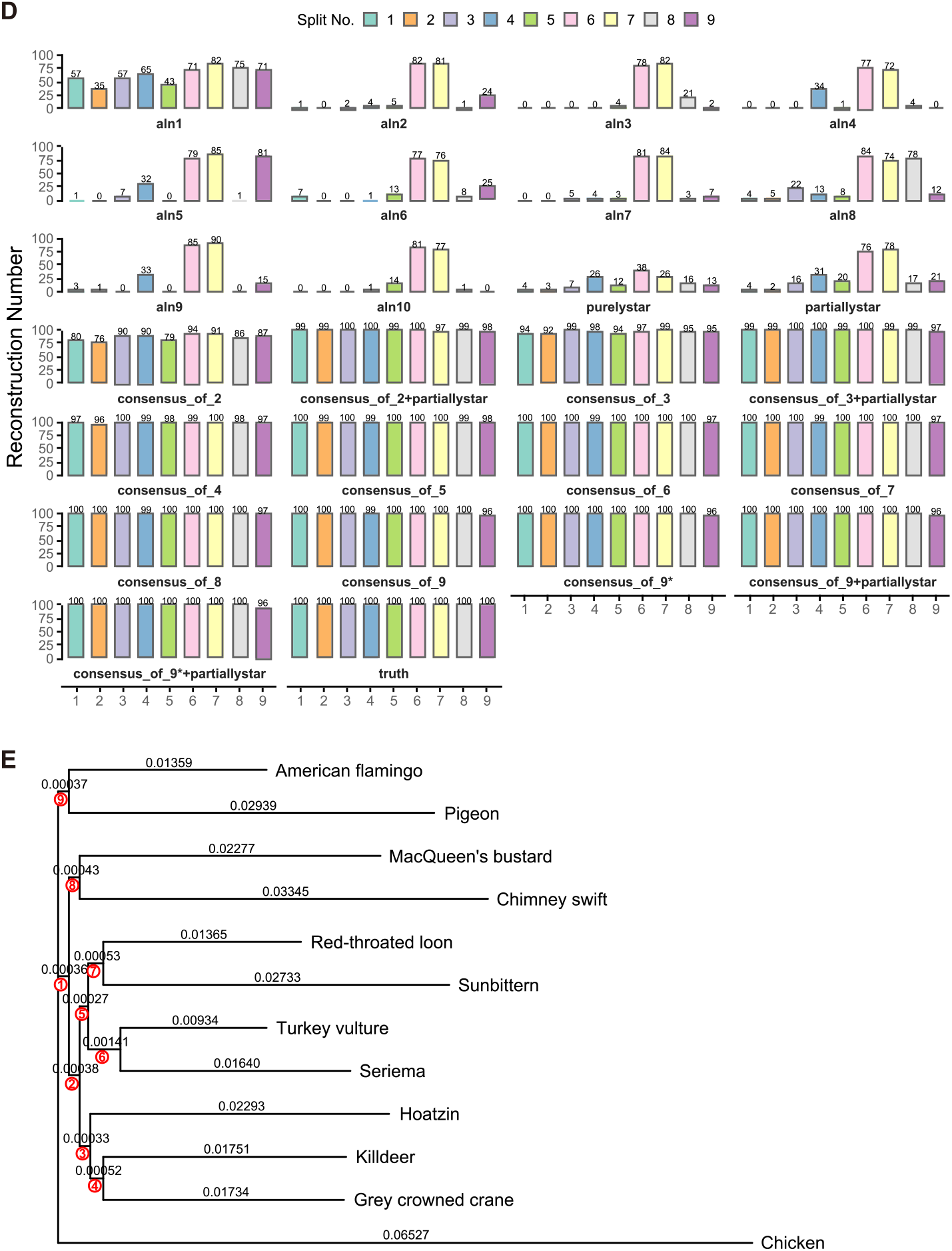
Simulation results for the 12-taxon dataset. An asterisk (*) indicates that the corresponding consensus alignment includes the alignment generated under the true tree as one of its input alignments. **A**, Average number of nucleotides of individual binary guide tree based alignments (pooled into a single box), the two star tree based alignments (shown separately), and consensus alignments constructed from different numbers of guide-tree-based alignments. **B**, Precision of individual binary guide tree based alignments (pooled into a single box), the two star tree based alignments, and consensus alignments constructed from different numbers of guide-tree-based alignments. **C**, Recall of individual binary guide tree based alignments (pooled into a single box), the two star tree based alignments, and consensus alignments constructed from different numbers of guide-tree-based alignments. **D**, Numbers of true-tree splits recovered from phylogenetic trees inferred from individual guide tree based alignments and consensus alignments. **E**, Simulation tree used in Evolver; red numbers indicate the split indices corresponding to those shown in D.

As guide trees, we used the true tree and nine additional fully resolved (binary) guide trees generated by our custom script under the default strict mode. This strict mode enforces two constraints: no overlapping clusters across guide trees except for the two fixed clusters, and a triplet distance (the proportion of three-taxon sets for which the trees induce different rooted triplets) at least 2/3 for any pair of guide trees. We also included two star-type guide trees: a proper star tree (no fixed clusters) referred to as “purely star”, and a partially resolved star tree with only the two fixed clusters. All guide trees are provided in Supplementary Figure 17.

We evaluated consensus alignments built from different numbers of guide trees using three metrics: alignment precision and average number of nucleotides, for which individual binary guide tree based alignments were pooled into a single box for comparison whereas the two star tree based alignments were shown separately, and tree accuracy measured by the number of true-tree splits recovered from IQ-TREE inference on concatenated alignments. In the plots consensus results marked with * indicate that the set of guide-tree alignments used for that consensus includes the alignment generated using the true tree as the guide tree. As more guide trees are used, average number of nucleotides (Figure 3A) and recall (Figure 3C) keep decreasing, while precision (Figure 3B) and tree accuracy increase (Figure 3D). Based on this trade-off, we chose a consensus built from four guide trees under the strict constraints as a practical default, as it yields a high-quality alignment and a high probability of recovering the correct topology. We also found that the consensus built from three fully resolved binary guide trees plus one partially resolved star tree (consensus_of_3+partiallystar) performed slightly better in both precision and tree reconstruction than the consensus built from four binary guide trees only (consensus_of_4). Based on these results, we chose to use four fully resolved binary guide trees for polytomies with at least four children, while we use all three rooted triples for polytomies with three children.

As a control experiment to check whether the actual consensus process or the conservative approach to keep only clearly supported sites in the alignment is responsible for the performance gain, we used three kinds of filters that reduce the average number of nucleotides to a value similar to consensus_of_4. We filter out paralogs (completely deleting multiple sequences from the same taxon within one aligned block), sites with any or at least two gaps, respectively, and short aligned blocks (deleting aligned blocks with length below 450 bp). Supplementary Figures 18-21 are analogues of Figures 3A-D, with filtered alignments included. Performance of the filtered alignments was quite similar to that of unfiltered single-guide tree alignments. The fraction of incorrectly aligned pairs among the removed parts relative to a single guide-tree alignment is shown for all consensus and filtered alignments in Supplementary Figure 22. For the consensus methods, this ratio is much higher than for the original alignments (ca. 50% versus 16%), while simple filters tend to have ratios close to 20%.

#### Simulated 43-taxon dataset

We performed simulations on a larger 43-taxon dataset, including all but one Neoaves taxon from Jarvis et al. (2014) plus chicken as the outgroup (Figure 4A, and drawn to scale in Supplementary Figure 23, splits are labeled from 1 to 40). Because bald eagle and white-tailed eagle are extremely closely related, we retained only bald eagle as a representative of this pair. By contracting fifteen very short edges, we get an unresolved global guide tree with an 8-way polytomy at the root, as well as two 5-way and a 4-way polytomies within landbirds. This tree is shown in Supplementary Figure 24, and all local guide trees used are shown in Supplementary Figures 25-29.

**Figure 4.**
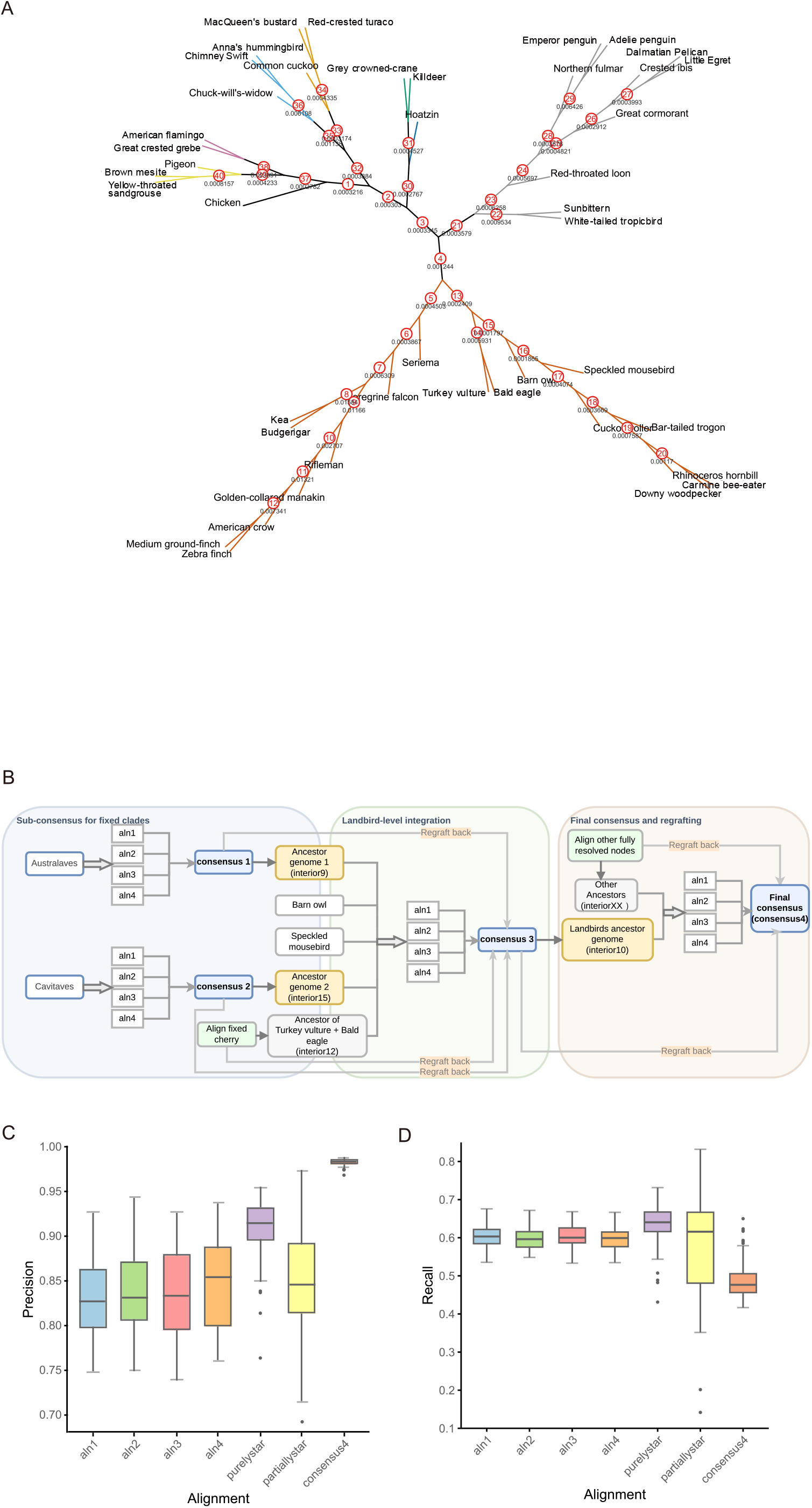
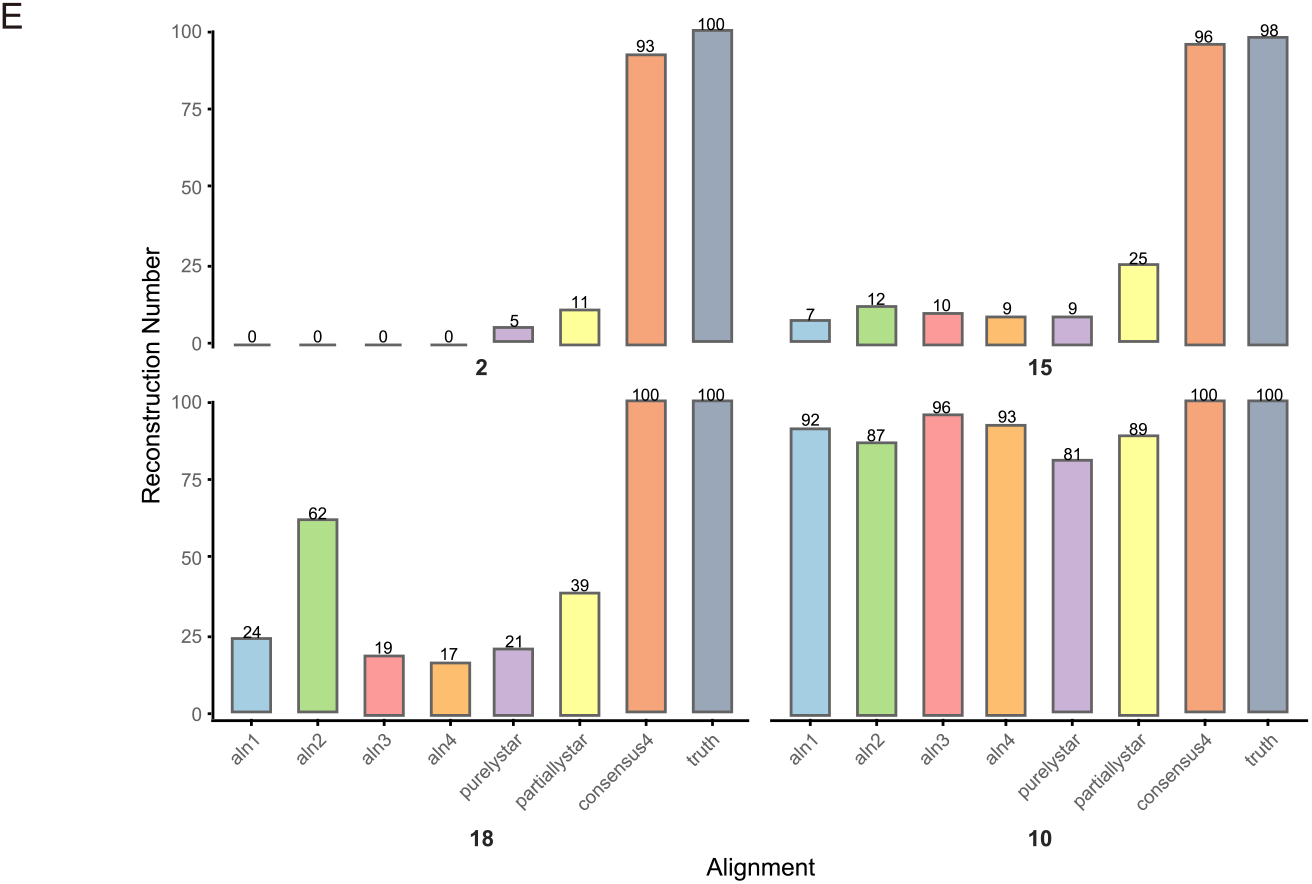
Hierarchical consensus alignment on the 43-taxon simulation dataset. **A**, Simulated species tree for the 43-taxon dataset, including all but one Neoaves taxon and chicken as the outgroup, the interior edges are labeled by their lengths and split indices. **B**, Hierarchical consensus pipeline applied to this dataset. **C**, Alignment precision of different strategies. **D**, Alignment recall of different strategies. **E**, Recovery of four representative true-tree splits in concatenation-based IQ-TREE analyses, ordered from more difficult to easier cases; the corresponding split indices in A are 2, 15, 18, and 10.

Following the default settings, we used four guide trees per polytomy for consensus extraction. The hierarchical consensus pipeline for this experiment is shown in Figure 4B. We include alignments where we use a single binary guide tree (“aln1” to “aln4”), two alignments where a global guide tree on all ingroup taxa is used (a star tree for “purelystar” and the partially resolved tree “partiallystar”), and the alignment where a consensus of the four guide trees is used for every polytomy (“consensus4”). For comparison with our hierarchical consensus pipeline, the four fully resolved guide trees for aln1-4 are obtained from the global guide tree by resolving all polytomies by the guide tree with the same index (Figure 4A). These guide trees are shown in Supplementary Figure 30.

We measure the accuracy of the competing strategies in terms of precision and frequency of correct splits reconstruction. Consensus4 achieved the highest alignment precision (shown in Figure 4C) with median above 0.983, and minimum of 0.968. Purelystar was second best with median of 0.915, while partiallystar had a median precision of 0.845. For binary guide trees, the median ranged from 0.827 to 0.854. The recall is also shown in Figure 4D, and the median of consensuse4 is 0.477, compared to 0.598 for single binary guide trees.

For each split of the true tree, we computed the number of correct reconstructions using concatenation-based IQ-TREE analyses (see Supplementary Material, Subsection 2.1, Point 7) for details). We highlight four representative splits in Figure 4E, ordered from more difficult to easier cases, and provide the full results in Supplementary Figure 34. Consensus4 was the only alignment strategy that recovered every split in more than 93 of 100 replicates. In most cases, partiallystar gave the second-best result. The main exception involved splits that were already present in a particular fully resolved guide tree, because the corresponding single guide-tree alignment could then gain a guide-tree-specific advantage.

The difference was largest for difficult splits associated with polytomy resolution. Split 2, which resolves the top-level polytomy, has a branch length of only 0.000303. None of the four fully resolved single guide-tree alignments recovered this split, and purelystar and partiallystar recovered it only 5 and 11 times, respectively. By contrast, consensus4 recovered it 93 times. Split 15 showed the same pattern. This lower-level landbird split, corresponding to Cavitaves + mousebird + barn owl, has the shortest internal edge in the tree (0.0001797), yet consensus4 still recovered it 96 times, far exceeding the best non-consensus result of 25, again from partiallystar.

The same advantage remained evident in less difficult but still informative cases. Split 18, corresponding to Cavitaves without cuckoo-roller, is present only in guide tree 2. As expected, aln2 recovered this split more often than the other single guide-tree alignments, consistent with a favorable guide-tree bias. Even so, consensus4 still outperformed aln2 for this split. A different pattern was seen for split 10, corresponding to Passeriformes without rifleman within Australaves. This split is present in all single guide trees except the purely star tree. Accordingly, all four fully resolved single guide-tree alignments recovered it at high frequency, whereas purelystar performed substantially worse. Consensus4 achieved 100 percent accuracy. These examples show that consensus4 improves recovery for the hardest splits and remains better even for splits already favored by one or more guide trees.

An alignment that uses the hierarchical pipeline but only based on a single guide tree is included in Supplementary Figures 31-34. This does not improve the performance.

We repeated the whole experiment using zebra finch as the reference. The precision and split reconstruction accuracy are shown in Supplementary Figures 35-37. The performance of consensus4 is similar and we conclude that the method is successful independently of the reference position (an outgroup to the taxa of interest or a taxon contained in a deep clade).

If the user would like to avoid a fixed reference, our method allows selecting a reference locally for every polytomy, always selecting the taxon with the longest genome. As shown in Supplementary Figures 33-34, this variant of the method performed slightly worse than one with a single global reference.

#### Computational cost

To obtain a practical runtime estimate for the hierarchical large-scale setting, we measured CPU time for the hierarchical 43-taxon simulation analysis on the 100 simulated datasets and report the mean across these repeats. We compared our consensus pipeline with purelystar, partiallystar and aln1-4 (represented as one average time) (Table 1). Values are mean CPU time, in seconds. For the consensus4 method, the total includes alignment of trusted nodes, four-way alignments within major subclades, conversion to MAF (multiple alignment format), which is commonly used for analysis tools, consensus extraction with ancestor inference, and regrafting of sub-consensus alignments back onto the final alignment.

**Table 1.** Mean CPU time in seconds for each step of the consensus pipeline and alternative 43-taxon alignment strategies across 100 simulation runs.

| Alignment method | Step |  | time/s | total/s |
|---|---|---|---|---|
| consensus4 | Align trusted nodes |  | 1543.34017 | 8378.86839 |
|  | Cavitaves | 4 alignments | 1391.17124 |  |
|  |  | Convert to MAF | 37.998 |  |
|  |  | Extract consensus+Infer ancestor | 143.07083 |  |
|  | Australaves | 4 alignments | 1081.65318 |  |
|  |  | Convert to MAF | 37.15408 |  |
|  |  | Extract consensus+Infer ancestor | 132.34952 |  |
|  | Landbirds | 4 alignments | 1338.18543 |  |
|  |  | Convert to MAF | 41.15662 |  |
|  |  | Extract consensus+Infer ancestor | 141.53045 |  |
|  | Top-level | 4 alignments | 2263.54913 |  |
|  |  | Convert to MAF | 42.33427 |  |
|  |  | Extract consensus+Infer ancestor | 171.64501 |  |
|  |  | regraft sub-consensus | 13.73046 |  |
| partiallystar | 43-taxon alignment |  | 2868.6194 |  |
| purelystar | 43-taxon alignment |  | 5595.14676 |  |
| Align with one totally resolved tree (Average of aln1-4) | 43-taxon alignment |  | 3245.861 |  |
| Only hierarchical pipeline | Align trusted nodes | 1543.34017 | 2855.40972 |  |
|  | Cavitaves | 261.48008 |  |  |
|  | Australaves | 186.3673 |  |  |
|  | Landbirds | 271.25241 |  |  |
|  | Top-level | 565.16633 |  |  |
|  | Regraft sub-alignments | 27.80343 |  |  |

Relative to the fully resolved guide-tree baseline, our consensus pipeline required 2.58 times the CPU time. This increase was markedly smaller than the fourfold increase that would be expected, if the four guide-tree-specific alignments were computed independently, because trusted nodes were aligned once and then reused across the four guide-tree-specific runs rather than being recomputed in every run. In the global guide tree shown in Supplementary Figure 24, we have 4 polytomies: two 5-way polytomies, one 4-way polytomy and one 8-way polytomy, and we have 28 interior vertices totally. Using Formulae (1) and (2) in Materials and Methods, a simple node-count approximation predicts that our pipeline requires 2.64 times as much CPU time as a single fully resolved alignment, assuming that binary nodes with fixed topology are aligned only once and that the four-guide-tree consensus procedure is applied to polytomies with at least four children. The additional cost was driven primarily by the extra Cactus alignment runs required for unresolved nodes, whereas consensus extraction itself contributed only a small fraction of the total CPU time.

### Application to a real eight-taxon landbird dataset

We applied the consensus-based alignment strategy to a real eight-taxon landbird dataset, including five taxonomic units: chicken as the outgroup, Accipitrimorphae (turkey vulture and white-tailed eagle), Cavitaves (bar-tailed trogon, cuckoo-roller and downy woodpecker), speckled mousebird, and barn owl. This empirical analysis involves only one unresolved ingroup node: the global guide tree contains a single four-way polytomy among Accipitrimorphae, Cavitaves, mousebird, and barn owl. We generated alignments under four binary guide trees (Figure 5A) (“aln1” to “aln4”) and extracted a consensus alignment across them (“consensus4”). We also included a purely star tree (“purelystar”), and a partially resolved tree that fixed the Accipitrimorphae clade and the pre-resolved Cavitaves topology (“partiallystar”). Generating the consensus4 alignment took 4331 CPU hours while alignments based on single guide tree took around 2700 CPU hours each (see Supplementary Table 4 for a complete list of CPU/memory usage).

**Figure 5.**
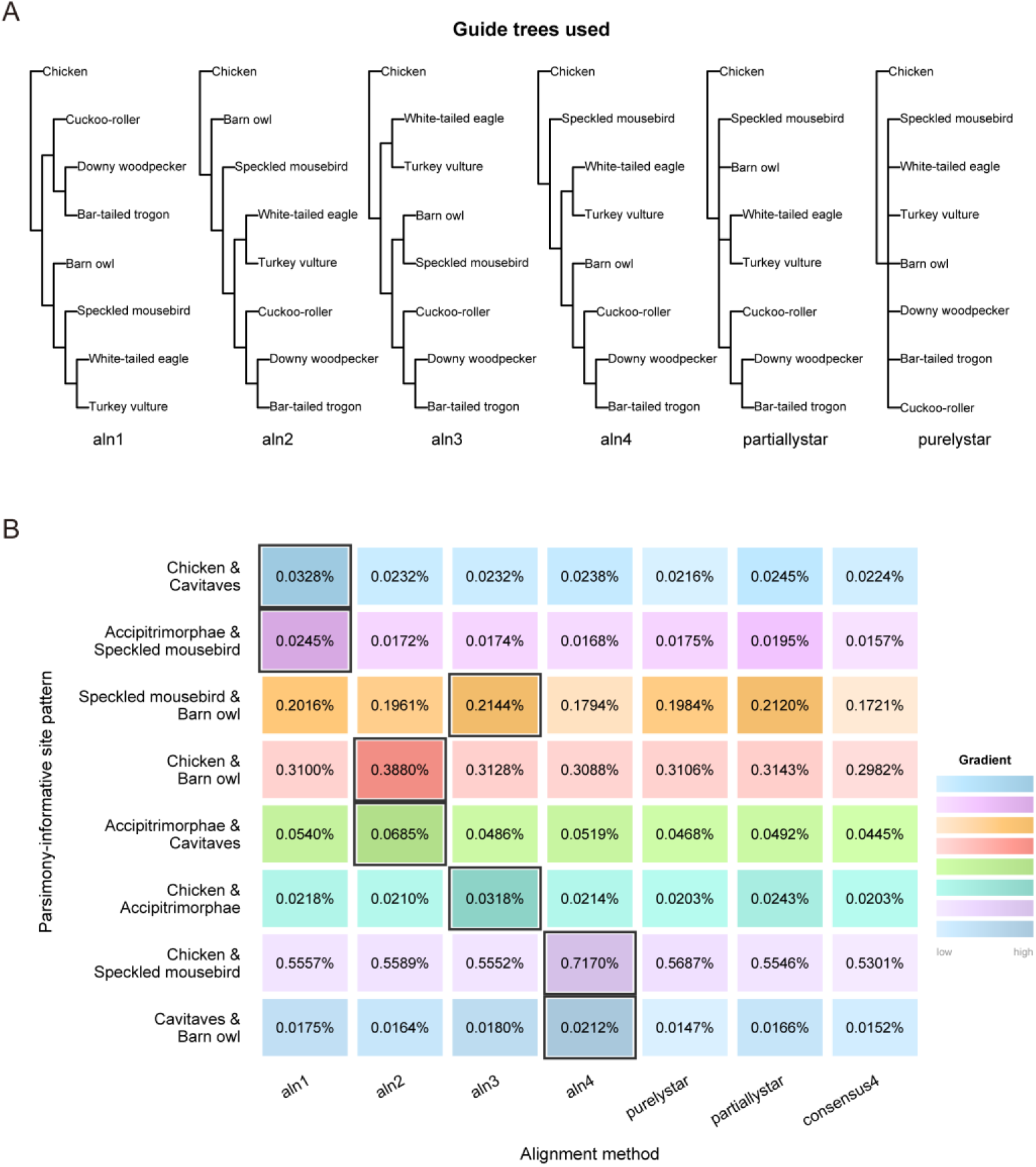
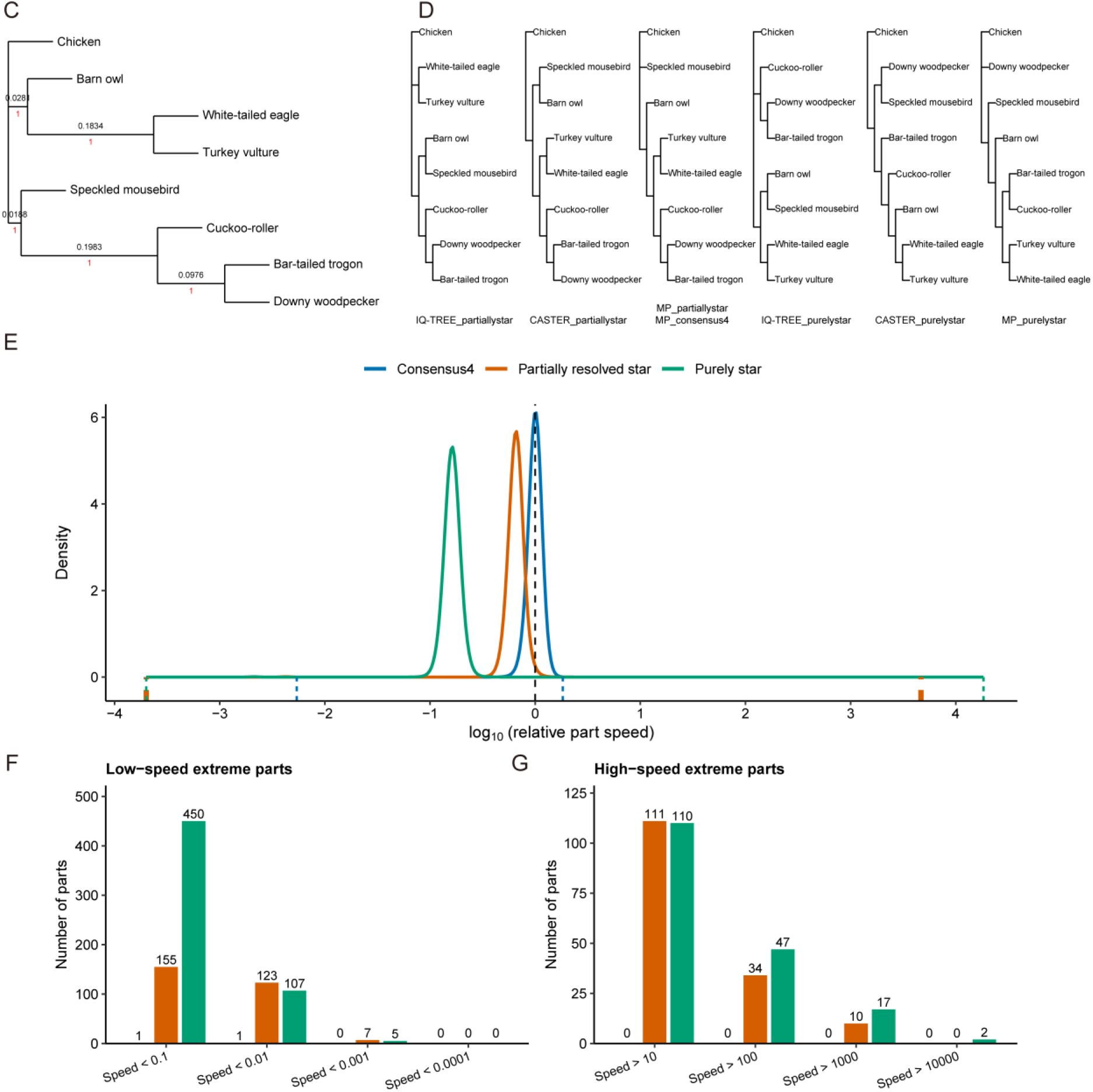
Application of the consensus method to a real eight-taxon avian dataset. **A**, Guide-tree topologies used for progressive alignment. **B**, Frequencies of parsimony-informative site patterns supporting the eight 2|3 classes among the five predefined groups across guide-tree-based, star-tree-based, and consensus alignments. **C**, Main tree inferred from the consensus alignment by loci-ASTRAL. Numbers in red indicate local posterior probabilities, whereas numbers in black indicate internal branch lengths in coalescent units, both estimated by ASTRAL. **D**, Alternative tree topologies reconstructed from the consensus4, partiallystar, and purelystar alignments. The six topologies shown comprise all tree topologies recovered by the alignment/reconstruction method combinations other than the main topology shown in (C). Drawn-to-scale trees with branch support values are shown in Supplementary Figures 38-41. **E**, Density distributions of relative evolutionary rates (“part speeds”) estimated for 10-kb blocks in the consensus4, partiallystar, and purelystar alignments. Part speed is expressed relative to the average rate of the entire alignment and is shown on a log10 scale. Dashed lines at both ends of each curve mark the minimum and maximum values. **F**, Numbers of 10-kb blocks with extremely low speed in the three alignments. **G**, Numbers of 10-kb blocks with extremely high speed in the three alignments.

Guide-tree effects seen in simulations were also detectable in the empirical dataset. We summarized the alignment at the level of the five units. We counted gap-free site patterns with exactly two nucleotides that were parsimony-informative for one of the eight 2|3 splits among units (Figure 5B) that occur in the four guide trees. For all eight 2|3 splits, the corresponding pattern class was most frequent in the alignment generated from the guide tree containing that same relationship. Thus, guide-tree bias is also evident in this empirical dataset: single guide tree alignments systematically enrich the site-pattern signal associated with the topology used to construct the alignment.

Phylogenetic reconstruction in this part of the landbird tree is inherently difficult. Previous studies have reported different resolutions, and differences in taxon sampling can further affect the inferred topology. In our empirical comparison, however, all alignments used the same eight-taxon set, allowing us to focus on alignment-dependent differences rather than taxon-sampling effects. Incomplete lineage sorting (ILS), in which ancestral genetic variation is not fully sorted during rapid successive speciation events, can cause individual gene trees to differ from the species tree and complicate phylogenetic inference (Feng et al. 2022). ILS has also been recognized as an important source of phylogenetic conflict in avian radiations (Suh et al. 2015), motivating the use of a locus-based summary-tree approach. Long-branch attraction is another relevant concern for this dataset. In particular, the long branch leading to downy woodpecker may create an LBA-like signal that pulls it towards the outgroup chicken rather than grouping it with other Cavitaves taxa.

We next examined how these alignment differences affect downstream tree inference using multiple inference strategies. Each alignment was analyzed using three primary pipelines: a Stiller-inspired loci/supertree pipeline (Stiller et al. 2024), a partitioned maximum likelihood analysis using consecutive 10-kb blocks with IQ-TREE v1.6.12 (Nguyen et al. 2015), and CASTER-site (Zhang, Nielsen, et al. 2025). In the Stiller-inspired pipeline, the alignment was partitioned into consecutive 10-kb loci along continuous chicken chromosomes’ coordinates. For each locus, a tree was inferred with IQ-TREE from the highest-occupancy (i.e., least missing data) 1-kb window within the first 5 kb, and these locus trees were then summarized with ASTRAL-III v5.7.8 (Zhang et al. 2018). For the concatenated maximum likelihood analysis, we partitioned alignments into consecutive 10-kb blocks. CASTER-site offers a recently proposed alternative based on a substantially different inference strategy and addresses both, ILS and rate heterogeneity. Although LBA is easy to illustrate in four-taxon examples, it is less transparent in larger taxon sets; we additionally ran maximum parsimony (MP) with PAUP* (Swofford 2001) as an LBA-sensitive reference for recognizing LBA-like behavior across trees inferred from different alignments. Full tree results for all alignments under IQ-TREE partition analysis, CASTER-site, MP, and the loci-ASTRAL pipeline are provided in Supplementary Figures 38-41. Alignment details (length, average nucleotides, and locus numbers, corresponding to the numbers of trees input to ASTRAL) are given in Supplementary Table 4.

Considering only the three alignments without single binary guide tree, a main tree topology, shown in Figure 5C was built by five out of the twelve alignment/reconstruction method combinations (the three non-parsimony methods for the consensus alignment, and ASTRAL for all alignments). The main tree topology happens to be identical with the Afroaves subtree of Stiller et al. (2024). Another tree topology is reconstructed by maximum parsimony for consensus4 and partiallystar. Five different tree topologies are produced by the remaining combinations, and all alternative trees are depicted in Figure 5D.

#### Purely star alignment tends to produce LBA trees

The purely star alignment shows a tendency of long-branch attraction. It produces a typical LBA tree with a cluster of mousebird and downy woodpecker for CASTER-site, and it yields unusually high proportions of LBA-preferred splits for the locus trees contributing to ASTRAL. While the supertree agrees with the other alignments, locus trees include almost 12% with a chicken–downy woodpecker cluster (9% and 8.3% for consensus4 and partiallystar, respectively), and 15.2% with a mousebird–downy woodpecker cluster (11% and 10% for consensus4 and partiallystar, respectively). On the other hand, the correct Cavitaves clade that includes the long-branched downy woodpecker and the short branched cuckoo-roller is only observed for 9.3% of the loci (16.3% and 16.2% for consensus4 and partiallystar, respectively).

#### The speed distribution across the 10-kb blocks has the lowest variance for consensus4

Our partitioned maximum likelihood analysis assigns a speed (relative rate to the whole alignment) to every 10-kb block. The distribution of the speed is narrow for consensus4 (all but one block have a speed between 0.3 and 2). For partiallystar and purelystar, there are many blocks with extremely low or extremely high speed. A density plot for all three alignments is shown in Figure 5E and outlier counts are visualized in Figures 5F and 5G. As a consequence, the edges of the reconstructed trees in Supplementary Figure 38 are much longer for purelystar and partiallystar than consensus4 (for example, 0.459 for purelystar, 0.111 for partiallystar, 0.07 for consensus4 for the cuckoo-roller branch, which has length 0.067 in the TENT tree in Jarvis et al. (2014)). The outliers mainly stem from the chicken microchromosomes, often with high gap frequency. While a preprocessing step filtering out such blocks might improve the speed distribution for competing methods, we conclude that consensus4 almost completely avoids such potentially biasing blocks.

#### Differences compared to a previous analysis of chicken chromosome 1 only

In an earlier version of this manuscript, we analyzed whole chromosome alignments for chicken chromosome 1 and same taxon set as here, using an older version of Progressive Cactus, and without soft masking repeats. We found that, for most reconstructed trees, the mousebird was classified as the outgroup to other Afroaves. The per-chromosome statistics for the current experiment shows that this mousebird position is indeed supported more by chicken chromosome 1 than for the rest of the genome (see Supplementary material for details). In addition, the other two changes to the pipeline weakened the support even on chromosome 1. This demonstrates that other difficulties in phylogenetic reconstruction like conflicting signals across chromosomes remain, even if whole genome alignments improve.

## Discussion

Our results show that guide-tree choice in progressive whole-genome alignment can bias downstream phylogenetic inference. This bias is reproducible, and can be particularly important when phylogenetic signal is weak. To reduce dependence on any single guide tree, we generated replicate alignments under multiple constrained guide trees and extracted the homology core shared across them. In simulations, these consensus alignments improved homology accuracy, site-pattern distributions, and recovery of correct splits. The same strategy was also applicable to real genomic data.

Consensus strategies inevitably reduce total alignment length. However, our results show that the reduction is controlled and that there is a trade-off between alignment length and precision. In multi-taxon simulations, a small number of replicate alignments already yields substantial accuracy gains while retaining a long alignment. More importantly, excluded regions are often unstable across guide trees and correspond to high homology uncertainty. Keeping them does not necessarily add reliable information and can instead introduce spurious support. For phylogenomic inference, retaining more aligned sequence is not always preferable if it distorts the site patterns distribution which is important for tree reconstruction. In this context, maximizing alignment coverage is not the primary objective. What matters more is whether the retained alignment is better suited for downstream tree reconstruction.

We also stress that a high-quality alignment is necessary but not sufficient to get the right tree and solve difficult phylogenetic problems. Even with a better input alignment, different tree inference frameworks may still disagree. Such disagreements can arise from model choices, different sensitivity to long-branch attraction, or genuine biological discordance, such as reticulate evolution. The value of consensus alignment lies in reducing guide-tree bias at the alignment stage. This makes the remaining discordance among tree inference more likely to arise from model differences or biological processes themselves, rather than from artefacts introduced during alignment.

Computational cost is a practical constraint for consensus strategies. Repeating whole-genome alignment becomes increasingly expensive as taxon sampling grows. Therefore, consensus methods require scalable workflow design. In our 43-taxon simulation, we addressed this with a hierarchical consensus strategy, showing that consensus alignment can be extended beyond small taxon sets while remaining computationally practical. This provides a practical route for applying similar strategies at larger scales. Under this hierarchical design, the additional runtime should depend primarily on how much of the tree can be reused and how much must be re-evaluated under multiple local guide trees. To give a concrete example, we considered the constraint tree reported for the 363-taxon Neoaves analysis of Stiller et al. (2024). If we extract the strict consensus tree following the procedure described on Page 84 of their Supplementary Information, 316 uncontroversial nodes are retained, whereas 46 nodes are left unresolved. These unresolved nodes correspond to 21 polytomies, including 16 three-way polytomies, with the root also counted as a three-way polytomy. Under our hierarchical reuse scheme, the 46 collapsed nodes determine the extra cost associated with multiple local guide-tree alignments. Using four guide trees by default for polytomies with more than three immediate child nodes, Formulae (1) and (2) in the Materials and Methods predict that our pipeline would require approximately 1.7 times as much CPU time as a single fully resolved guide-tree alignment. In practical terms, this suggests that an analysis requiring roughly three weeks under the original pipeline might remain feasible in about five weeks under our consensus-based strategy. In highly unresolved cases where little resolved structure can be reused, the analysis may take about four times as much CPU time.

While consensus alignments are more robust across multiple evaluation criteria in our analyses, the scope of the current implementation should be stated explicitly. The method is reference-centered because consensus columns are defined with respect to reference coordinates. It is therefore only suited for datasets with an appropriate reference genome. In addition, sampling multiple guide trees reduces dependence on any single guide tree but does not completely remove guide-tree effects; if some alternative placements or clusters are represented among the sampled trees, the stability of some columns may be overestimated. Finally, consensus extraction is conservative, so easier-to-align regions are more likely to be retained whereas difficult-to-align regions are more likely to be excluded. This selectivity may affect signal composition, although in our simulations the site-pattern distribution of the consensus alignment was closer to that of the true alignment than those of the individual guide-tree-specific alignments; downstream analyses should therefore remain alert to it.

Future improvements can proceed along four directions. First, beyond strict intersection, threshold-based consensus criteria can be explored. For example, one may retain columns or homology relationships consistent in at least a specified proportion of replicate alignments, providing a tunable trade-off between recall and robustness to guide-tree bias. This direction may be particularly useful for WGA applications in which alignment coverage is important, including orthology inference and comparative genomic analyses of conserved, constrained, or lineage-specific genomic features (Feng et al. 2020; Christmas et al. 2023). Future implementations could generate both strict and relaxed consensus outputs within the same alignment process: a strict consensus alignment aimed at phylogenetic tree reconstruction, and a relaxed consensus alignment designed to retain broader homology information for other WGA-based analyses. Second, guide-tree selection can be made more targeted. In addition to the constraints described in the main text, when competing phylogenetic hypotheses exist, guide trees supporting each hypothesis can be included to test which homology statements remain stable across hypotheses. Third, support for adding new taxa remains limited. Under the current framework, a new taxon cannot be incorporated directly into an existing consensus alignment without reintroducing guide-tree dependence. In practice, expanding the taxon set usually requires re-aligning both the original and new taxa under multiple guide trees and re-extracting the consensus, which is computationally expensive. A more efficient update strategy remains to be developed. Fourth, although consensus-based strategies have already been used for gene-or protein-scale multiple-sequence alignment, often by combining alignments generated with different aligners, substitution matrices, models, or parameter settings, the same principle could be applied more specifically to guide-tree uncertainty beyond Progressive Cactus and whole-genome alignment. For shorter gene-or locus-scale alignments, potentially unstable loci could first be prioritized from features such as low occupancy, high gap content, or weak locus-tree support. These loci could then be realigned under multiple alternative guide trees using progressive aligners that allow user-specified guide trees, and a locus-level consensus alignment could be extracted from the resulting alignments. This would differ from iterative refinement procedures that update guide trees internally during alignment optimization, because the alternative guide trees would be specified in advance and used as separate, predefined replicate conditions. This would apply the same logic used here: retaining homology statements that are stable to guide-tree choice in alignments where guide-tree dependence is likely to affect tree inference. Finally, we suggest that guide-tree sensitivity, long-branch attraction risk, and broader alignment robustness should be treated as routine aspects in evaluating WGA quality, so that reliability can be considered alongside scale and speed.

## Materials and Methods

### Hierarchical consensus-alignment workflow

#### Overall framework of the hierarchical consensus-alignment method

Our method takes a set of genomes, a rooted, partially resolved tree, and, by default, a reference genome as input. Before task construction, the input tree is rerooted at the pendant edge incident with the reference genome if the reference is not an immediate child of the root, so that all downstream alignment tasks are defined according to the reference-rooted tree. Resolved relationships in the rerooted tree are kept, and uncertain relationships are represented as polytomies. Instead of rerunning the entire dataset under multiple global guide trees, we address guide-tree uncertainty only within these local unresolved subproblems. The outputs are consensus whole-genome alignments including all leaf taxa in MAF and FASTA format, the inferred sequence of the root ancestor, and an alignment in HAL format including all leaf taxa and the inferred root ancestor. The general task-construction and execution workflow is summarized in Figure 6, and its application to the Neoaves simulation analysis is illustrated in Figure 4B.

**Figure 6.**
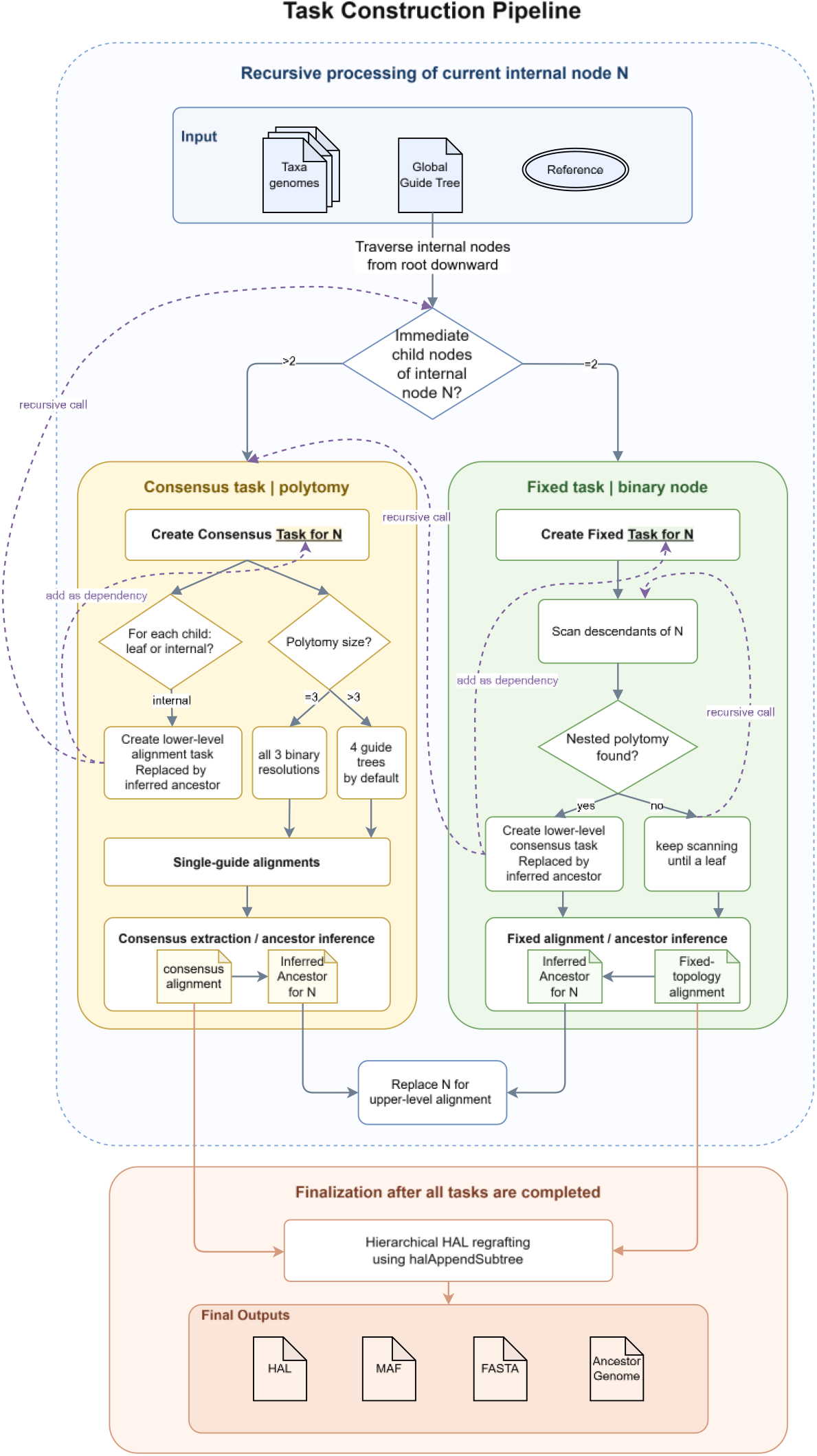
Recursive alignment task construction in the hierarchical consensus-alignment workflow.

#### Hierarchical decomposition of the input global guide tree into alignment tasks

The task-construction procedure is summarized in Figure 6 and its pseudocode is provided in Supplementary Note 1.1. The planner takes the reference-rooted tree and recursively traverses internal nodes and creates alignment tasks according to node resolution. At the root, when the non-reference child X is polytomous, the planner does not create a separate task for X; instead, the alignment of the reference with all children of X is treated as the top-level alignment task.

If a node is a polytomy, a consensus task is created. Any non-leaf descendant nodes immediately below the polytomy must first be converted into corresponding ancestor outputs, so these nodes are recursively created as lower-level tasks. These lower-level tasks may be either consensus tasks or fixed tasks, and they are added as dependencies of the current consensus task. If all immediate descendants are leaves, the consensus task has no lower-level dependencies. After all required lower-level tasks have been completed, the consensus task generates multiple alternative guide trees for the current polytomy. If it is a three-way polytomy, all three possible binary resolutions are generated. For larger polytomies, four alternative guide trees are generated by default. Each guide tree is then used for a separate Cactus alignment, and the consensus alignment is extracted by comparing the single guide-tree alignments, and the ancestor for the current task is inferred from this consensus alignment.

If a node is binary, it is treated as a fixed task. The planner then continues to recursively scan the descendants of this node to check whether any nested descendant polytomy is present. If no descendant polytomy is found, the fixed task has no lower-level dependencies and is aligned once using the fully resolved subtree under that node. In this case, the Cactus input tree is this fully resolved subtree, and all terminal labels are original taxa. If a descendant polytomy is found, that polytomy is first created as a lower-level consensus task and added as a dependency of the current fixed task. The fixed task is still aligned only once, but in its Cactus input tree, the descendant polytomy is replaced by its consensus-inferred ancestor.

Finally, the instruction commands are written in dependency order. Tasks without dependencies are placed in the first execution level, whereas tasks with dependencies are run only after their lower-level ancestor outputs have been generated. Thus, task generation and dependency recording proceed recursively from top to bottom, whereas command execution proceeds from lower-level to higher-level tasks according to the dependencies. After all tasks have been completed, lower-level HAL alignments are regrafted hierarchically into the final root-level alignment using halAppendSubtree from the HAL package (Hickey et al. 2013). The final HAL alignment is then exported to MAF and FASTA formats for downstream analyses.

#### Guide-tree design for consensus alignment task

In the default implementation, we use all three rooted triples for polytomies with three children and four fully resolved binary guide trees for polytomies with at least four children. Candidate guide trees are generated by randomly sampling from either the uniform or the Yule model distribution. A sampled tree is added to the list of accepted trees if its RF-distance and triplet distance from all previously accepted trees is above pre-defined threshold (default 1 for the RF distance and 2/3 for the triplet distance). In practice, under these default strict thresholds, the tree generator may not always find the requested number of mutually distant guide trees. In such cases, the program returns the largest valid set found under the current constraints. Users may modify the requested number of guide trees, the distance thresholds, the model, and the maximum number of sampled trees. By default, these settings are applied globally across the hierarchical alignment workflow. Users who wish to use different guide-tree-generation settings for particular levels or subproblems can instead execute the corresponding code and commands separately for those steps.

For each consensus task, consensus columns are extracted relative to the reference genome. In Figure 7, Anc denotes the current polytomy being processed, ref denotes the reference genome, and A, B, and C denote the child genomes in the local guide tree. A, B, and C may each be either a taxon genome or an ancestral genome inferred for a cluster. For a given consensus task, depending on the reference position, (1) if the global reference is one of the immediate child nodes of the root and root is a polytomy, the reference participates directly in guide-tree generation, so its position can vary among the sampled guide trees (2) if the root has an exactly one child X besides the reference, and X is a polytomy, the reference is treated as external to each polytomy during local guide-tree generation, local candidate guide trees are generated by resolving the child nodes of Anc, while ref is placed outside the resolved ingroup subtree as a fixed external reference.

**Figure 7.**
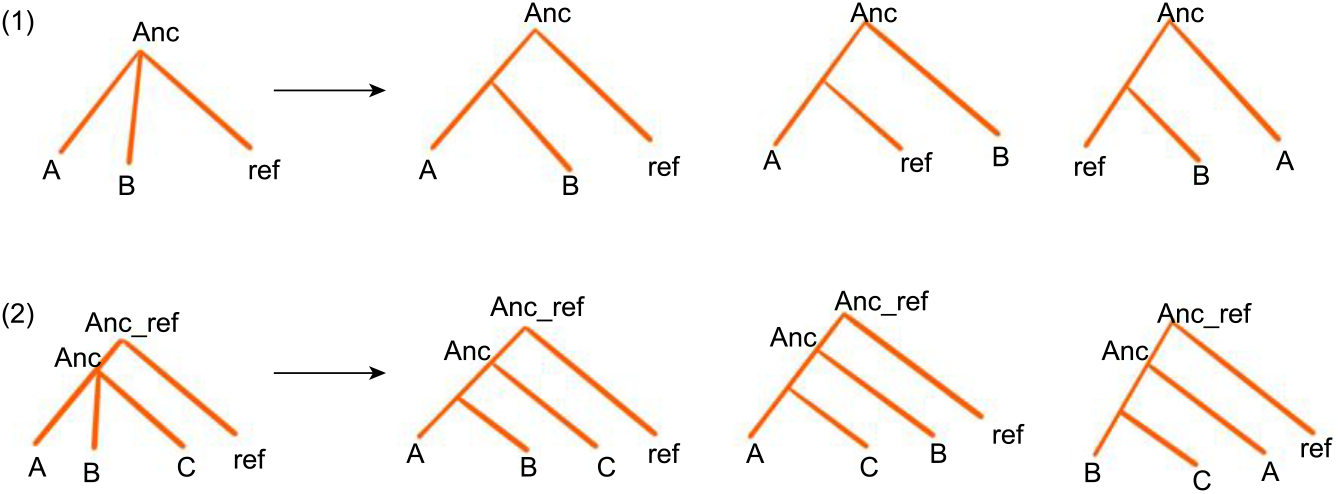
Reference placement in local guide-tree generation under two possible relationships between the reference and the current polytomy.

#### Consensus column extraction

For each polytomy, we construct a consensus alignment by rerunning Progressive Cactus under multiple candidate guide trees and retaining only stable homology across runs. For the same set of genomes, we use a set of candidate guide trees T={*T*_1_,…,*T_m_*} and run Progressive Cactus with identical non-tree parameters to obtain *m* HAL files.

Each HAL is converted to a RefSpecies-referenced MAF using cactus-hal2maf --outType single --chunkSize 500000 --refGenome RefSpecies –noAncestors. For cactus-hal2maf outputs, the reference row is fixed by --refGenome, duplicates are collapsed to at most one record per taxon per block by -- outType single, and blocks are ordered by reference chromosomes and their coordinates. If MAFs are obtained from other sources, equivalent normalization (duplicate removal, fixing the reference as the first row, and sorting by reference chromosomes and coordinates) is required before consensus extraction.

To scale to genome-wide alignments, we use a two-level partitioning and parallelization strategy (Figure 8A). We first split the genome-wide MAF into chromosome-level MAFs by the reference genome; for long chromosomes, we further partition each chromosome MAF into interval-based subfiles (Algorithm 1 in Supplementary Note 1.2) to reduce per-task memory need and improve throughput. Within each subfile, we divide its reference-coordinate range across multiple workers to perform column-wise comparison and block construction in parallel (Algorithm 2 in Supplementary Note 1.2). Each worker outputs blocks for its assigned interval; blocks are concatenated in reference-coordinate order without connecting across worker or subfile boundaries, ensuring task independence and deterministic aggregation.

**Figure 8.**
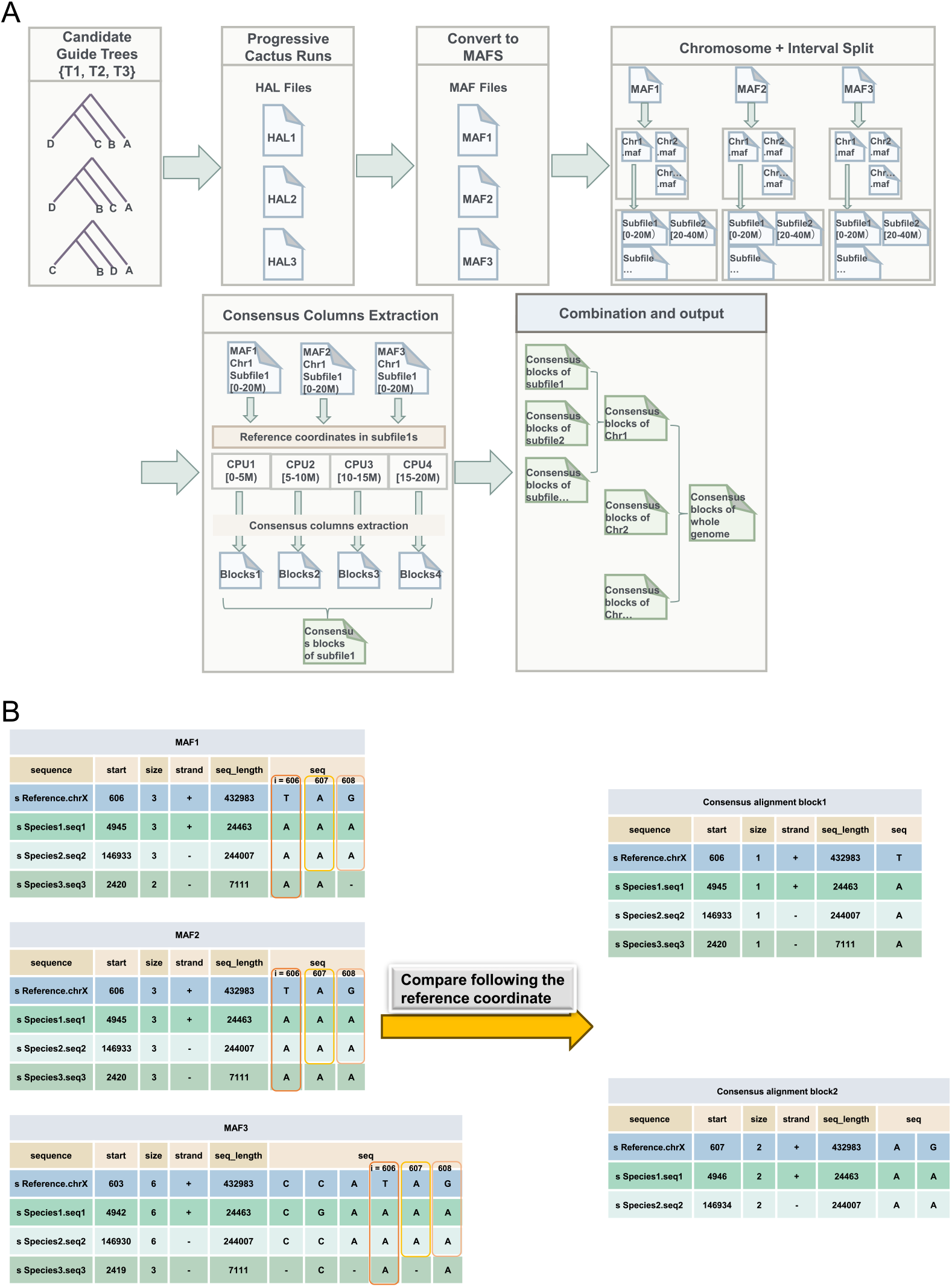
Consensus alignment extraction procedure. **A**, Partitioning genome-wide MAF alignments into chromosome-level and interval-based subfiles based on reference coordinates, followed by parallel processing of subfiles for scalable consensus extraction. **B**, Column-wise consensus extraction in which homology assignments are compared across multiple guide-tree-based alignments and only consistent taxon assignments are retained.

Consensus is extracted in a column-wise manner, with each column indexed by the reference coordinate i; the decision rule is illustrated in Figure 8B. For each i, we retrieve the corresponding column from all guide-tree MAFs and evaluate, for each non-reference taxon, whether its homology assignment is consistent across inputs. We represent the assignment at i as (seqName, strand, coord, base) and retain a taxon only if this information is identical across all input MAFs; otherwise the taxon is treated as missing (gap) at that column. If no non-reference taxa are retained at a coordinate, the column is discarded. Retained columns are assembled into MAF blocks in reference-coordinate order: two adjacent reference columns are concatenated only if they retain the same set of records, and each taxon preserves the same seqName and strand within the block and satisfies the MAF start/size coordinate progression (coord=start+size, where size is the cumulative non-gap count); otherwise the current block is terminated and a new block is started.

#### Ancestor inference and HAL construction

After a consensus alignment has been extracted for a consensus task, the current polytomy ancestor is inferred and a corresponding HAL alignment is generated. The details are provided in Supplementary Note 1.3 and illustrated in Supplementary Figures 1, 2.

The reference lies outside the current polytomy after rerooting the input global guide tree. The task-level HAL output represents the relationship hierarchically, with an embedded tree such as (reference,(A,B,C,D)ingroup_ancestor)top_ancestor. When this HAL is used as a lower-level task HAL output for regrafting into an upper-level HAL, ingroup_ancestor is used as the subtree root for halAppendSubtree.

#### Approximation of computational cost

To approximate how many times as much CPU time our hierarchical consensus pipeline requires relative to a single fully resolved alignment, we use the following node-count-based formula. We counted the number of Cactus alignment runs required under the default guide-tree-generation strategy. Let *n* denote the number of taxa. A fully resolved rooted binary tree on the same taxon set contains *n*−1 internal nodes, corresponding to *n*−1 alignment runs in a standard fully resolved progressive alignment.

Let *m* be the number of polytomous clusters, *p* be the number of interior vertices in the global guide tree, and *x* be the number of 3-way polytomies. Under the default setting, each three-way polytomy is resolved using all three possible binary guide trees, whereas each polytomy with more than three immediate child nodes is resolved using four randomly generated guide trees. For each consensus task, the reference is added as a fixed external reference to each candidate guide tree. This increases each guide-tree-specific alignment by one additional alignment run. Therefore, the additional cost is 3*x* runs for three-way polytomies and 4(*m* − *x*) runs for polytomies with more than three immediate child nodes. Before adding this external-reference cost, a *k*-way polytomy contributes *k* − 1 alignment rounds. The sum of all out-degrees minus one over all interior vertices equals *n* − 1 for every rooted phylogenetic tree. The bifurcation vertices contribute *p* − *m* to the sum, and trifurcations contribute 2*x*. Therefore, polytomies with at least four children contribute *n* − 1 − (*p* − *m*) − 2*x*.

We denote the total number of required alignment runs as *N_align_* and this number can be written as

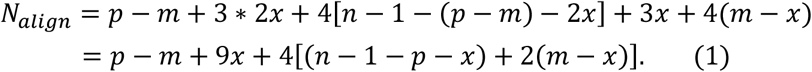

Depending on whether the reference-rooted tree is obtained by rerooting, and on whether the root is binary or polytomous, up to four root-level alignment runs can be saved. The CPU-time ratio is then approximated as

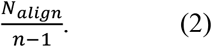

The worst case is a guide tree where all interior vertices have out-degree four. Then the running time increases by a factor of 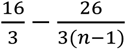, thus 5.34 is an upper bound.

### Method details

A detailed description of all simulations and real data experiments can be found in Supplementary Material Notes 2 and 3.

## Data availability

The genomes for the real data experiments were taken from Jarvis et al. (2014), see Subsection 3.1 in Supplementary Material for details.

All alignments can be downloaded from the EMBL-EBI repository BioStudies under accession S-BSST3394.

The implementation of the consensus-alignment framework, including the hierarchical alignment workflow and consensus extraction modules, is available at GitHub (https://github.com/taoqxtaozi/Common_Col).

## Notes

https://www.ebi.ac.uk/biostudies/studies/S-BSST3394

### Competing Interest Statement

The authors have declared no competing interest.

### Summary of Updates

We changed our manuscript based on the remaining points raised by the referees and editor during the publication process.

