## Supplementary Material for "Guide-tree bias of whole genome alignment can mislead phylogenomic analyses"

The following supplementary notes, figures, and tables provide pseudocode for the consensus algorithm and additional methodological details for the simulation and empirical analyses.

Supplementary Note 1 provides pseudocode for hierarchical alignment tasks construction and the consensus column extraction procedures described in the Methods. Section 1.1 includes the main recursive planner and three subroutines. Section 1.2 describes the consensus-column extraction workflow in two steps: Algorithm 1 describes how reference-coordinate intervals were generated and how input alignments were split by interval. Algorithm 2 describes how consensus columns were extracted across input alignments. Section 1.3 describes ancestor inference from consensus alignment.

Supplementary Note 2 provides details of the simulation experiments involving 4, 12, and 43 taxa. It first describes procedures shared across the three simulation settings, including true-tree construction, extraction of simulated genomes and true alignments, Progressive Cactus alignment, format conversion, tree inference, and alignment-precision calculation. It then provides experiment-specific details for the 4-taxon simulations, including site-pattern classification and Jensen–Shannon (JS) divergence calculations, and for the 12- and 43-taxon simulations, including true-tree split recovery and hierarchical consensus pipeline settings.

Supplementary Note 3 provides details of the empirical analysis involving eight taxa. It describes the data sources, reference genome choice, constraints used to define guide trees for Progressive Cactus, conversion of HAL outputs to MAF format, generation of FASTA alignments, tree inference from concatenated alignments using IQ-TREE, CASTER-site, maximum parsimony in PAUP*, and the loci-ASTRAL supertree pipeline. This section also includes the comparison between current and our previous study based on chicken chromosome 1.

**Figures**

| Section | Figure | Description |
|---|---|---|
| Methods | Figure S1 | Infer two ancestors from consensus alignment |
| | Figure S2 | Infer one ancestor from consensus alignment |
| Guide Tree Bias of Progressive Cactus for Bird Data | Figure S3 | Average quartet support across 1,000 consecutive loci for the first 15 and Z chicken chromosomes |
| | Figure S4 | Guide trees and result trees of the 4-taxon set chicken, seriema, zebra finch, and turkey vulture. |
| 4-taxon simulation | Figure S5 | Guide trees used in four-taxon simulation. |
| | Figure S6 | Boxplot of recall of Realistic tree scenario. |
| | Figure S7 | Boxplot of JS-divergence of Realistic scenario. |
| | Figure S8 | Boxplot of precision of Felsenstein tree scenario. |
| | Figure S9 | Boxplot of recall of Felsenstein tree scenario. |
| | Figure S10 | Boxplots of JS-divergence of Felsenstein tree scenario. |
| | Figure S11 | Boxplot of precision of Farris tree scenario. |
| | Figure S12 | Boxplot of recall of Farris tree scenario. |
| | Figure S13 | Boxplot of JS-divergence of Farris tree scenario. |
| | Figure S14 | Boxplot of precision of Clock-like tree scenario. |
| | Figure S15 | Boxplot of recall of Clock-like tree scenario. |
| | Figure S16 | Boxplots of JS-divergence of Clock-like tree scenario. |
| 12-taxon simulation | Figure S17 | Guide trees used in 12-taxon simulation. |
| | Figure S18 | Average number of nucleotides of all alignments including filtered ones. |
| | Figure S19 | Precision of all alignments including filtered ones. |
| | Figure S20 | Recall of all alignments including filtered ones. |
| | Figure S21 | Split reconstruction results of all alignments including filtered ones. |
| | Figure S22 | Fraction of erroneous homology pairs in the simulated 12-taxon dataset. |
| 43-taxon simulation | Figure S23 | Simulated species tree (drawn to scale). |
| | Figure S24 | Global guide tree used to define polytomies. |
| | Figures S25-S29 | Local guide trees used in hierarchical consensus pipeline. |
| | Figure S30 | Globally resolved guide trees used in single-guide tree alignments. |
| | Figure S31 | Average number of nucleotides of all alignments. |
| | Figure S32 | Recall of all alignments (using chicken as reference). |
| | Figure S33 | Precision of all alignments (using chicken as reference). |

| | | |
|---|---|---|
| | Figure S34 | Full split reconstruction results (using chicken as reference). |
| | Figure S35 | Recall of all alignments (using zebra finch as reference). |
| | Figure S36 | Alignments precision (using zebra finch as reference). |
| | Figure S37 | Full split reconstruction results (using zebra finch as reference). |
| Real 8-taxon experiment | Figure S38 | Result trees inferred by IQ-TREE partition (10k). |
| | Figure S39 | Result trees inferred by CASTER-site. |
| | Figure S40 | Result trees inferred by maximum parsimony (PAUP*). |
| | Figure S41 | Result trees inferred by loci-ASTRAL super tree method |

**Tables**

| Analysis section | Table | Description |
|---|---|---|
| Guide Tree Bias of Progressive Cactus for Bird Data + Real 8-taxon experiment | Table S1 | Summary information of genome assemblies |
| Guide Tree Bias of Progressive Cactus for Bird Data | Tables S2, S3 | Log-likelihood differences and optimized interior branch lengths of two real bird 4-taxon datasets, respectively. |
| Real 8-taxon experiment | Table S4 | Computational cost and alignment statistics |

**Note 1: Details of algorithms**

**1.1 Pseudocode of hierarchical alignment tasks construction**

**Main procedure: Construct hierarchical alignment tasks**

Input:

T: rooted partially resolved guide tree, with Root as the root name

R: reference genome

genome_paths: paths to leaf genomes

Output:

final_HAL, final_MAF, final_FASTA, ancestor_FASTA

Definitions:

Task(N): an alignment task created for internal node N. A task is either a fixed-topology task or a consensus task. Each task records its input child nodes, lower-level dependencies, Cactus input files, task-level ancestor FASTA, and task-level HAL output.

AllTasks: the set of all tasks created from the input tree.

DependencyRecords: records specifying lower-level tasks that must be completed before an upper-level task can be executed.

```
Parse Newick tree T
AllTasks ← ∅
DRs ← ∅ # short for DependencyRecords
Levels ← ∅
If T has the top-level form (R, X)Root and X is a polytomy then
    ExpandTask(root(T), children(X), external_reference = R, AllTasks, DRs)
Else
    ExpandTask(root(T), children(root(T)), external_reference = None, AllTasks, DRs)
End if
UnassignedTasks ← AllTasks
CT ← ∅ # short for CompletedTasks
While UnassignedTasks ≠ ∅ do
    L ← ∅
    For each task t in UnassignedTasks do
        If t has no dependencies in DRs, or all dependencies of t in DRs are in CT then
            Add t to L
        End if
    End for
    Append L to the end of Levels
    UnassignedTasks ← UnassignedTasks − L
    CompletedTasks ← CompletedTasks ∪ L
End while
For each execution level L in Levels, from first to last, do
    For each task t in L do
        If t is a fixed-topology task then
```

Run Cactus alignment once
**Else if** t is a consensus task **then**
**For** each local guide tree g of t **do**
Run Cactus alignment under g
**End for**
Extract the consensus alignment
**End if**
Generate one task-level HAL output
Infer the task-level ancestor FASTA
**End for**
**End for**
final_HAL ← task-level HAL output of the root-level task
**For** each lower-level task t in regrafting order **do**
halAppendSubtree final_HAL task_HAL(t) t.name t.name --merge
**End for**
Convert final_HAL to MAF
Convert final_MAF to FASTA
**Return** final_HAL, final_MAF, final_FASTA, ancestor_FASTA

---

**Subroutine1: ExpandTask(N, child_nodes, external_reference, AllTasks, DRs)**
Input:
N: an internal node to be converted into an alignment task
child_nodes: child nodes used to classify N as a fixed-topology or consensus task

**If** number of child_nodes > 2 **then**
Create consensus task Task(N)
Add Task(N) to AllTasks
GenerateGuideTrees(Task(N), child_nodes, external_reference)
**For** each internal child c in child_nodes **do**
ExpandTask (c, children(c), external_reference = None, AllTasks, DRs)
Add Task(c) as a dependency of Task(N) to DRs
**End for**
**Else if** number of child_nodes = 2 **then**
Create fixed-topology task Task(N)
Add Task(N) to AllTasks
**For** each child c in child_nodes **do**
ScanFixedSubtree(c, Task(N), AllTasks, DRs)
**End for**
**End if**

---

**Subroutine 2: GenerateGuideTrees(Task(N), child_nodes, external_reference)**

```
# Consider the role of reference R in the current consensus task
external_outgroup ← None
If R is an immediate child of the current polytomy then
    sampled_nodes ← child_nodes
Else
    sampled_nodes ← child_nodes
    external_outgroup ← R
End if
If number of sampled_nodes = 3 then
    guide_trees ← all 3 binary resolutions of sampled_nodes
Else
    guide_trees ← 4 local guide trees of sampled_nodes by default
End if
If external_outgroup is not None then
    place external_outgroup as the outgroup of each guide tree
End if
Return guide_trees
```

**Subroutine 3: ScanFixedSubtree(v, parent_task, AllTasks, DRs)**

```
If v is a leaf then
    Stop scanning this branch
Else if v is a polytomy then
    ExpandTask(v, children(v), external_reference = None, AllTasks, DRs)
    Add Task(v) as a dependency of parent_task to DRs
    Replace v by the ancestor label of Task(v) in the fixed-task input tree
    Stop scanning this branch
Else if v is a binary internal node then
    For each child u of v do
        ScanFixedSubtree(u, parent_task, AllTasks, DRs)
    End for
End if
```

#### 1.2 Pseudocode of consensus column extraction

**Algorithm 1: Generate intervals and Separate Alignments based on intervals**

**Input:**

start coordinate
end coordinate
one alignment
num = number of files the user want to separate into

**Output:**

a list of intervals
a list of separated alignment based on intervals

```
intervals ← ∅
seperated_alignment_list ← ∅
step = int((end - start)/(num + 1))
interval_lst ←[interval([start, start+step))..., interval([start+num*step,end))]
For interval in interval_lst do
    tmp_list ← ∅
    For block in alignment do
        If coordinates of block overlaps interval then
            tmp_list ← tmp_list ∪ block
        End if
    seperated_alignment_list ← seperated_alignment_list ∪ tmp_list
    End for
End for
Return seperated_alignment_list
```

**Algorithm 2: Extract Consensus Columns**

**Input**:

lists: Lists of all input alignments

interval_used: Interval of coordinates to compare

**Output**:

blocks: List of consensus sequence blocks

```
blocks ← ∅
con_lst ← ∅ (to store consensus sequences)
For each position i in the range of interval_used do
    list_info ← ∅
    For list in lists do
        Add sequences information of aln_column corresponding to i to list_info
    End for
    Identify common_elements across list_info
    If common_elements are found then
        temp_con_lst ← common_elements
        If con_lst = ∅ then
            con_lst ← temp_con_lst
        Else
            If temp_con_lst is contiguous with con_lst then
                Concatenate temp_con_lst to con_lst
            Else
                blocks ← blocks ∪ con_lst
                con_lst ← temp_con_lst
            End if
        End if
    Else
        If con_lst not ∅ then
            blocks ← blocks ∪ con_lst
            con_lst ← ∅
        End if
    End if
End For
If con_lst ≠ ∅ then
    Add con_lst to blocks
End if
Return: blocks
```

#### 1.3 Infer Ancestor and construct HAL from consensus alignments

This function is implemented in the consensus-extraction pipeline provided with our program. After a consensus alignment has been extracted in MAF format, the pipeline constructs ancestor-containing HAL alignments under two ancestor-inference modes, depending on the placement of the reference relative to the current polytomy (Figures S1, S2). The resulting HAL alignment is used for hierarchical regrafting, and can also be used directly when the consensus-extraction workflow is run independently.

The first mode is the external-reference mode, which is used in the reference-rooted hierarchical workflow described in Methods (Figure S1). In this mode, the reference is placed outside the current polytomy, and the local consensus task has the form (ref,(A,B,C)A1)A2. The pipeline therefore infers two ancestor genomes: the ingroup ancestor A1, inferred from the non-reference child nodes of the current polytomy, and the upper ancestor A2, inferred from the reference and the inferred ingroup ancestor A1.

The second mode is the reference-included mode, which applies when the reference is one of the immediate child nodes of the current polytomy, giving a local consensus task of the form (ref,A,B,C)A1. In this case, only one ancestor genome, A1, is inferred (Figure S2).

Both modes use the same chromosome-level ancestor-row addition step, hereafter referred to as add_Anc_process. For each chromosome-specific consensus MAF, the pipeline adds an ancestor row named as <ancestor>.<chromosome>. In aligned consensus blocks, this ancestor row is initially filled with N characters. To keep the inferred ancestor genome in the same chromosome-coordinate system as the reference, intervals between adjacent consensus blocks are filled using the corresponding sequence from the reference chromosome. Therefore, the ancestor chromosome has the same coordinate span as the reference chromosome, while positions supported by consensus alignment blocks are later updated by maximum-likelihood ancestral reconstruction.

The pipeline uses a base JC69 model file, init.mod, which contains the substitution model parameters but leaves the TREE line to be filled according to the taxa used in each ancestor-inference step. For each ancestor level, the pipeline copies init.mod to a task-specific model file and appends the corresponding TREE: line. In this TREE line, zero-length internal branches are used to represent the unresolved star-like relationship, whereas terminal branches are initialized with length 1. The value 1 in the input TREE line serves only as an initial branch-length value, whereas the branch lengths used by ancestorsML are those estimated by phyloFit from the merged MAF.

The base init.mod file is:

```
ALPHABET: A C G T
ORDER: 0
SUBST_MOD: JC69
BACKGROUND: 0.250000 0.250000 0.250000 0.250000
RATE_MAT:
-1.000000    0.333333    0.333333    0.333333
0.333333    -1.000000    0.333333    0.333333
0.333333    0.333333    -1.000000    0.333333
0.333333    0.333333    0.333333    -1.000000
```

TREE:

**External-reference mode: two-ancestor inference**
In the external-reference mode, illustrated in Figure S1, the reference is placed outside the current polytomy, as in (ref,(A,B,C)A1)A2. This is the mode used by the reference-rooted hierarchical workflow described in the main method. The first step infers the ingroup ancestor A1 from the non-reference child nodes. For each chromosome, the pipeline applies add_Anc_process to the chromosome-level consensus MAF, adding an A1.<chromosome> row and producing consensus_A1.maf. The reference row is then removed, producing consensus_A1_noref.maf. These chromosome-level consensus_A1_noref.maf files are merged into one task-level MAF, called whole_consensus_A1_noref.maf.

The model tree for the ingroup ancestor includes only the non-reference child nodes of the current polytomy. For example:
TREE: (A:1,(B:1,C:1):0!);

The TREE line in init.mod is updated separately and this file is called consensus_A1_tryMLstartree_useThis.mod. The fitted model consensus_A1_JC69modle.mod for A1 is obtained from the merged reference-excluded MAF:
phyloFit --init-model consensus_A1_tryMLstartree_useThis.mod
--subst-mod JC69
--out-root consensus_A1_JC69modle
--msa-format MAF whole_consensus_A1_noref.maf

Then, for each chromosome, the chromosome-level reference-excluded MAF is converted to HAL with A1 as the HAL reference genome:
maf2hal --refGenome A1 consensus_A1_noref.maf consensus_A1_noref.hal

The fitted model is used by ancestorsML to infer nucleotide states for A1.<chromosome>:
ancestorsML --printWrites consensus_A1_noref.hal A1 consensus_A1_JC69modle.mod \
> consensus_A1_JC69modle.tsv

The inferred nucleotides are written back into the HAL alignment:
halWriteNucleotides consensus_A1_noref.hal consensus_A1_JC69modle.tsv

The inferred ingroup ancestor chromosome is extracted in FASTA format:
hal2fasta consensus_A1_noref.hal A1 > A1.<chromosome>.fa

The inferred A1.<chromosome> sequence is then written back into the corresponding chromosome-level consensus_A1.maf, replacing the original N characters in the A1.<chromosome> row and producing consensus_A1_new.maf. The same inferred sequence is also written back into consensus_A1_noref.maf, producing consensus_A1_noref_new.maf.

The chromosome-level A1.<chromosome>.fa files are concatenated to generate the full inferred ingroup ancestor genome:

cat A1.<chromosome1>.fa A1.<chromosome2>.fa ... > A1.fa

After all chromosomes have been processed, the chromosome-level consensus_A1_noref_new.maf files are merged into one MAF, generating whole_consensus_A1_noref_new.maf, and converted into the ingroup HAL alignment:
maf2hal --refGenome A1 whole_consensus_A1_noref_new.maf whole_consensus_A1_noref_new.hal

For the second step, the upper ancestor A2 is inferred from the reference and the inferred ingroup ancestor A1. For each chromosome, the original ingroup child genomes are removed from consensus_A1_new.maf, leaving a reduced MAF containing the reference and the inferred A1 sequence. This reduced MAF is called consensus_A1_new_noIngroup.maf. An A2.<chromosome> row filled with N characters is then added to this MAF, producing consensus_A1_new_noIngroup_A2.maf. These chromosome-level consensus_A1_new_noIngroup_A2.maf files are merged into one task-level MAF, called whole_consensus_A1_new_noIngroup_A2.maf.

The model tree for the upper ancestor is:
TREE: (ref:1,A1:1);

The TREE line in init.mod is updated separately and this file is called
consensus_A1_new_noIngroup_A2_tryMLstartree_useThis.mod. The fitted model
consensus_A1_new_noIngroup_A2_JC69modle.mod for A2 is obtained from the merged MAF:

phyloFit --init-model consensus_A1_new_noIngroup_A2_tryMLstartree_useThis.mod
--subst-mod JC69
--out-root consensus_A1_new_noIngroup_A2_JC69modle
--msa-format MAF whole_consensus_A1_new_noIngroup_A2.maf

Then, for each chromosome, the chromosome-level MAF is converted to HAL with A2 as the HAL reference genome:
maf2hal --refGenome A2 consensus_A1_new_noIngroup_A2.maf consensus_A1_new_noIngroup_A2.hal

The fitted model is used by ancestorsML to infer nucleotide states for A2.<chromosome>:
ancestorsML --printWrites consensus_A1_new_noIngroup_A2.hal A2 \
consensus_A1_new_noIngroup_A2_JC69modle.mod \
> consensus_A1_new_noIngroup_A2_JC69modle.tsv

The inferred nucleotides are written back into the HAL alignment:
halWriteNucleotides consensus_A1_new_noIngroup_A2.hal \
consensus_A1_new_noIngroup_A2_JC69modle.tsv

The inferred upper-ancestor chromosome is extracted in FASTA format:
hal2fasta consensus_A1_new_noIngroup_A2.hal A2 > A2.<chromosome>.fa

The inferred A2.<chromosome> sequence is then written back into the corresponding chromosome-level consensus_A1_new_noIngroup_A2.maf, replacing the original N characters in the A2.<chromosome> row and producing consensus_A1_new_noIngroup_A2_new.maf. After all chromosomes have been processed, the chromosome-level consensus_A1_new_noIngroup_A2_new.maf files are merged into one MAF, called whole_consensus_A1_new_noIngroup_A2_new.maf, and converted into the upper HAL alignment:

```
maf2hal --refGenome A2 whole_consensus_A1_new_noIngroup_A2_new.maf \
whole_consensus_A1_new_noIngroup_A2_new.hal
```

The chromosome-level A2.<chromosome>.fa files are concatenated to generate the full inferred upper ancestor genome:

```
cat A2.<chromosome1>.fa A2.<chromosome2>.fa ... > A2.fa
```

Finally, the ingroup HAL is regrafted into the upper HAL at the shared A1 node:

```
halAppendSubtree whole_consensus_A1_new_noIngroup_A2_new.hal \
whole_consensus_A1_noref_new.hal A1 A1 --merge
```

The resulting HAL contains both ancestor levels:

(ref,(A,B,C)A1)A2;

In the hierarchical workflow, lower-level consensus HAL alignments are regrafted onto the corresponding higher-level HAL alignment using the shared ancestor node as the merge point. For the example above, A1 is used as the merge point:

```
halAppendSubtree top-level.hal lower-level.hal A1 A1 --merge
```

This operation regrafts only the subtree descending from A1 in the lower-level HAL into the corresponding A1 node of the higher-level alignment.

**Reference-included mode: one-ancestor inference**

If the reference for the current consensus-extraction task is not an outgroup of the current polytomy, for example (A,B,C,ref)A1, only one ancestor genome, A1, is inferred (illustrated in Figure S2). For each chromosome, the pipeline applies add_Anc_process to the chromosome-level consensus MAF, adding an A1.<chromosome> row and producing consensus_A1.maf. These chromosome-level consensus_A1.maf files are then merged into one task-level MAF, called whole_consensus_A1.maf.

The model tree includes the reference and the other immediate child nodes of the current polytomy. For example, the appended tree will be:

TREE: (ref:1,(A:1,(B:1,C:1):0!):0!);

The TREE line in init.mod is updated separately and this file is called consensus_A1_tryMLstartree_useThis.mod. The fitted model consensus_A1_JC69modle.mod is obtained from the merged ancestor-augmented MAF:

```
phyloFit --init-model consensus_A1_tryMLstartree_useThis.mod --subst-mod JC69 \
```

--out-root consensus_A1_JC69modle --msa-format MAF whole_consensus_A1.maf

Then, for each chromosome, the chromosome-level ancestor-augmented MAF is converted to HAL with A1 as the HAL reference genome:
maf2hal --refGenome A1 consensus_A1.maf consensus_A1.hal

The fitted model is used by ancestorsML to infer nucleotide states for A1.<chromosome>:
ancestorsML --printWrites consensus_A1.hal A1 consensus_A1_JC69modle.mod \
> consensus_A1_JC69modle.tsv

The inferred nucleotides are written back into the HAL alignment:
halWriteNucleotides consensus_A1.hal consensus_A1_JC69modle.tsv

The inferred ancestor chromosome is extracted in FASTA format:
hal2fasta consensus_A1.hal A1 > A1.<chromosome>.fa

The inferred A1.<chromosome> sequence is then written back into the corresponding chromosome-level consensus_A1.maf, replacing the original N characters in the A1.<chromosome> row and producing consensus_A1_new.maf. After all chromosomes have been processed, the chromosome-level consensus_A1_new.maf files are merged into one MAF, called whole_consensus_A1_new.maf. This MAF can be converted to the final ancestor-containing HAL alignment for this task:
maf2hal --refGenome A1 whole_consensus_A1_new.maf consensus.hal

The chromosome-level ancestor FASTA files are concatenated to generate the full inferred ancestor genome:
cat A1.<chromosome1>.fa A1.<chromosome2>.fa ... > A1.fa

The final outputs of a consensus task include the consensus MAF, the consensus FASTA, the HAL alignment, and the inferred ancestor genome or genomes. Here, maf2hal, ancestorsML, halWriteNucleotides, halAppendSubtree, and hal2fasta are from the HAL package, and phyloFit is from the PHAST package (Hubisz et al. 2011).

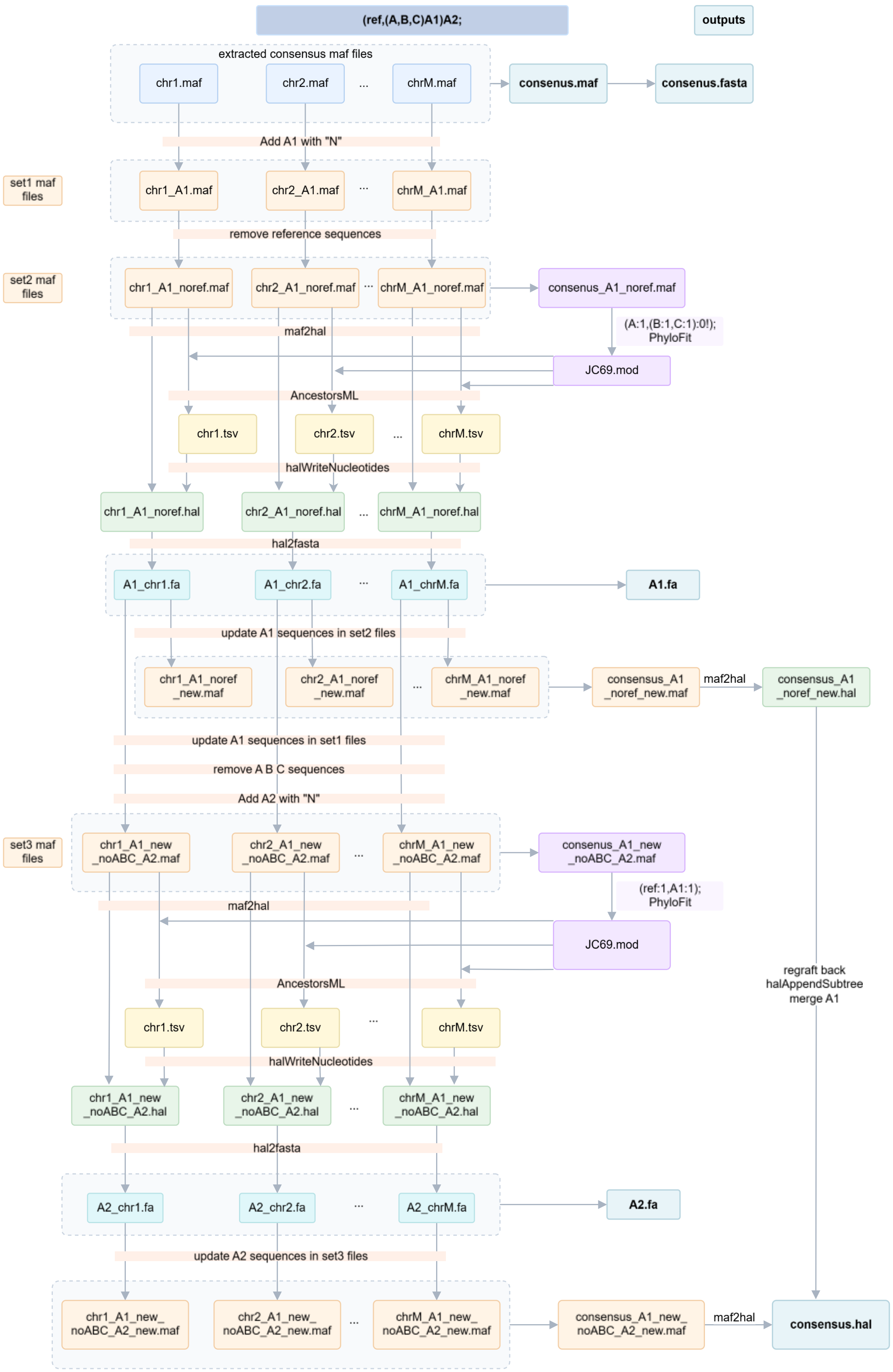


**Figure S1** Ancestor inference and HAL construction in the external-reference mode.

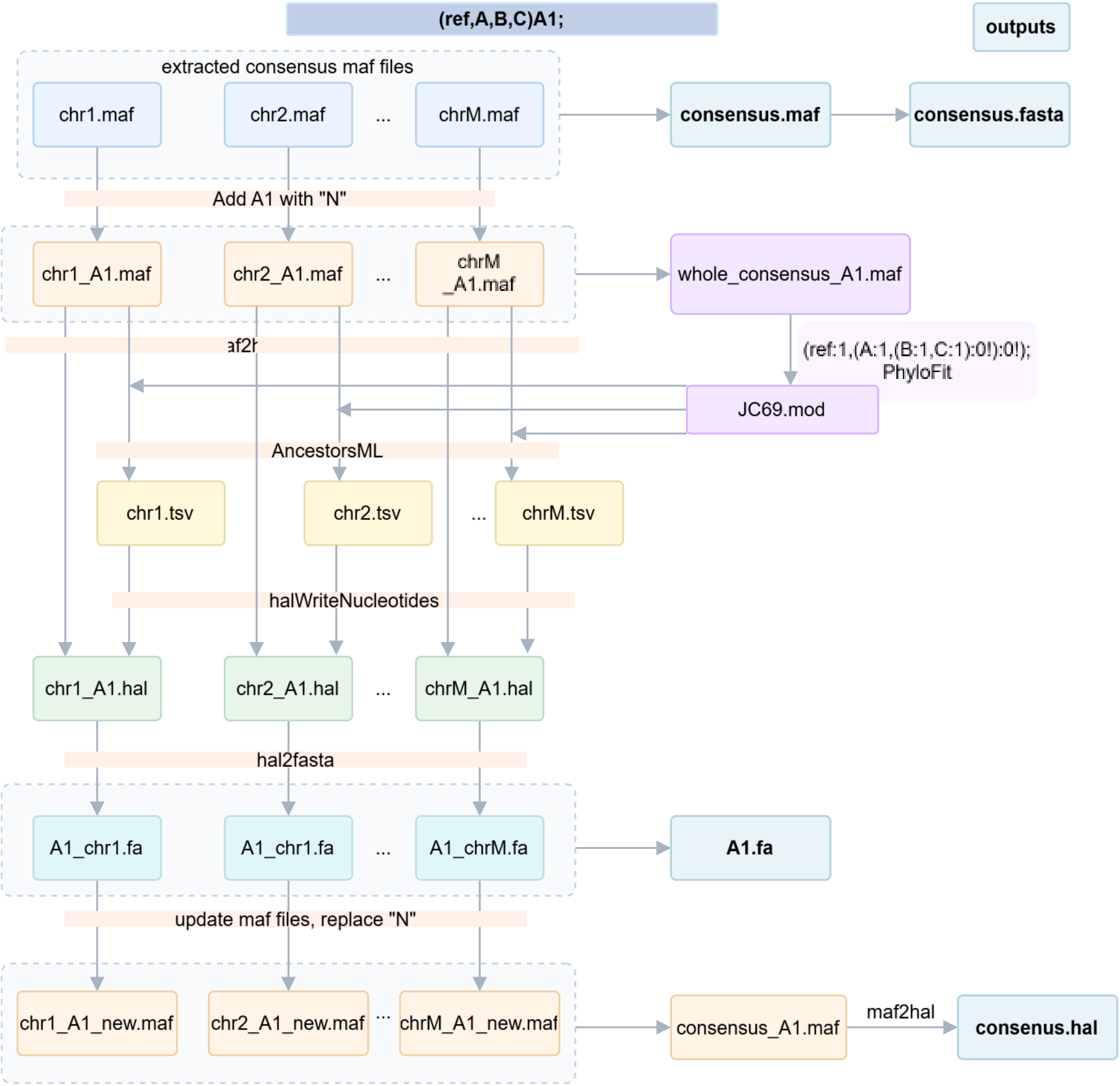


**Figure S2** Ancestor inference and HAL construction in the reference-included mode.

**Note 2: Details of simulation experiments**

**2.1 Common process shared by simulation experiments**

**1) Build evolutionary true tree we used**

Step 1. Extraction subtree from Jarvis TENT tree

We performed simulations on multiple taxon sets with varying sizes, for example 4, 12, and 43 taxa. We pruned a subtree from the Jarvis TENT backbone tree, used as evolutionary true tree.

We used nw_prune from Newick Utilities v1.6. Here ${taxon set} denotes the taxon list for the current simulation.

nw_prune -v TENT.ExaML.tre ${taxon set} > subtree.tre

Step 2. Branch-length rescaling and TrueTree construction

Evolver did not reliably finish under the original branch lengths. We therefore uniformly rescaled all branch lengths in subtree.tre by a factor of 0.3, applied to both pendant and internal edges, to generate TrueTree.tre.

**2) Evolver simulations and replicate settings**

We ran Evolver simulations for each taxon set with 100 replicates. Replicate i used seed i for i from 1 to 100. The root/ and params/ directories were taken from the resource package provided in the Progressive Cactus study (http://courtyard.gi.ucsc.edu/~jcarmstr/datastore/progressiveCactusEvolverSim.tar.gz, and it is archived at the Internet Archive https://web.archive.org/web/20220501000000*/http://courtyard.gi.ucsc.edu/~jcarmstr/datastore/progressiveCactusEvolverSim.tar.gz). We used these directories directly, except that root/seq.fa was truncated to 120,000 bp, corresponding to three simulated chromosomes of 40,000 bp each. The root genome sequence, Evolver configuration files, and simulation scripts used in this study are available in BioStudies (accession: S-BSST3394). The command to generate simulated genome is following and TrueTree.tre should be replaced with the Newick-formatted species tree used for the simulation.

simCtrl_runSim.py --inputNewick TrueTree.tre --outDir Sim${i} --rootDir root/ --rootName hg18 --paramsDir params/ --jobTree jobtreeSim${i} --maxThreads 8 --seed ${i} --noMEs True

**3) Post-simulation extraction**

After simulation, we extracted the simulated genomes using command: simCtrl_postSimFastaExtractor.py --simDir Sim${i}.

And we extracted the true alignment using command: simCtrl_postSimMafExtractor.py --simDir Sim${i}, which are both Evolver post-processing scripts. The true alignment is denoted as hg18.maf, where a block may contain multiple sequence records per taxon.

**4) Single-guide tree alignments using Progressive Cactus**

We aligned simulated genomes using cactus v3.3.0 and produced HAL outputs. Guide trees were specified in aln.txt.

cactus --realTimeLogging --defaultMemory 20G --maxCores 15 --maxMemory 30G ./jobstoreclusteraln ./aln.txt ./aln.hal

For alignments using star-like guide trees, we modified the Cactus configuration file by setting `allow_multifurcations="1"` under the `<decomposition>` section (the default setting is `allow_multifurcations="0"`), thereby allowing internal nodes with more than two children. The modified configuration file was supplied to Cactus using the `--configFile` option. All other configuration parameters were kept unchanged.

**5) HAL-to-MAF conversion and single-copy normalization**

To standardize downstream processing, we converted HAL to a chicken-referenced MAF and enforced at most one record per taxon per block using outType single. Here chicken is a fixed reference label in our simulation setup. It defines the coordinate system for reference-based scanning and consensus extraction.

For predicted alignments:

cactus-hal2maf --refGenome chicken --noAncestors --targetGenomes ${taxon set} --outType single --chunkSize 500000 jobstorealn aln.hal aln.maf

We used cactus v3.3.0 for HAL-to-MAF conversion because it provides the required duplicate-handling behavior.

**6) MAF to concatenated FASTA**

We converted single-guide tree and the consensus alignments from MAF to concatenated FASTA using maf_to_concat_fasta.py from bx-python:

maf_to_concat_fasta species1,species2,... < input.maf > output.fasta

The processes to get FASTA files from true alignments (hg18.maf) are different between 4-taxon and multi-taxon simulations and will be described separately.

**7) Tree reconstruction of concatenated alignments**

We inferred maximum-likelihood (ML) topologies from concatenated FASTA alignments using IQ-TREE v1.6.12. We enabled ModelFinder (Kalyaanamoorthy et al. 2017) for substitution model selection and the command was: iqtree -s input.fasta -ninit 1.

**8) The precision calculation using mafComparator**

Because the Evolver truth MAF (hg18.maf) contains ancestral and internal (non-leaf) sequences, we first removed non-leaf records using maffilter v1.3.1 (Dutheil et al. 2014) and generated a leaf-only truth MAF (hg18.noance.maf). The command we ran was maffilter param=filter.bpp. We used the following filter.bpp template (replace ${taxon set} with the taxon list used in the simulation):

DATA=hg18
input.file=./$(DATA).maf
input.file.compression=none
input.format=Maf
output.log=$(DATA).maffilter.log
maf.filter=Subset(species=(${taxon set}),strict=no,keep=no,remove_duplicates=no),Output(file=$(DATA).noance.maf,compression=none,mask=no)

We computed alignment precision following the Alignathon definition using mafComparator from mafTools (https://github.com/dentearl/mafTools). For each evaluated alignment (alignment.maf), we ran mafComparator against hg18.noance.maf, the command was: mafComparator --maf1 hg18.noance.maf --maf2 alignment.maf --out result.xml. The precision value in the XML output was the average attribute under

<homologyTests fileA="alignment.maf" fileB="hg18.noance.maf">
<aggregateResults>

<all totalTests="···" totalTrue="···" totalFalse="···" average="${precision}"/>
</aggregateResults>

### 2.2 Details of 4-taxon simulations

**1) The Newick formats of trees we used as true trees in Evolver:**

a) Realistic scenario:

(chicken:0.06804, (turkey vulture:0.00987, (seriema:0.0165, medium ground-finch:0.04854):0.0003):0.0123);

b) Felsenstein tree scenario:

(chicken:0.04, (turkey vulture:0.0005, (seriema:0.0005, medium ground-finch:0.05):0.0005):0.01);

c) Farris tree scenario:

(chicken:0.06804, (turkey vulture:0.04854, (seriema:0.0165, medium ground-finch:0.00987):0.0003):0.00123);

d) Clock-like tree scenario:

(chicken:0.0265, (turkey vulture:0.02527, (seriema:0.02497, medium ground-finch:0.02497):0.0003):0.00123);

**2) True alignment processing**

Because hg18.maf can contain multiple records per taxon within a block, direct concatenation would cause maf_to_concat_fasta.py to keep only the first record per taxon. We therefore exhaustively enumerated one sequence choice per taxon within each block, counted site patterns directly from the enumerated columns, and wrote the concatenated FASTA from the site pattern counts.

**3) Guide trees used in Progressive Cactus to generate single-guide tree alignments**

We used all three fully resolved binary trees and one star tree, see Figure S5.

**4) Site pattern counting and classifying for Jensen-Shannon (JS) divergence**

a) Column filtering and normalization

Site patterns were counted from concatenated FASTA alignments after excluding any column that contains a gap character (-) or an ambiguous base (N) in any taxon. Counts were then normalized into empirical distributions for JS divergence comparisons.

b) Five category definitions:

We used five coarse categories defined on gap-free and N-free columns:

Invariant: exactly one nucleotide present (e.g., AAAA).

Singleton: exactly two nucleotides present with counts 3+1 (e.g., AAAG).

Parsimony-informative: at least two nucleotides present and at least two nucleotides have counts $\geqslant 2$ (e.g., AAGG).

Three-nucleotide: exactly three distinct nucleotides present (e.g., AATG).

Four-nucleotide: exactly four distinct nucleotides present (e.g., AGTC).

c) Jensen–Shannon divergence

To quantify the deviation of concatenated alignments from the truth in site-pattern frequency distributions, we calculated the Jensen–Shannon divergence (Lin 1991). When counting site patterns, we excluded columns containing gaps or ambiguous base N. Let $\mathcal{X}$ be the set of site pattern categories. Let $P$ and $Q$ be empirical distributions over $\mathcal{X}$ from the truth and the evaluated alignment, obtained by normalizing site pattern counts. With $M = \frac{1}{2}(P + Q)$,

$$JS(P||Q) = \frac{1}{2}\sum_{x\in\mathcal{X}} P(x) log \frac{P(x)}{M(x)} + \frac{1}{2}\sum_{x\in\mathcal{X}} Q(x) log \frac{Q(x)}{M(x)}.$$

We computed the JS divergence for all individual site patterns, singleton patterns, parsimony-informative patterns and subtypes, and five coarse categories: invariant, singleton, parsimony-informative, three-nucleotide, and four-nucleotide.

#### 2.3 Details of 12-taxon simulations

**1) The Newick formats of trees we used as true trees in Evolver:**

(chicken:0.0652738014,((((grey crowned crane:0.0173370903,killdeer:0.0175099290):0.0005205566,hoatzin:0.0229300404):0.0003284769,((seriema:0.0163967970,turkey vulture:0.0093447350):0.0014081682,(sunbittern:0.0273341387,red-throated loon:0.0136485311):0.0005313832):0.0002658022):0.0003808777,(chimney swift:0.0334481075,MacQueen's bustard:0.0227700211):0.0004268751):0.0003632007,(pigeon:0.0293869380,American flamingo:0.0135891416):0.0003677007);

**2) True alignment processing**

Exhaustive enumeration like 4-taxon simulation was computationally infeasible because a block may contain multiple sequence records per taxon. We therefore converted hg18.maf to HAL and re-exported a single-copy truth MAF hg18.fromhal.maf using the same cactus-hal2maf --outType single workflow.

maf2hal --refGenome hg18 hg18.maf hg18.hal

cactus-hal2maf --refGenome chicken --noAncestors --outType single --chunkSize 500000 jobstoretruth hg18.hal hg18.fromhal.maf

We then used maf_to_concat_fasta.py to convert hg18.fromhal.maf to the FASTA file using command:

maf_to_concat_fasta species1,species2,... < hg18.fromhal.maf > hg18.fasta

**3) Guide trees used in Progressive Cactus to generate single-guide tree alignments**

We generated guide trees 2-10 in Figure S17 using a custom script in strict mode where RF distance is 1, triplet distance threshold is 2/3, and the taxa evolve along yule mode.

The command used was: python generate_random_guidetrees_2models_2modes_usethis_fast2_finalver.py --taxa grey_crowned_crane killdeer " (sunbittern,red-throated_loon)" American_flamingo chimney_swift "(seriema,turkey_vulture)" MacQueen's_bustard hoatzin pigeon --outgroup chicken --num_trees 9 --max-tries 20000000 --restarts 10 --seed 399304004

We used one true tree (tree 1 in Figure S17), and a partially resolved tree and a purely star tree as single-guide trees.

**4) A control experiment filtering gaps and paralogs**

As a control experiment, we filter out paralogs (completely deleting multiple sequences from the same taxon within one aligned block), sites with more any or at least two gaps, respectively, and short aligned blocks (deleting aligned blocks with length below 450bp), reducing the average number of nucleotides to a value similar to consensus_of_4. The performance is shown in Figures S18-S21. Paralogous regions were removed using a custom script. Sites with excessive gaps and short aligned blocks were filtered using maffilter (param=maffilter.bpp) (Dutheil et al. 2014).

Gap filtering was performed using the AlnFilter2 function in maffilter. For the main analysis, we used max.gap=0.08 with relative=yes, which removes columns containing more than zero gaps among the 12 selected taxa. A relaxed gap threshold (max.gap=0.12) was additionally evaluated, allowing columns containing at most one gap to be retained. The parameter file used for gap filtering is shown below.

```
input.file=ALN.maf
input.file.compression=none
input.format=Maf
output.log=ALN.log
maf.filter=\
    AlnFilter2(\
species=(GALGA,COLLI,PHORU,CHAPE,CHLUN,BALRE,CHAVO,OPHHO,GAVST,EURHE,CATAU,CAR
CR),\
        window.size=1,\
        window.step=1,\
        max.gap=0.08,\ (or 0.12)
        max.pos=0,\
        missing_as_gap=1,\
        relative=yes),\
    Output(\
        file=./ALN.08.maf,\ (or ALN.12.maf)
        compression=none,\
        mask=yes)
```

Short aligned blocks were removed using the MinBlockLength function in maffilter, retaining only aligned blocks with length ≥450 bp. The corresponding parameter file is shown below.

```
input.file=ALN.maf
input.file.compression=none
input.format=Maf

output.log=ALN.log

maf.filter=\
    MinBlockLength(\
        min_length=450),\
    Output(\
        file=ALN.blockover450.maf,\
        compression=none,\
        mask=yes)
```

**5) Calculate the recovery of true tree splits in trees inferred by each alignment method**

Unlike the four-taxon simulations, where we counted the frequencies of all three possible quartet topologies, we evaluated the multi-taxon simulations by counting how often each true-tree split was recovered in trees inferred from single-guide-tree alignments, consensus alignments, and true alignments. To quantify split recovery, we represented each internal split of the true tree as a binary vector under a fixed taxon order, where taxa on one side of the split were coded as 1 and taxa on the other side as 0. Trivial splits separating a single taxon from all remaining taxa were excluded.

For each inferred tree, we generated the corresponding set of non-trivial split vectors using the same taxon order. A true-tree split was counted as recovered if its binary vector matched any split vector in the inferred tree, either in the same orientation or in the complementary orientation, because the two sides of a split are unordered. We assigned a score of 1 when the split was recovered and 0 otherwise. For each alignment method, this procedure was repeated across 100 simulation replicates, and the recovery count for each true-tree split was calculated as the sum of the 0/1 scores across replicates.

**6) Erroneous fraction among removed homology pairs**

For the 12-taxon simulations, homology pairs were extracted from each single guide-tree alignment, the corresponding consensus alignment, and the true alignment. Homology pairs present in the single-guide-tree alignment but absent from the consensus alignment were considered removed pairs. Among these removed pairs, those that were also absent from the true alignment were classified as erroneous removed pairs. The erroneous fraction among removed homology pairs was calculated as the number of erroneous removed pairs divided by the total number of removed pairs. This calculation was performed independently for each replicate and each single guide-tree alignment.

#### 2.4 Details of 43-taxon simulations

**1) The Newick formats of trees we used as true trees in Evolver:**

((((((((((kea:0.01261,budgerigar:0.01755)interior2:0.01584,((((medium ground-finch:0.01116,zebra finch:0.01259)interior7:0.007341,American crow:0.01154)interior6:0.01321,golden-collared manakin:0.02079)interior5:0.002707,rifleman:0.02507)interior4:0.01166)interior3:0.0006309,peregrine falcon:0.02287)interior8:0.0003867,seriema:0.0165)interior9:0.0004503,((turkey vulture:0.009021,bald eagle:0.0131)interior12:0.0005931,(barn owl:0.02026,((cuckoo-roller:0.02009,((rhinoceros hornbill:0.02993,(downy woodpecker:0.04302,carmine bee-eater:0.0309)interior18:0.00117)interior17:0.0007587,bar-tailed trogon:0.03144)interior16:0.0003669)interior15:0.0004074,speckled mousebird:0.03615)interior14:0.0001865)interior13:0.0001797)interior11:0.0002409)interior10:0.001244,((white-tailed tropicbird:0.01772,sunbittern:0.02841)interior34:0.0009534,(red-throated loon:0.01334,((great cormorant:0.01917,(little egret:0.01686,dalmatian pelican:0.01249)interior39:0.0003993,crested ibis:0.01195)interior38:0.0002912)interior37:0.0004821,((adelie penguin:0.004059,emperor penguin:0.003273)interior41:0.006426,northern fulmar:0.01025)interior40:0.0003576)interior36:0.0005697)interior35:0.0006258)interior33:0.0003579)interior19:0.0003345,(hoatzin:0.02267,(killdeer:0.01805,grey crowned crane:0.01712)interior22:0.0004527)interior21:0.0002767)interior20:0.000303,(((red-crested turaco:0.02124,MacQueen's bustard:0.02195)interior30:0.0004335,common cuckoo:0.03345)interior29:0.0003174,((chimney swift:0.02723,Anna's hummingbird:0.03378)interior32:0.006108,chuck-will's-widow:0.0209)interior31:0.001138)interior28:0.0003384)interior23:0.0003216,((American flamingo:0.009351,Great crested grebe:0.01753)interior25:0.003891,(pigeon:0.0289,(yellow-throated sandgrouse:0.0215,brown mesite:0.02614)interior27:0.0008157)interior26:0.0004233)interior24:0.0002782)interior42:0.0096,chicken:0.05

748);

**2) Command used to run the hierarchical consensus pipeline and guide trees we used**

We ran the hierarchical consensus pipeline described in Figure 4B of the main text using the code provided on the GitHub page. The command was:

bash make_plan_UseThis.sh --planner plan_partially_resolved_cactus.py --tree-file aln.tre --reference GALGA --paths genome.txt --threads 15 --common_workers 5

The file aln.tre contains the global guide tree shown in Figure S24, and genome.txt provides the taxon names and paths to the corresponding genome files. The script automatically generated local guide trees for each polytomy and output commands that could be copied and run directly.

For the 100 simulation replicates, this command was initially run separately for each replicate, with local guide trees generated randomly each time. For ease of comparison and summary across replicates, we then used the local guide trees generated for replicate 1 in all other replicates, so that the same set of local guide trees was used across all replicates. These local guide trees are shown in Figure S25-S29.

The four globally fully resolved guide trees used for comparison were constructed by combining, across all polytomies, the first, second, third, or fourth local guide tree generated for each polytomy, respectively (Figure S30). For example, the guide tree used to generate aln1 in Figure S30 was obtained by combining the first local guide tree from each polytomy shown in Figure S26-S29.

**3) True alignment processing and the recovery of true-tree splits are the same as that in 12-taxon simulation.**

**Note 3: Details of applying method on real dataset**

**3.1 Real data resources and reference setting**

We used the avian genomes provided by Jarvis 2014 and analyzed the downloaded genomes and associated files. The genome files are downloaded from https://s3.ap-northeast-1.wasabisys.com/gigadb-datasets/live/pub/10.5524/100001_101000/101000/bird_phylogenomics_data.tar.gz.

We also uploaded the genomes to the EMBL-EBI repository BioStudies (accession S-BSST3394).

Prior to whole-genome alignment, genome assemblies were soft-masked to reduce the potential impact of repetitive regions on alignment. For each genome assembly, a species-specific repeat library was first generated de novo using RepeatModeler v2.0.3 (Flynn et al. 2020) with the LTR structural analysis option (-LTRStruct). The resulting repeat library was then used to soft-mask the corresponding genome assembly using RepeatMasker v4.1.2-p1 (Smit et al. 2013–2015) with the -xsmall option. The following commands were used as an example for chicken (GALGA) genome assembly:

BuildDatabase -name GALGA GALGA.fa

RepeatModeler -database GALGA -pa 20 -LTRStruct

RepeatMasker -lib GALGA-families.fa -s -pa 20 -xsmall GALGA.fa

**3.2 Guide-tree generation under five-group constraints**

To assess guide-tree effects in progressive whole-genome alignment, we generated four fully resolved guide trees under predefined five-group constraints. Two groups had fixed internal structure, and the remaining relationships were completed in strict mode (RF distance=1, triplet distance>=2/3). The guide trees used are shown in Figure 5A in the main text.

**3.3 Progressive Cactus alignments and HAL-to-MAF conversion**

We ran Progressive Cactus under each of the four guide trees to produce four HAL alignments. The Progressive Cactus version and key parameters matched those used in the simulation section. Each HAL was converted to a chicken-reference MAF using cactus-hal2maf.

**3.4 Concatenation analyses: IQ-TREE, PAUP*, and CASTER-site**

Before phylogenetic reconstruction, we removed alignment sites containing no unambiguous nucleotide characters (i.e., sites consisting only of gaps and/or ambiguous characters) and excluded regions corresponding to the chicken (Gallus gallus) Un_random sequences.

**1) Concatenated FASTA**

We converted MAF alignments to concatenated FASTA for concatenation-based inference and scoring. Scripts were the same as what was used in simulations.

**2) IQ-TREE inference with 10,000-site partitions**

We applied a partitioned model by splitting each concatenated alignment into consecutive partitions of 10,000 alignment sites. We used IQ-TREE v1.6.12 with GTR+F+I model under NEXUS partition format. Result trees are shown in Figure S38. The command was iqtree -s ${alignment}.fasta -spp ${alignment}_partitions.txt -pre ${alignment}_partition10kGTRFI -ninit 1 -quiet -nt 30 -m GTR+F+I

**3) CASTER-site**

We ran CASTER-site on concatenated FASTA alignments, result trees are shown in Figure S39.

Free mode:

The command we used was: caster-site -i alignment.fasta -o caster_site.txt 2> caster_site.log

**4) PAUP* maximum parsimony inference**

We used PAUP* test version (Version 4.0a build 168 for Unix/Linux) with branch-and-bound search. Input NEXUS files converted from FASTA was initially performed with EMBOSS seqret (v6.6.0.0) (Rice et al. 2000), however, for the largest alignments (~9 GB), we used a custom streaming Python script to avoid loading entire sequences into memory. The script verified sequence-length consistency across taxa and generated the corresponding NEXUS. Result trees are shown in Figure S40.

The command we used to do the unconstrained search:

```
paup4a168_centos64 <ALIGNMENT>.nex
paup> BandB
paup> SaveTrees file=<ALIGNMENT>_mp.tree replace=yes;
```

#### 3.5 Loci-ASTRAL pipeline

Step1: 10 kb genomic partitions and 1 kb occupancy windows

We defined consecutive 10 kb genomic regions on chicken chromosomes. Within the first 5 kb of each region, we scanned 1 kb sliding windows and selected the window with the highest occupancy, where occupancy was defined as the fraction of non-gap characters.

Step2: Gene trees and species tree inference by ASTRAL

We inferred locus gene trees using IQ-TREE with ModelFinder and summarized gene trees using ASTRAL v5.7.8. (Result trees are shown in Figure S41).

java -jar astral.5.7.8.jar -i ./all_loci_trees.tre -o astral_loci_trees.tre

#### 3.6 Comparison between current and previous real data analyses

In an earlier version of this manuscript (available at https://www.biorxiv.org/content/10.64898/2026.07.06.736671v1), we analyzed whole chromosome alignments for chicken chromosome 1 and the same taxon set as here.

In addition to including the whole genome, we used an updated version of Progressive Cactus (v-3.3.0 vs v-2.1.1), and we did soft repeat mask using RepeatMasker on all genomes before aligning. The main difference with respect to phylogenetic analysis is that the mousebird was classified as the outgroup to other Afroaves. In Figure S3, we show the support for the three quartets formed by chicken (A), mousebird (B), Cavitaves (C, three taxa), and Accipitrimorphae + barn owl (D, three taxa). For every locus tree, we compute weights for each of the quartets AD|BC, AB|CD and AC|BD by giving equal weight to all of the nine choices of one taxon from C and one taxon from D. We then used a sliding window to summarize the average support for 1,000 consecutive loci within each chromosome.

We find that some parts of the genome, e.g. long stretches within chromosomes 2, 5, 6, and Z, the quartet AD|BC, which is preferred by our current experiment dominates. AC|BD tends to be the least supported quartet throughout the genome, and AB|CD, which was preferred by our previous analysis, is generally well supported whenever AD|BC is not unusually strong. The difference in support across the genome is considerable but not as extreme as for an outlier region in chromosome 4 for a different phylogenetic question that was reported by Mirarab et al. (2024). We conclude that for resolving difficult phylogenetic questions like the mousebird position from whole genome alignments, these problems need to be resolved, for example, by filtering loci from outlier regions.

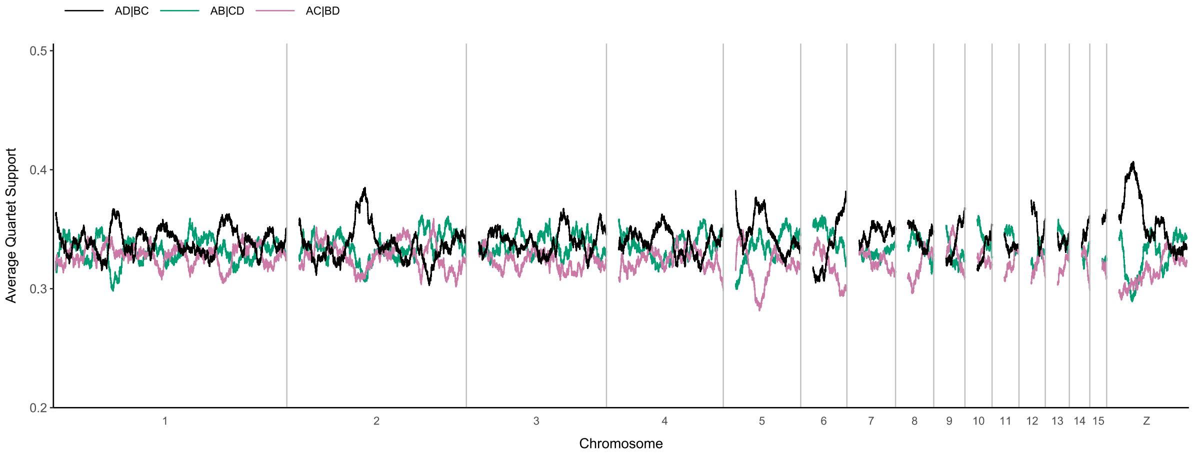


**Figure S3** Average quartet support across 1,000 consecutive loci for the first 15 and Z chicken chromosomes

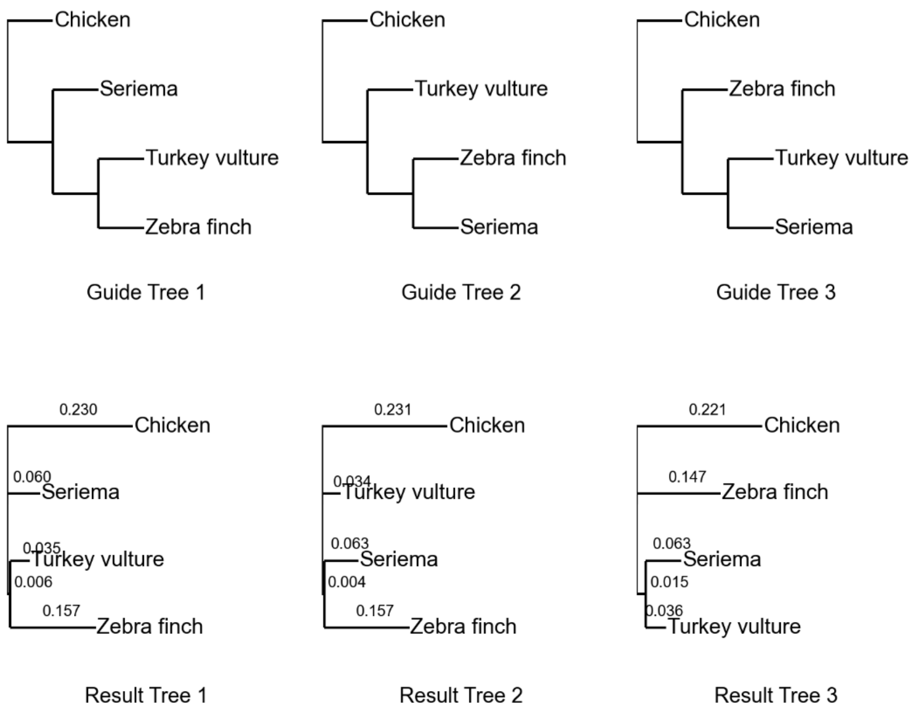


**Figure S4** Trees inferred from alignments generated under three binary guide trees of the 4-taxon set chicken, seriema, zebra finch, and turkey vulture.

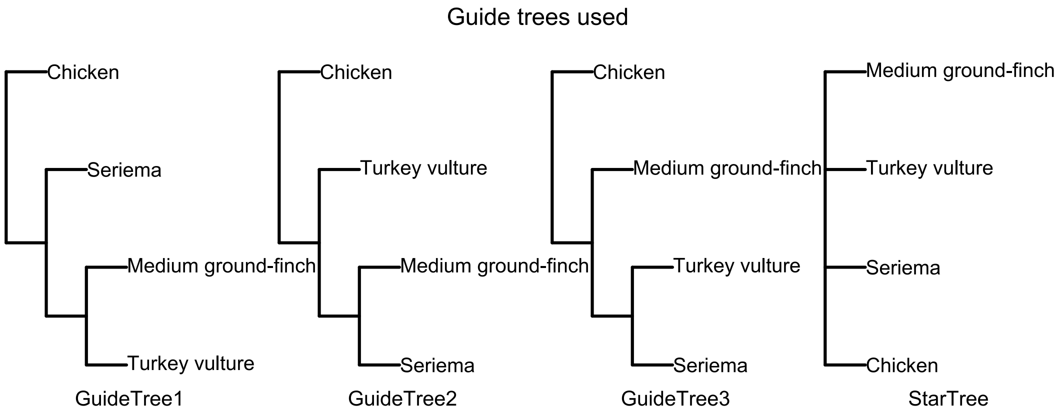


**Figure S5** Guide trees used in four-taxon simulation.

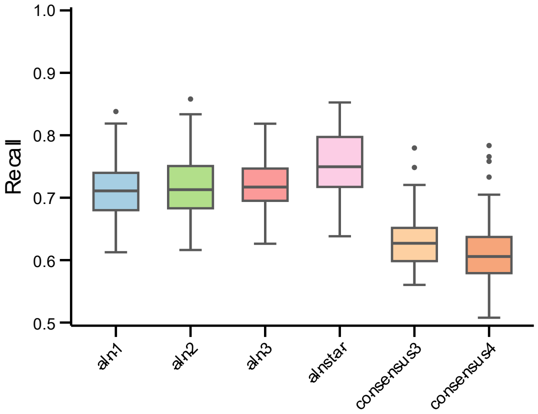


**Figure S6** Boxplot of recall of Realistic tree scenario.

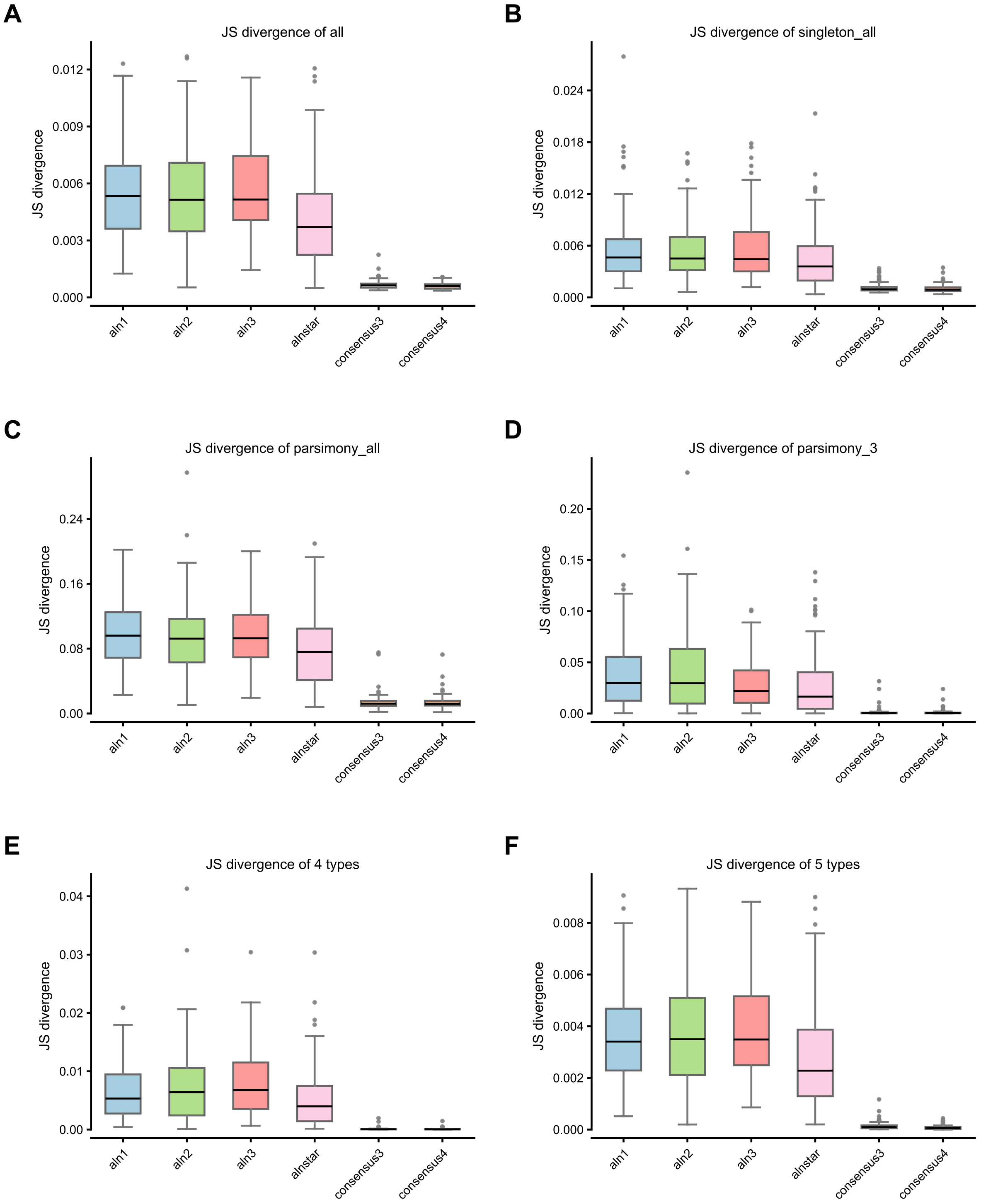


**Figure S7** All boxplots of JS-divergence at multiple levels of Realistic scenario. A, JS divergence of individual site patterns. B, JS divergence of individual singletons. C, JS divergence of individual parsimony-informative site pattern. D, JS-divergence of three parsimony-informative site-pattern classes supporting the three candidate topologies. E, JS-divergence of five broader site-pattern types. F, JS-divergence of four broader site-pattern types (five types excluding invariants).

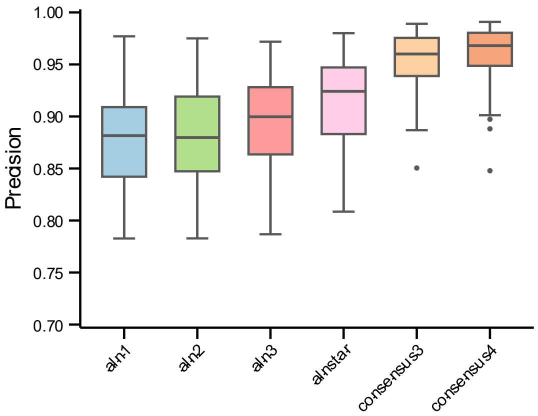


**Figure S8** Boxplot of precision of Felsenstein tree scenario.

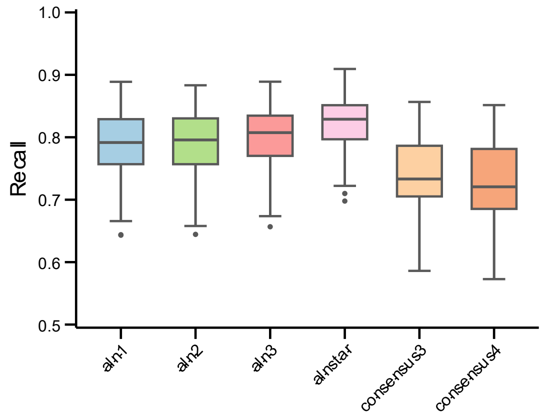


**Figure S9** Boxplot of recall of Felsenstein tree scenario.

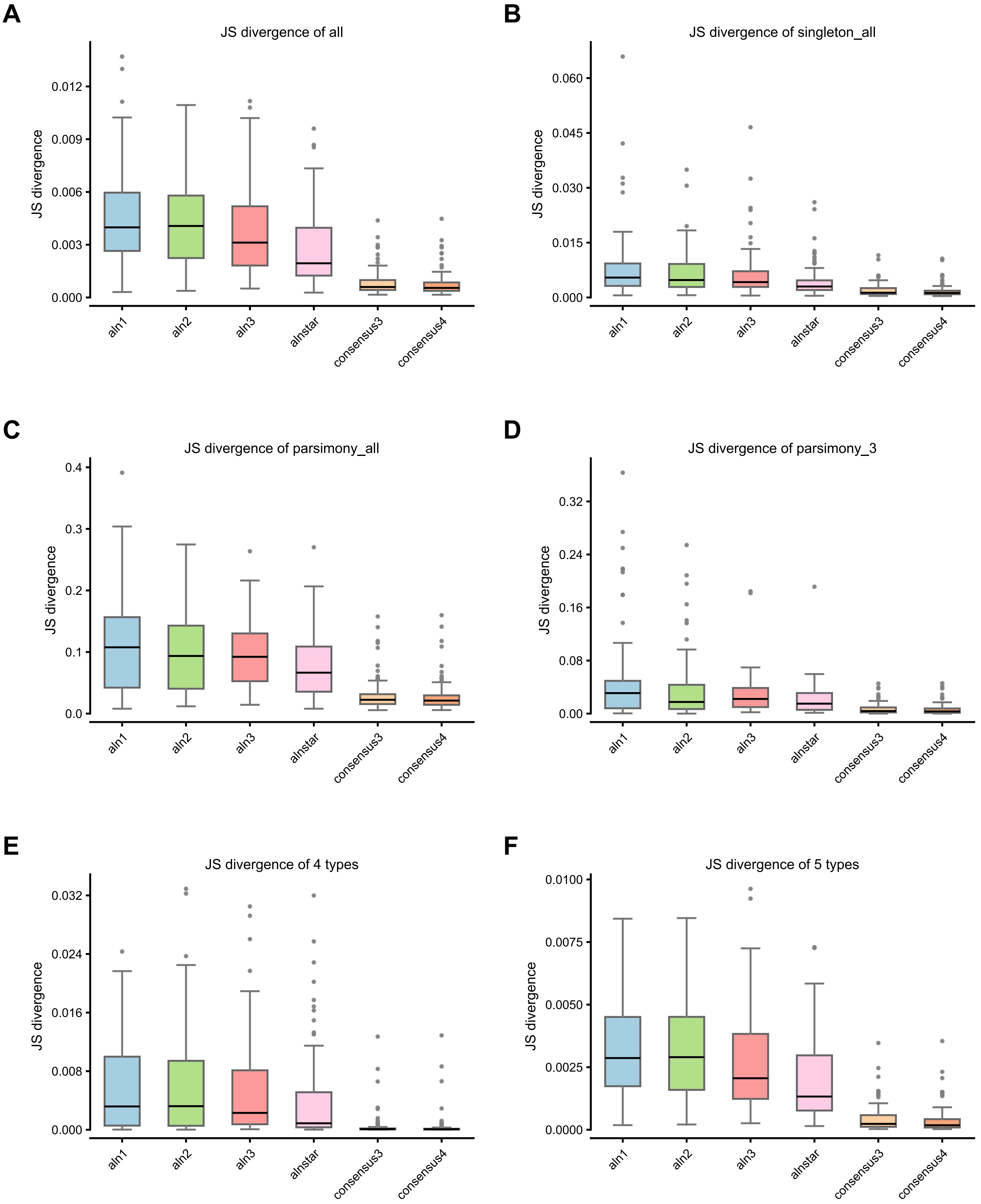


**Figure S10** All boxplots of JS-divergence at multiple levels of Felsenstein tree scenario. A, JS divergence of individual site patterns. B, JS divergence of individual singletons. C, JS divergence of individual parsimony-informative site pattern. D, JS-divergence of three parsimony-informative site-pattern classes supporting the three candidate topologies. E, JS-divergence of five broader site-pattern types. F, JS-divergence of four broader site-pattern types (five types excluding invariants).

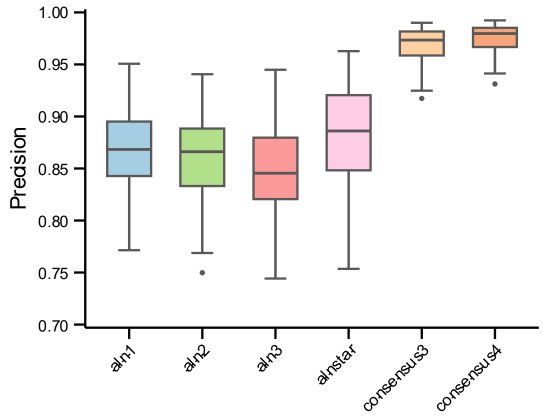


**Figure S11** Boxplot of precision of Farris tree scenario.

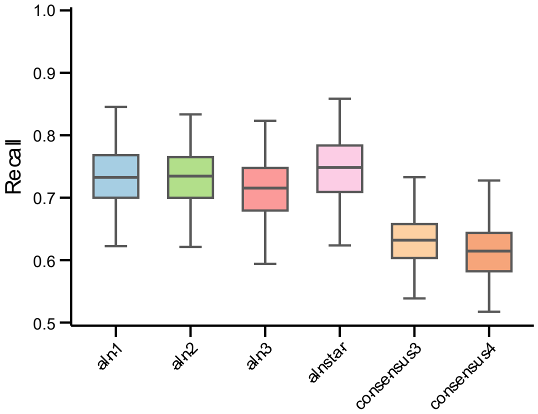


**Figure S12** Boxplot of recall of Farris tree scenario.

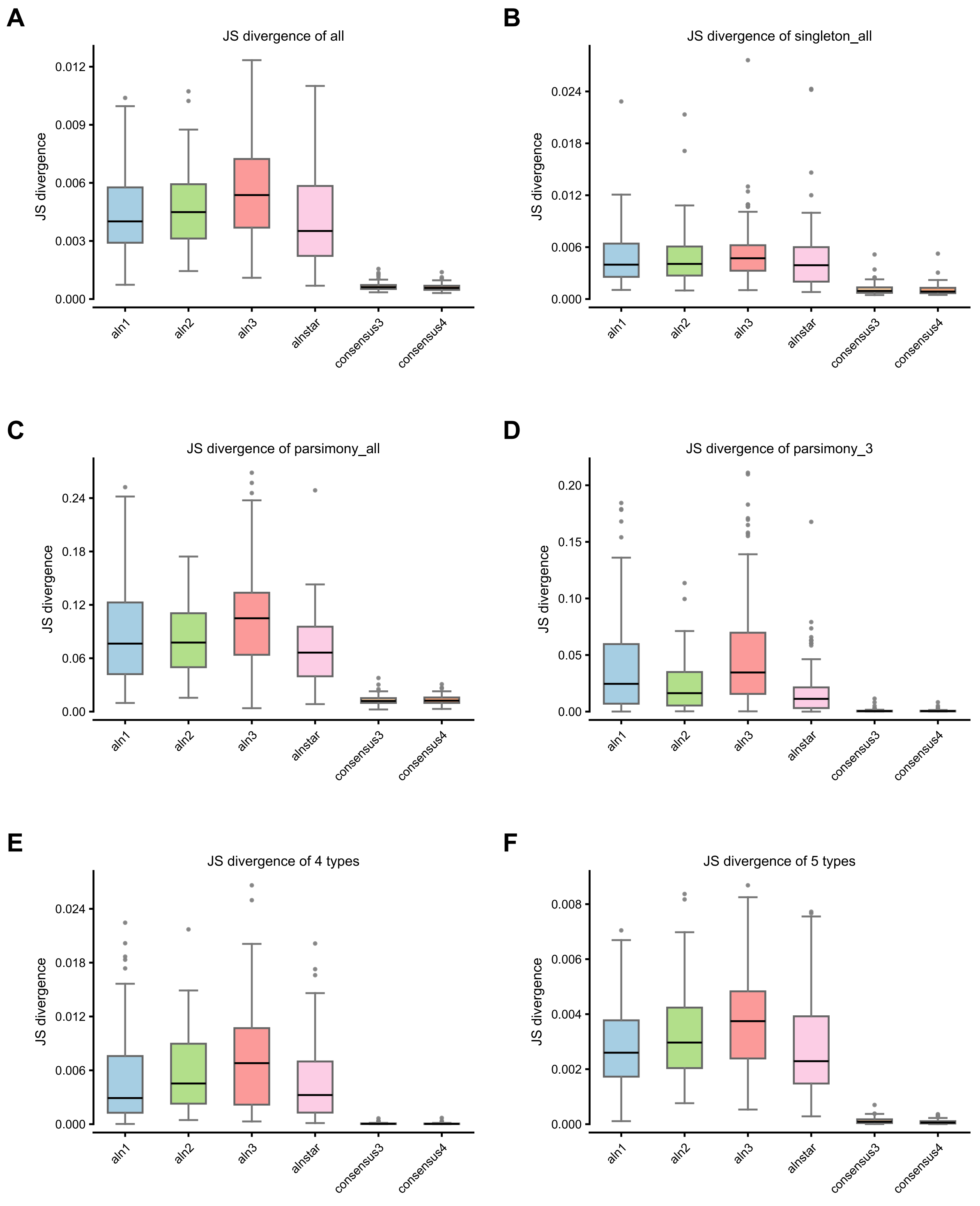


**Figure S13** All boxplots of JS-divergence at multiple levels of Farris tree scenario. A, JS divergence of individual site patterns. B, JS divergence of individual singletons. C, JS divergence of individual parsimony-informative site pattern. D, JS-divergence of three parsimony-informative site-pattern classes supporting the three candidate topologies. E, JS-divergence of five broader site-pattern types. F, JS-divergence of four broader site-pattern types (five types excluding invariants).

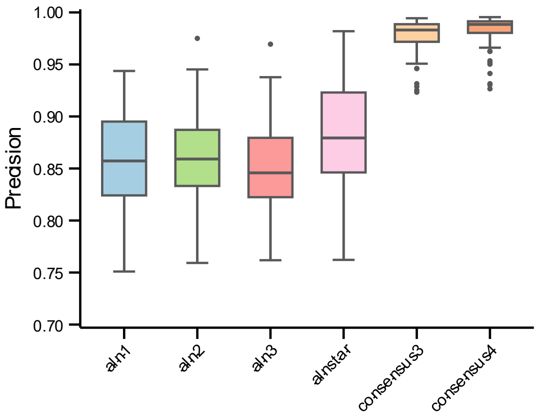


**Figure S14** Boxplot of precision of Clock-like tree scenario.

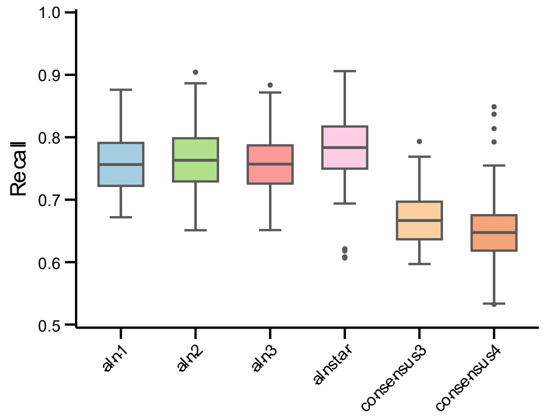


**Figure S15** Boxplot of recall of Clock-like tree scenario.

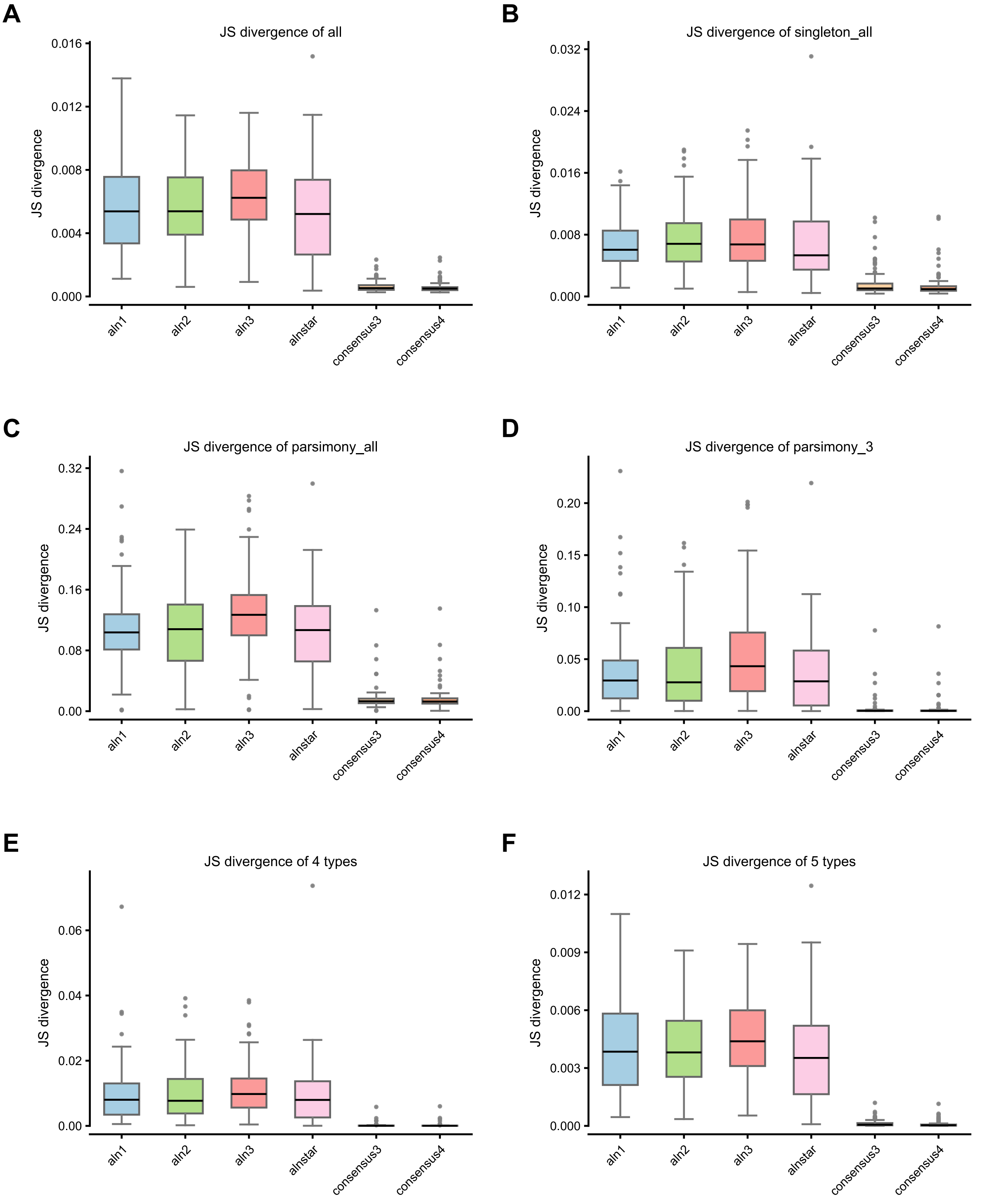


**Figure S16** All boxplots of JS-divergence at multiple levels of Clock-like tree scenario. **A**, JS divergence of individual site patterns. **B**, JS divergence of individual singletons. **C**, JS divergence of individual parsimony-informative site pattern. **D**, JS-divergence of three parsimony-informative site-pattern classes supporting the three candidate topologies. **E**, JS-divergence of five broader site-pattern types. **F**, JS-divergence of four broader site-pattern types (five types excluding invariants).

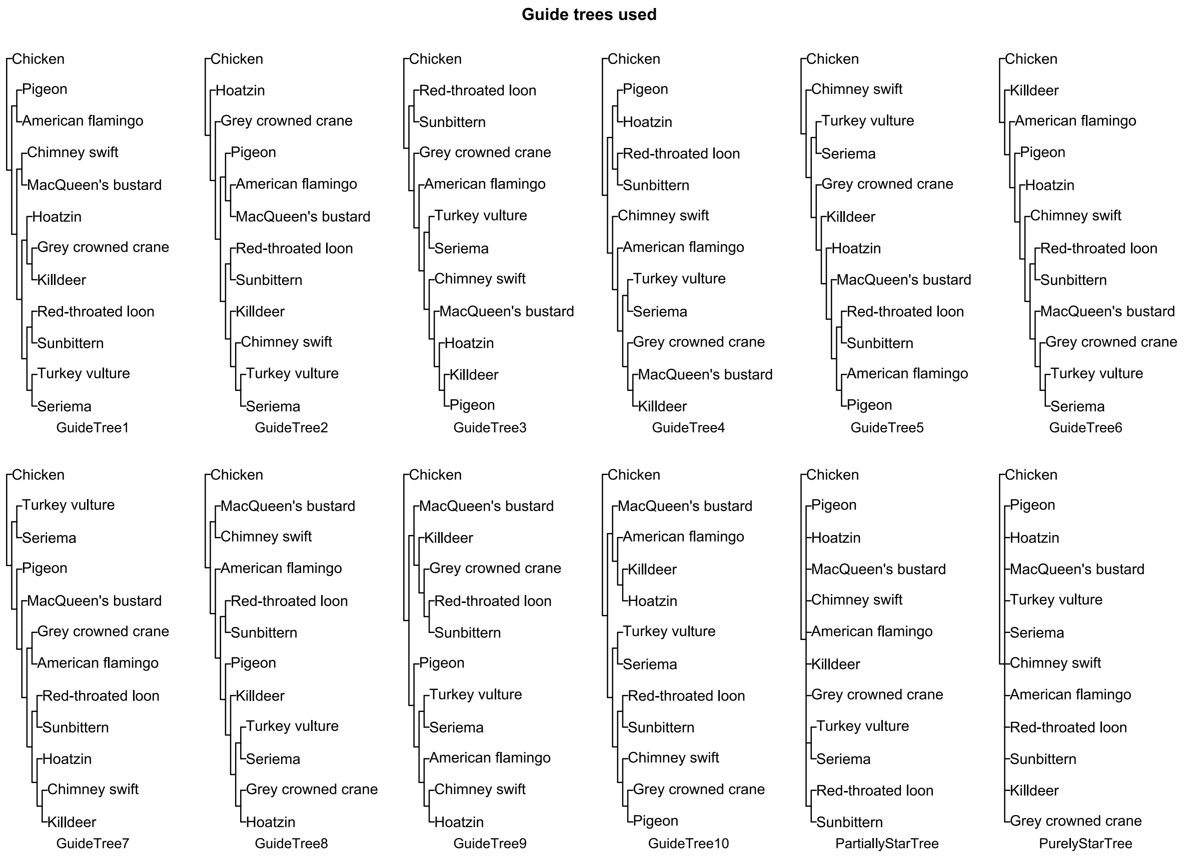


**Figure S17** Guide trees used in 12-taxon simulation, including ten resolved binary trees and two star trees.

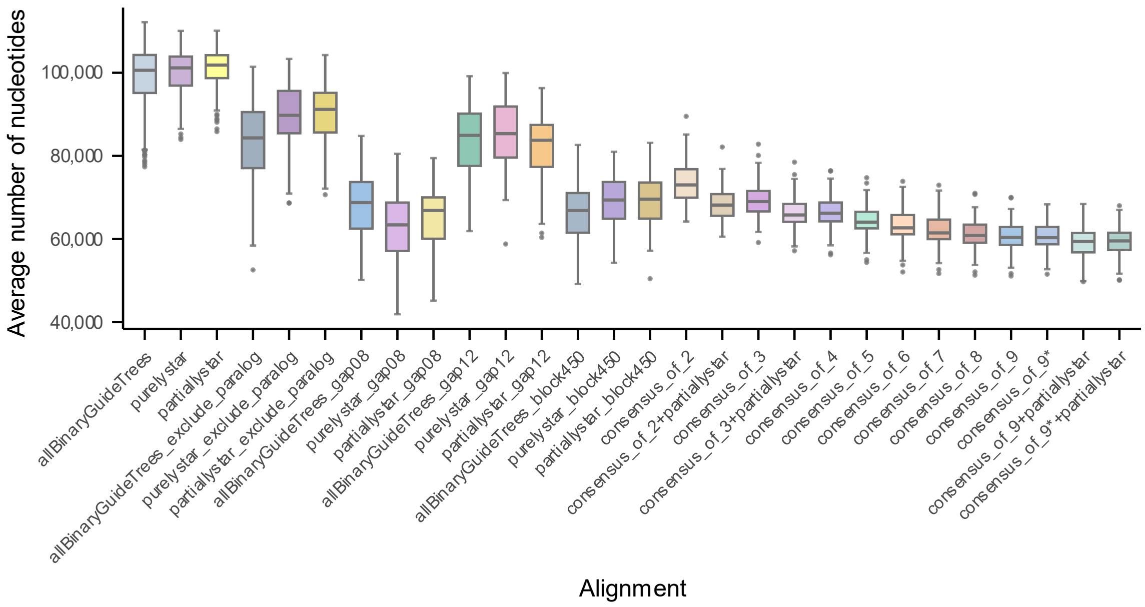


**Figure S18** Average number of nucleotides of all alignments including filtered ones in 12-taxon simulation, individual binary guide tree based alignments are pooled into a single box. An asterisk (*) indicates that the corresponding consensus alignment includes the alignment generated under the true tree as one of its input alignments. Alignment suffixes indicate filtering steps: _gap08, removal of gap-containing sites; _gap12, removal of sites with more than one gap; _block450, removal of aligned blocks shorter than 450 bp; and _exclude_paralog, removal of paralogous regions.

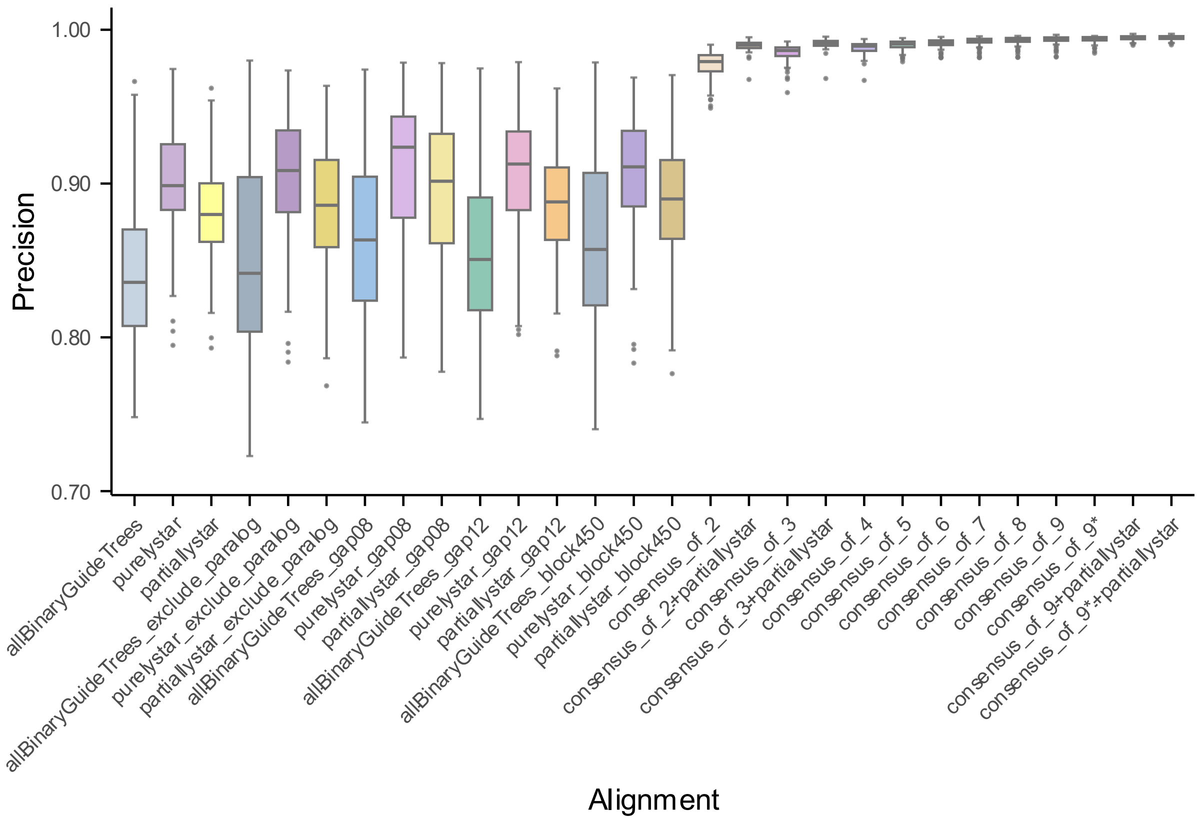


**Figure S19** Precision of all alignments including filtered ones in 12-taxon simulation, individual binary guide tree based alignments are pooled into a single box. An asterisk (*) indicates that the corresponding consensus alignment includes the alignment generated under the true tree as one of its input alignments. Alignment suffixes indicate filtering steps: _gap08, removal of gap-containing sites; _gap12, removal of sites with more than one gap; _block450, removal of aligned blocks shorter than 450 bp; and _exclude_paralog, removal of paralogous regions.

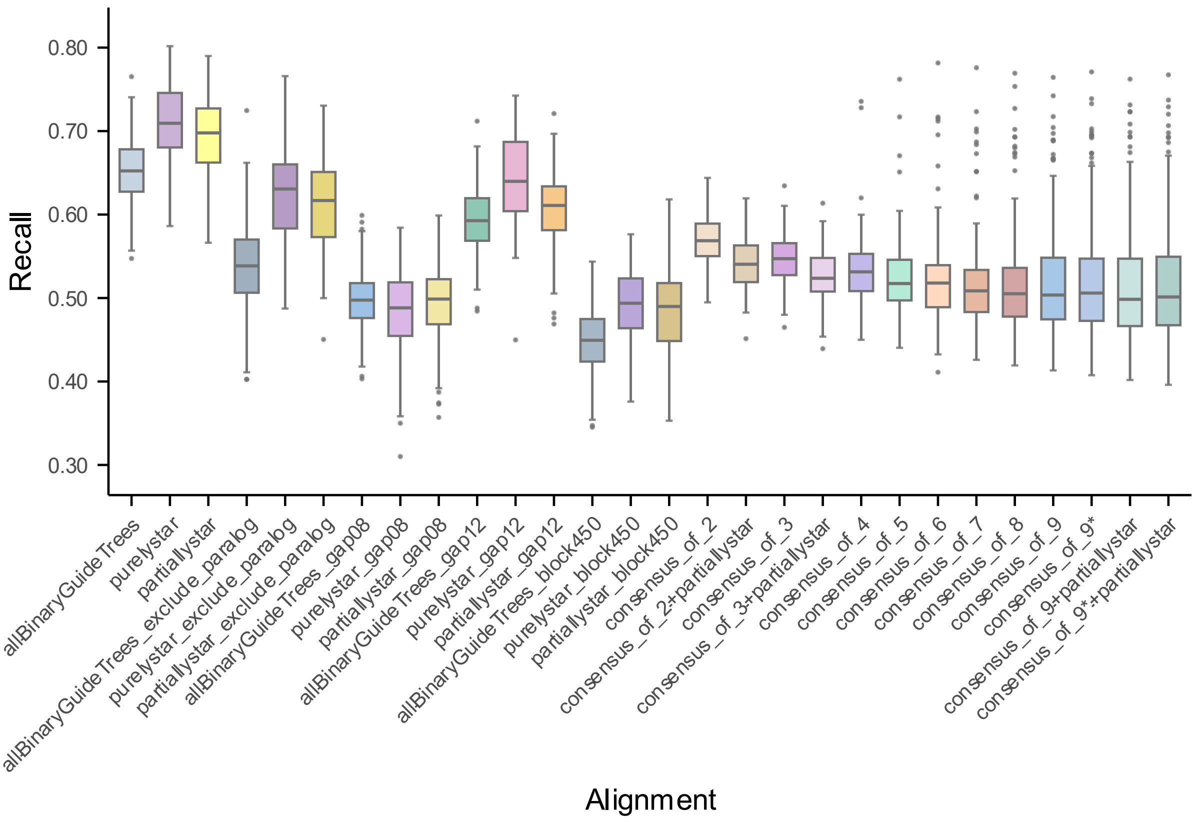


**Figure S20** Recall of all alignments including filtered ones in 12-taxon simulation, individual binary guide tree based alignments are pooled into a single box. An asterisk (*) indicates that the corresponding consensus alignment includes the alignment generated under the true tree as one of its input alignments. Alignment suffixes indicate filtering steps: _gap08, removal of gap-containing sites; _gap12, removal of sites with more than one gap; _block450, removal of aligned blocks shorter than 450 bp; and _exclude_paralog, removal of paralogous regions.

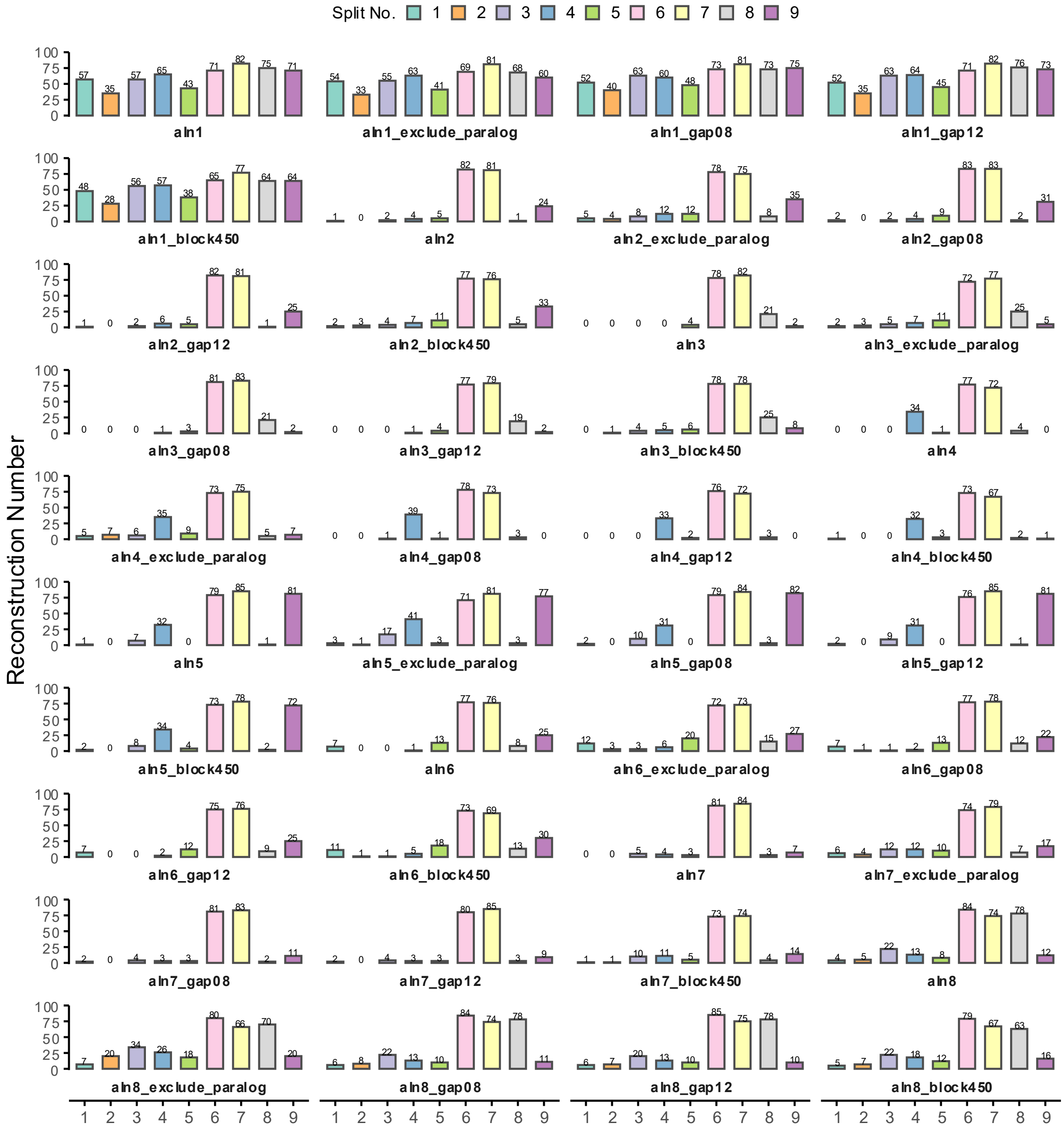


**Figure S21** Numbers of true-tree splits recovered from phylogenetic trees inferred from all alignments including filtered ones in 12-taxon simulation. An asterisk (*) indicates that the corresponding consensus alignment includes the alignment generated under the true tree as one of its input alignments. Alignment suffixes indicate filtering steps: _gap08, removal of gap-containing sites; _gap12, removal of sites with more than one gap; _block450, removal of aligned blocks shorter than 450 bp; and _exclude_paralog, removal of paralogous regions.

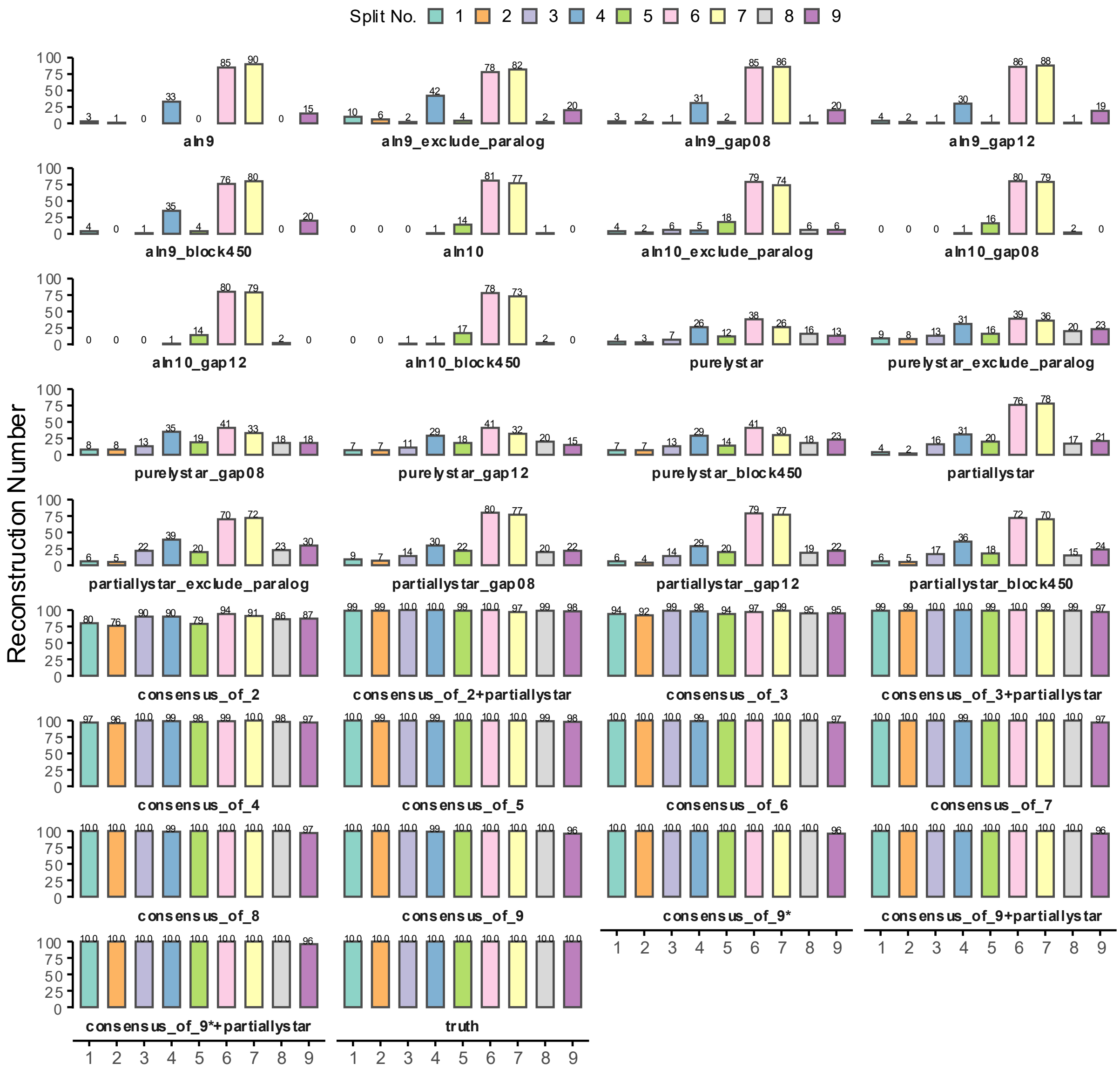


**Figure S21** (continued) Numbers of true-tree splits recovered from phylogenetic trees inferred from all alignments including filtered ones in 12-taxon simulation. An asterisk (*) indicates that the corresponding consensus alignment includes the alignment generated under the true tree as one of its input alignments. Alignment suffixes indicate filtering steps: _gap08, removal of gap-containing sites; _gap12, removal of sites with more than one gap; _block450, removal of aligned blocks shorter than 450 bp; and _exclude_paralog, removal of paralogous regions.

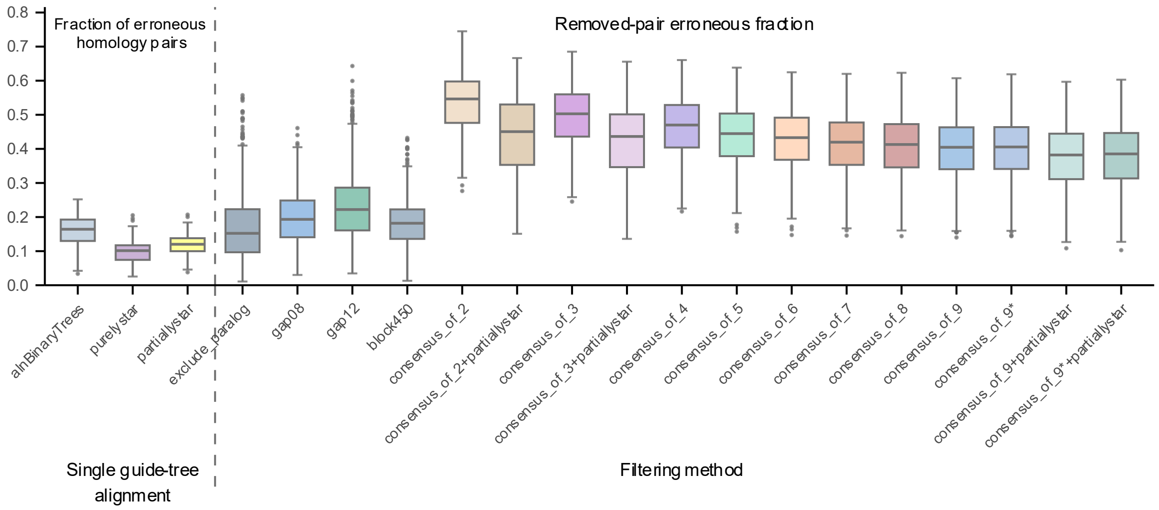


**Figure S22** Fraction of erroneous homology pairs in the simulated 12-taxon dataset. The left side of the dashed line shows the fraction of erroneous pairs in the original alignments, while the right side shows the fraction of erroneous pairs among those removed by different filtering and consensus methods. Alignments generated using individual binary guide trees are pooled into a single box. An asterisk (*) indicates that the corresponding consensus alignment includes the alignment generated under the true tree as one of its input alignments. Simple filtering steps: gap08, removal of gap-containing sites; gap12, removal of sites with more than one gap; block450, removal of aligned blocks shorter than 450 bp; and exclude_paralog, removal of paralogous regions.

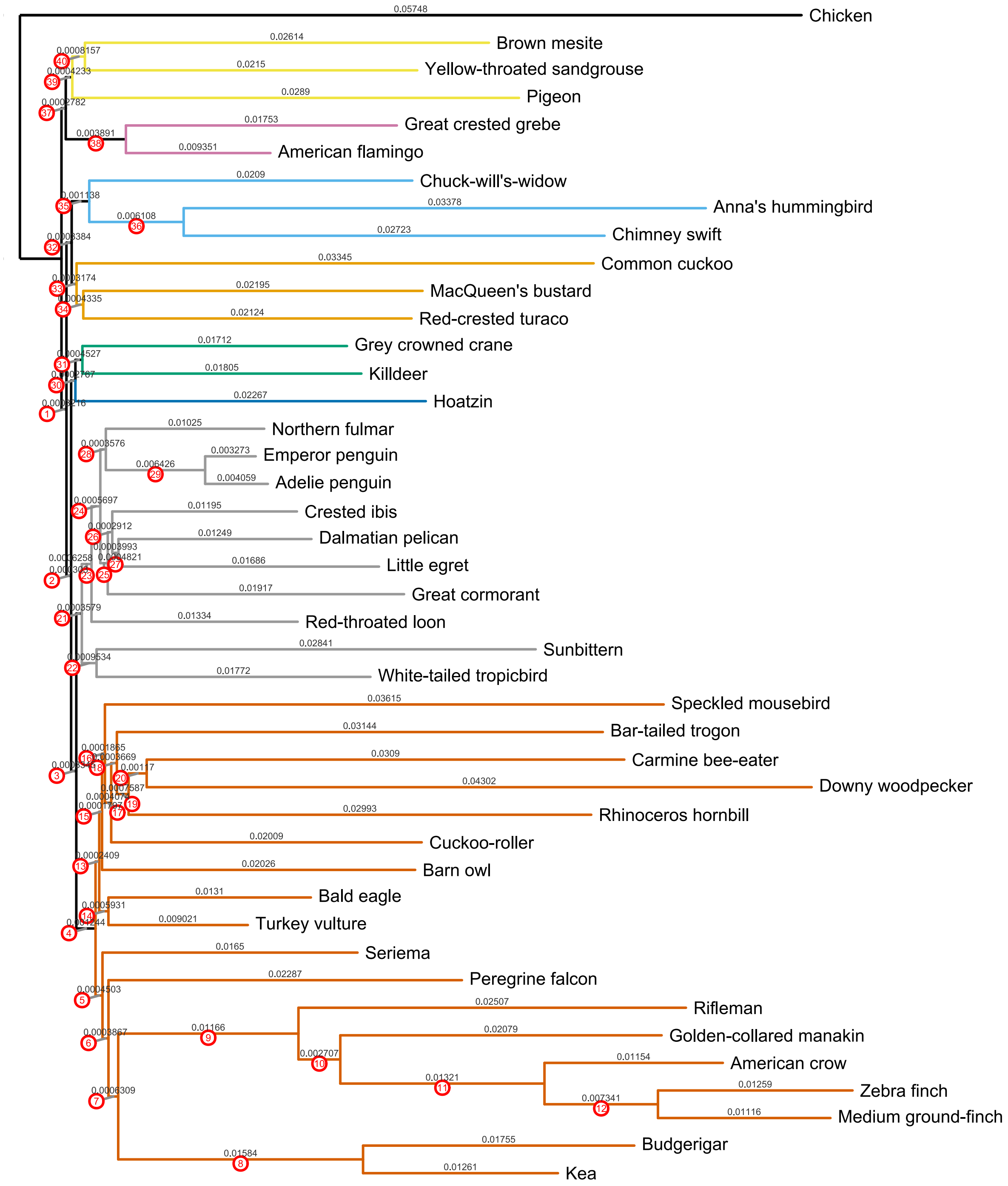


**Figure S23** Simulated species tree for the 43-taxon simulation, the interior edges are labeled by their lengths and split indices (drawn to scale).

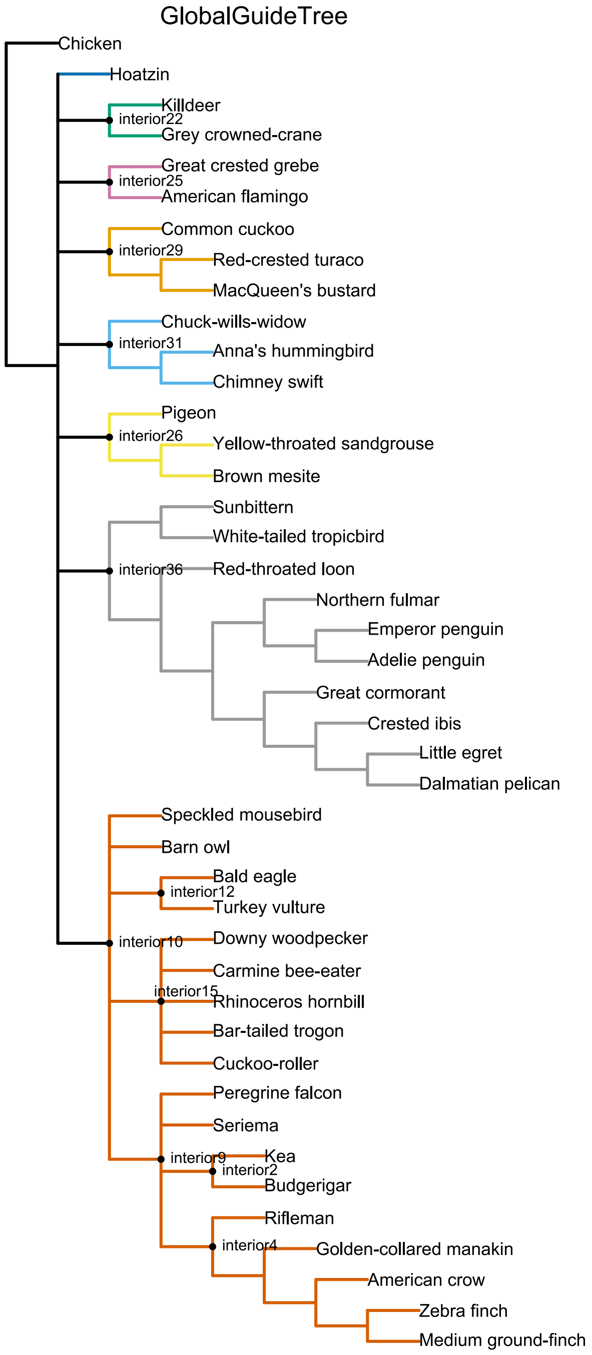


**Figure S24** Global guide tree used to define polytomies in 43-taxon simulated dataset. At the root level, the 8-polytomy outside the outgroup chicken consists of hoatzin, interior22, interior25, interior29, interior31, interior26, interior36 and interior10. Interior10 is a 5-polytomy, which consists of speckled mousebird, barn owl, interior12, interior15 and interior9. Interior15 is a 5-way polytomy consists of 5 taxa of Cavitaves. Interior9 is a 4-polytomy which consists of peregrine falcon, seriema, interior2 and interior4.

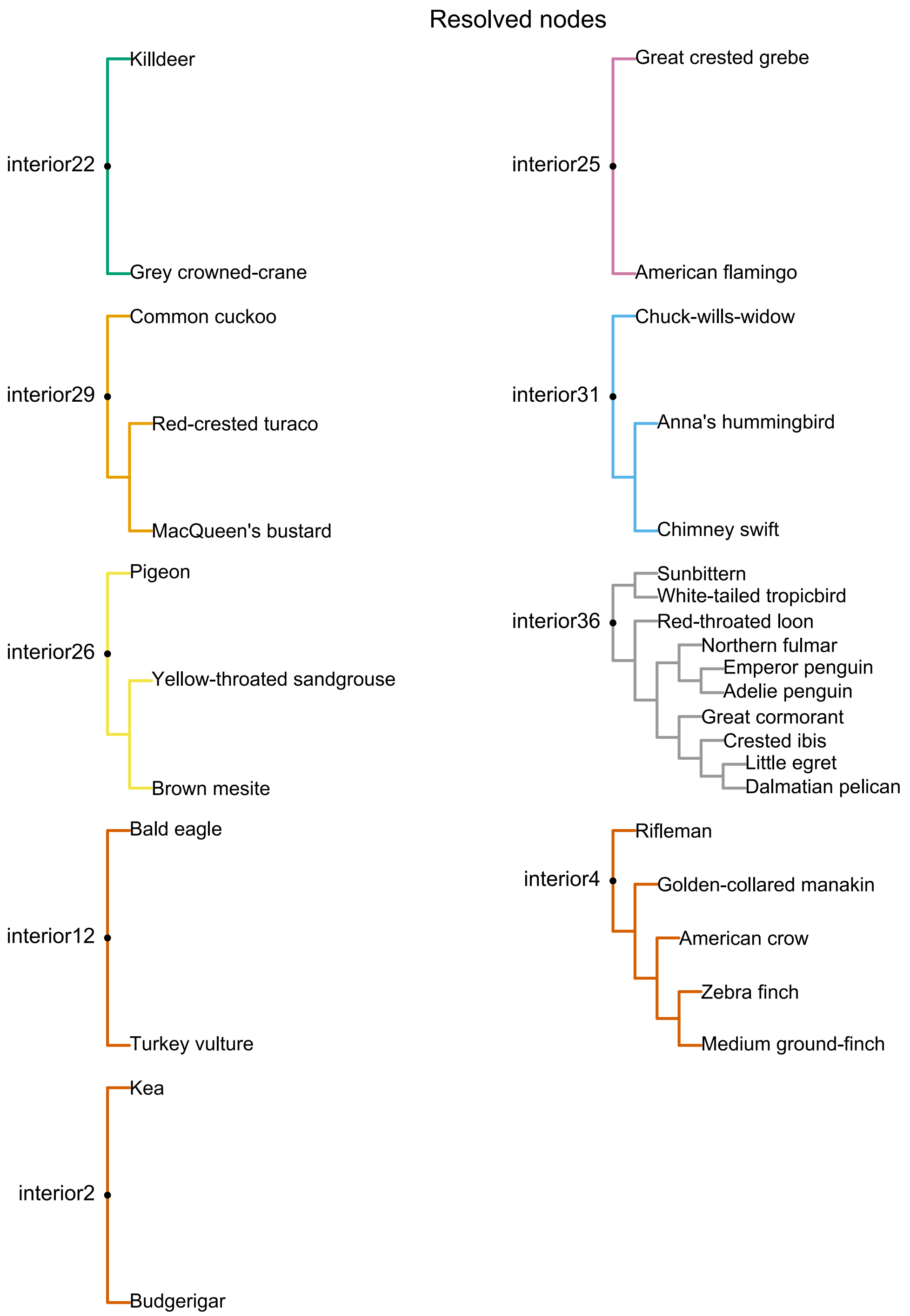


**Figure S25** Resolved nodes that were aligned first and reused in multiple-guide-trees alignments.

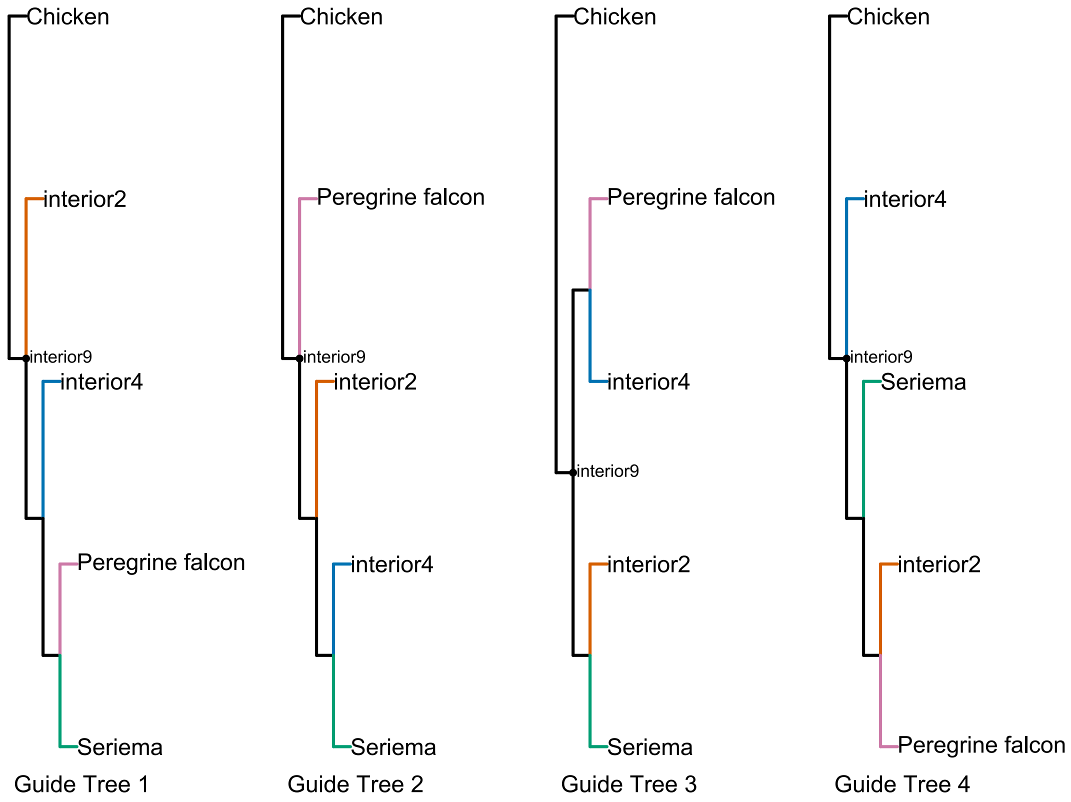


**Figure S26** Guide trees used in Australaves alignment.

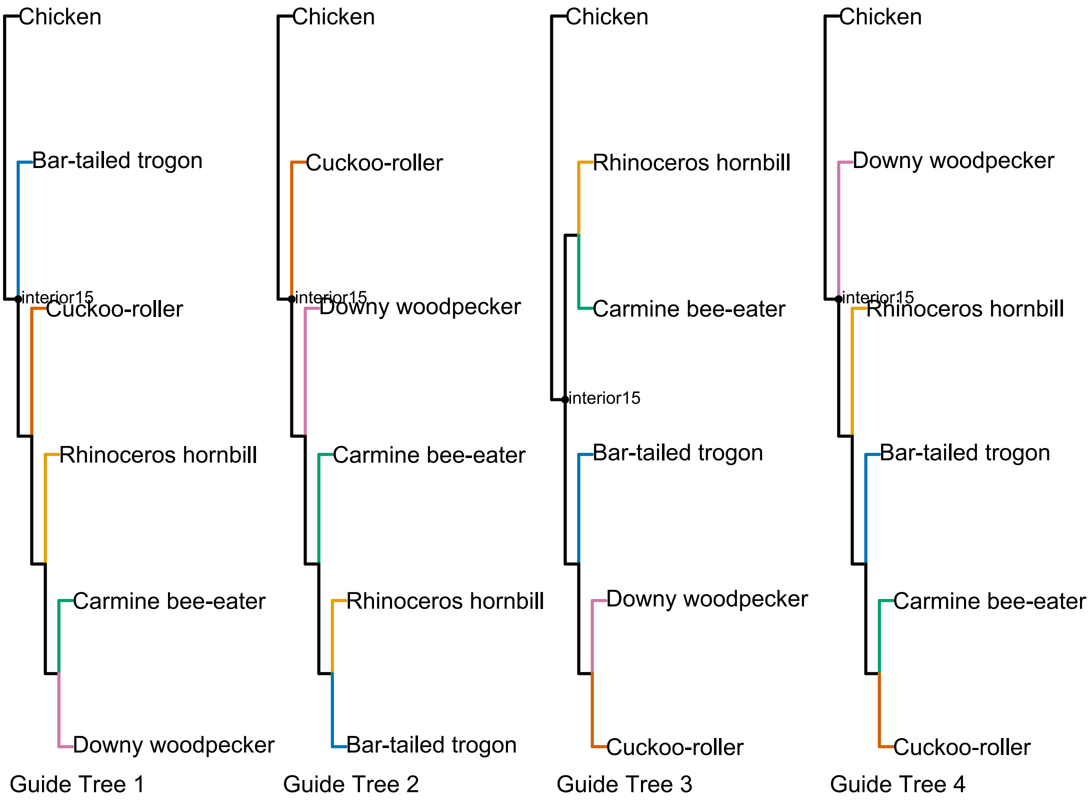


**Figure S27** Guide trees used in Cavitaves alignment.

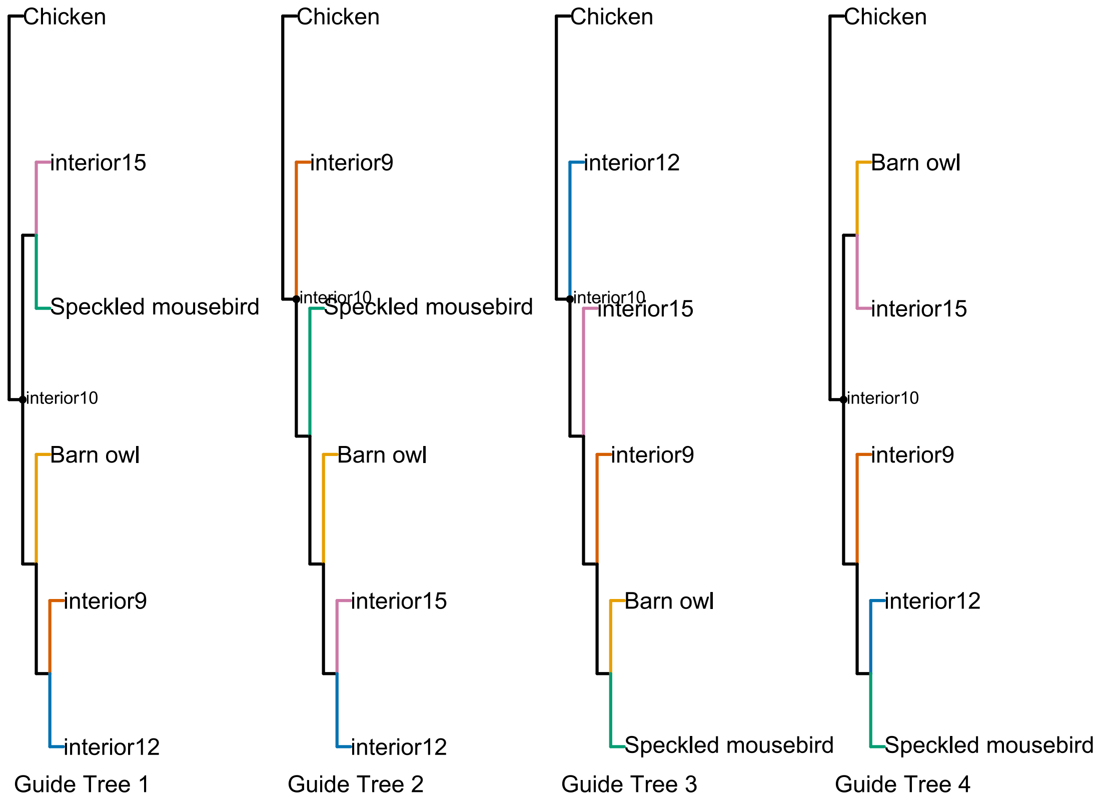


**Figure S28** Guide trees used in landbirds alignment.

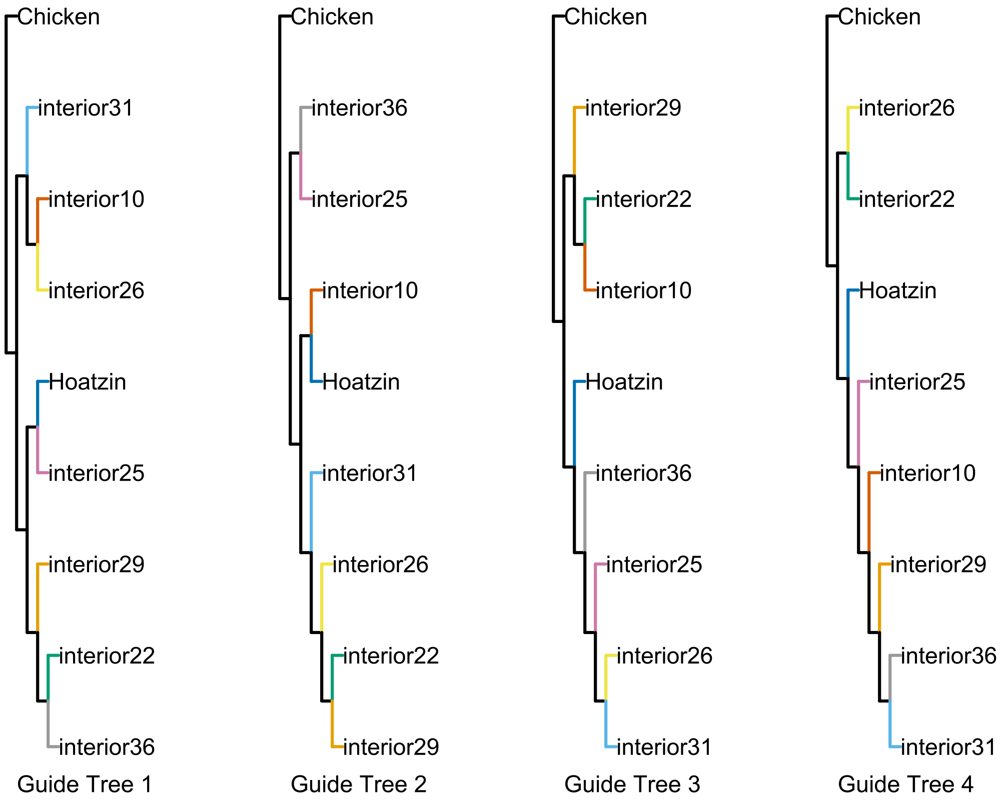


**Figure S29** Guide trees used in the root alignment.

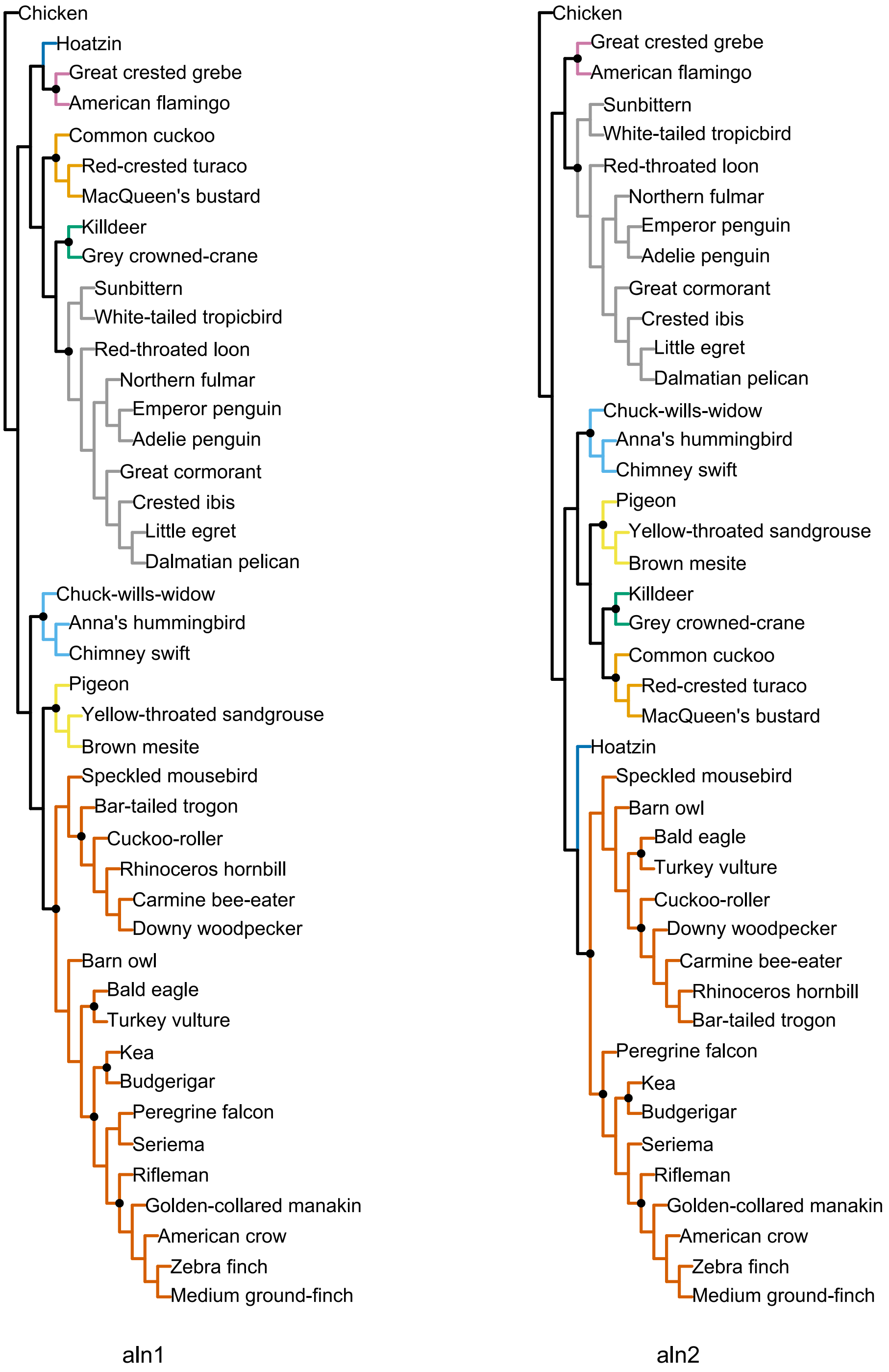


**Figure S30** Globally resolved guide trees used for comparison with the hierarchical consensus pipeline (aln1 and aln2).

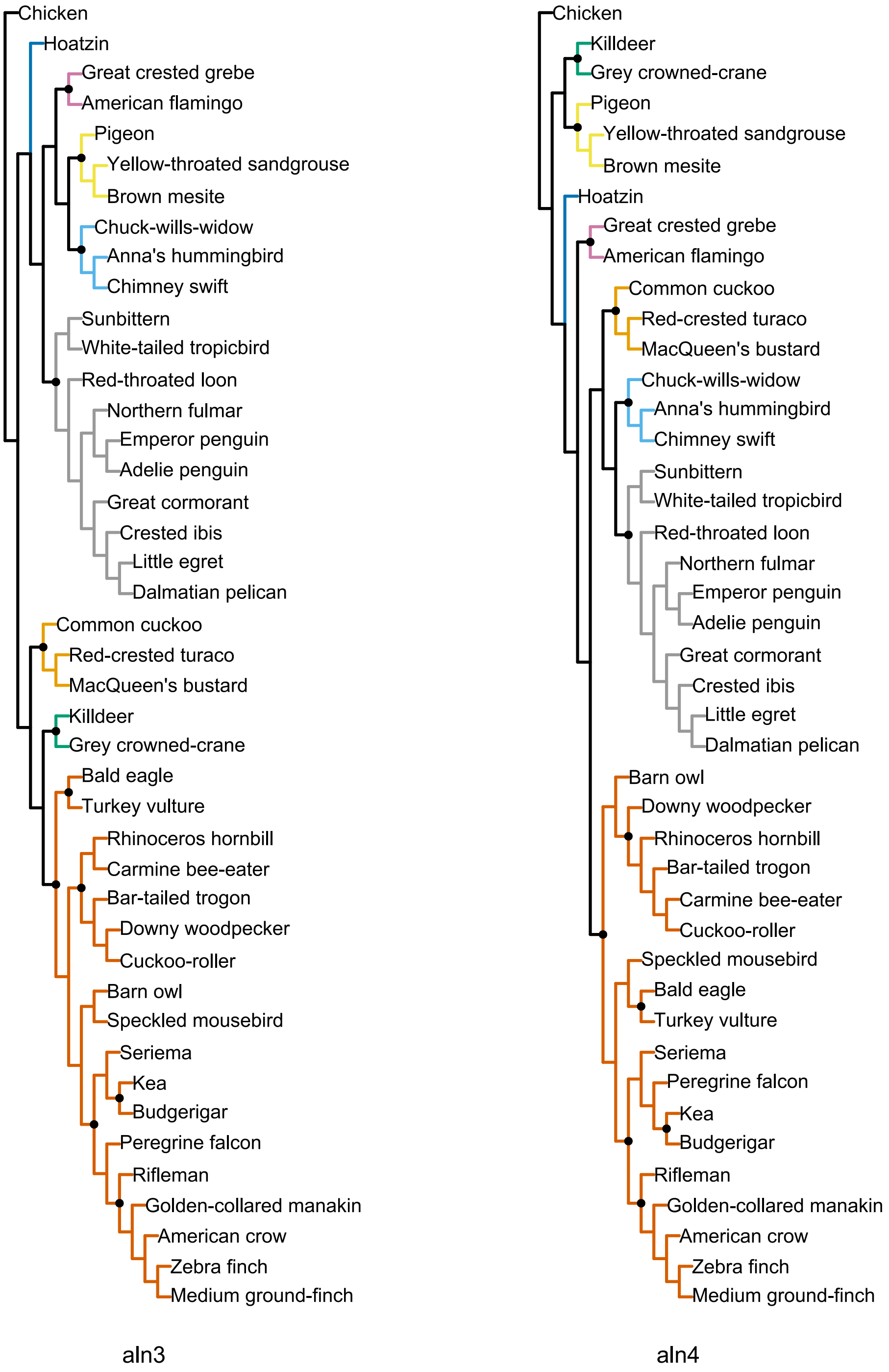


**Figure S30** (continued) Guide trees used for comparison with the hierarchical consensus pipeline (aln3 and aln4).

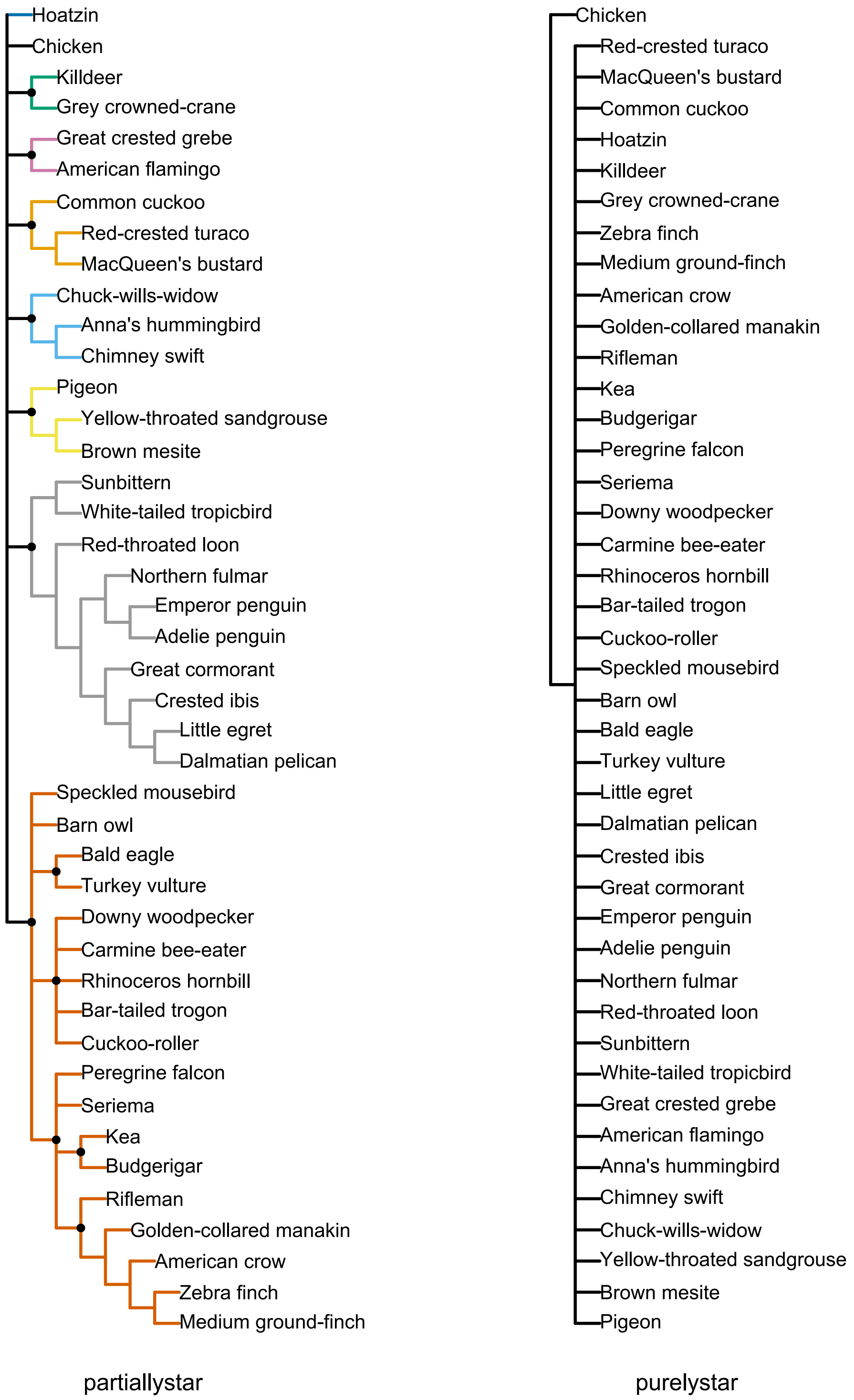


**Figure S30** (continued) Guide trees used for comparison with the hierarchical consensus pipeline (partiallystar and purelystar).

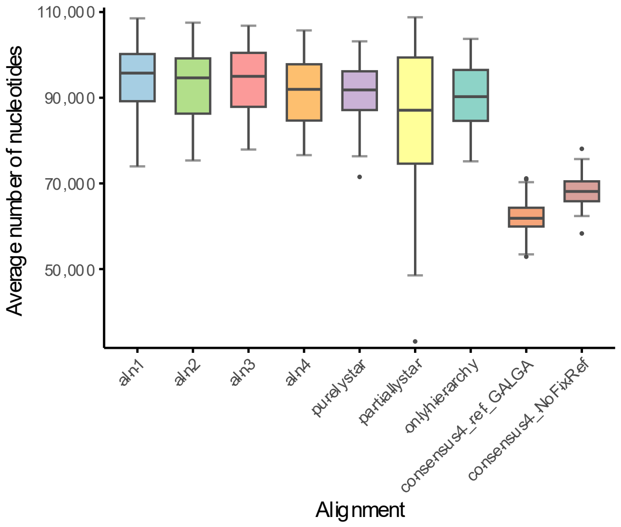


**Figure S31** Average number of nucleotides of all alignments including the one only going through the hierarchical pipeline and the one without a fixed global reference in 43-taxon simulation. "onlyhierarchy" refers to an alignment generated using the hierarchical pipeline with a single guide tree, without consensus construction. "consensus4_ref_GALGA" refers to the hierarchical consensus alignment using chicken (GALGA) as the fixed global reference, whereas "consensus4_NoFixRef" refers to the hierarchical consensus alignment without a fixed global reference.

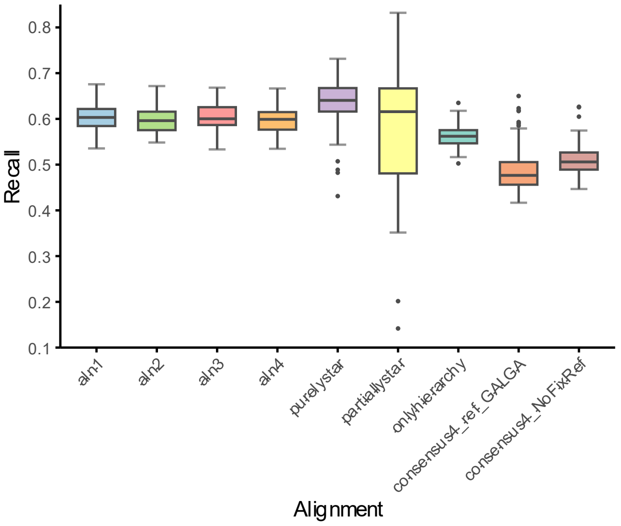


**Figure S32** Recall of 43-taxon simulated dataset using chicken as reference, also including the alignment only going through the hierarchical pipeline and the alignment without a fixed global reference. “onlyhierarchy” refers to an alignment generated using the hierarchical pipeline with a single guide tree, without consensus construction. “consensus4_ref_GALGA” refers to the hierarchical consensus alignment using chicken (GALGA) as the fixed global reference, whereas “consensus4_NoFixRef” refers to the hierarchical consensus alignment without a fixed global reference.

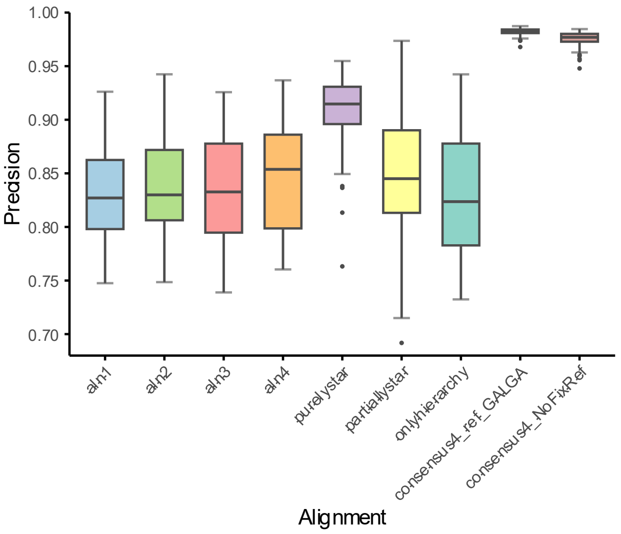


**Figure S33** Precision of 43-taxon simulated dataset using chicken as reference, also including the alignment only going through the hierarchical pipeline and the alignment without a fixed global reference. "onlyhierarchy" refers to an alignment generated using the hierarchical pipeline with a single guide tree, without consensus construction. "consensus4_ref_GALGA" refers to the hierarchical consensus alignment using chicken (GALGA) as the fixed global reference, whereas "consensus4_NoFixRef" refers to the hierarchical consensus alignment without a fixed global reference.

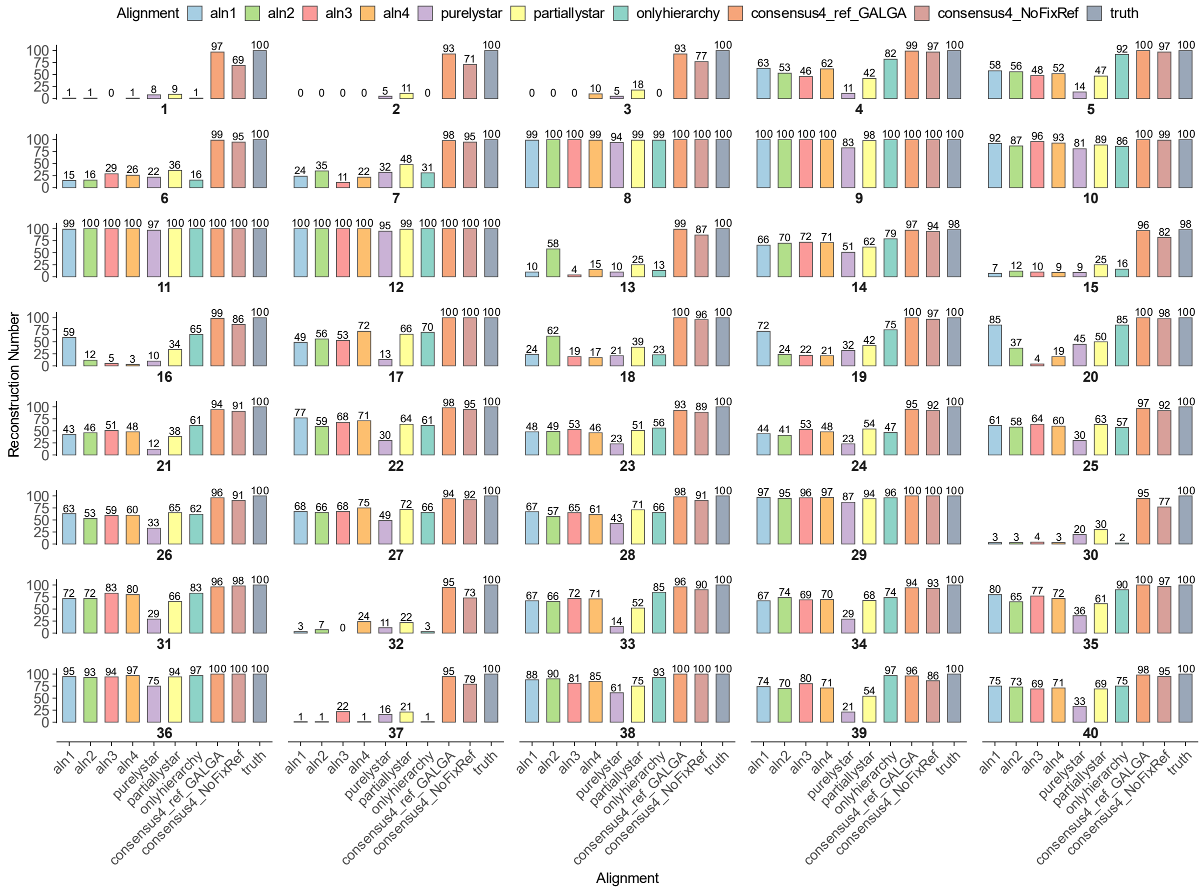


**Figure S34** Full split reconstruction results of 43-taxon simulated dataset using chicken as reference, also including the alignment only going through the hierarchical pipeline and the alignment without a fixed global reference. “onlyhierarchy” refers to an alignment generated using the hierarchical pipeline with a single guide tree, without consensus construction. “consensus4_ref_GALGA” refers to the hierarchical consensus alignment using chicken (GALGA) as the fixed global reference, whereas “consensus4_NoFixRef” refers to the hierarchical consensus alignment without a fixed global reference.

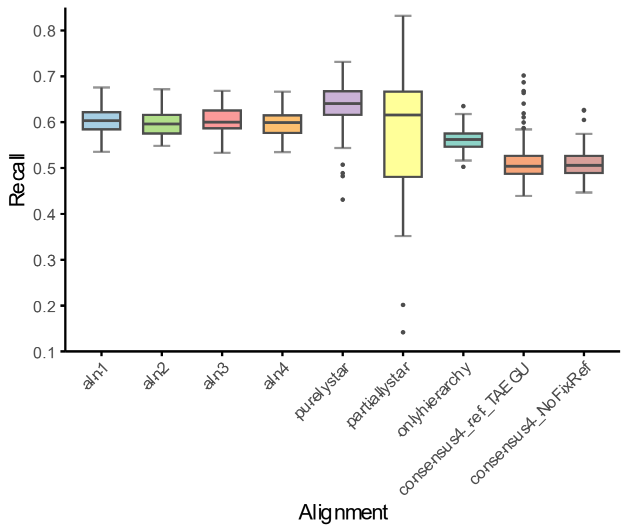


**Figure S35** Alignment recall of different strategies of 43-taxon simulated dataset using zebra finch as reference. "onlyhierarchy" refers to an alignment generated using the hierarchical pipeline with a single guide tree, without consensus construction. "consensus4_ref_TAEGU" refers to the hierarchical consensus alignment using zebra finch (TAEGU) as the fixed global reference, whereas "consensus4_NoFixRef" refers to the hierarchical consensus alignment without a fixed global reference.

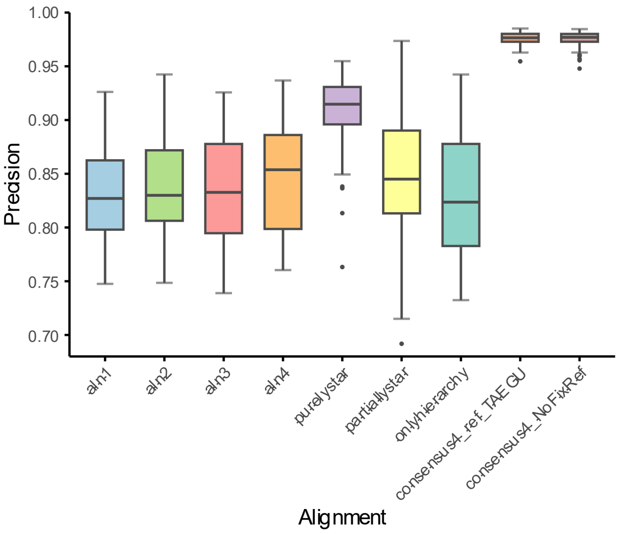


**Figure S36** Alignment precision of different strategies of 43-taxon simulated dataset using zebra finch as reference. “onlyhierarchy” refers to an alignment generated using the hierarchical pipeline with a single guide tree, without consensus construction. “consensus4_ref_TAEGU” refers to the hierarchical consensus alignment using zebra finch (TAEGU) as the fixed global reference, whereas “consensus4_NoFixRef” refers to the hierarchical consensus alignment without a fixed global reference.

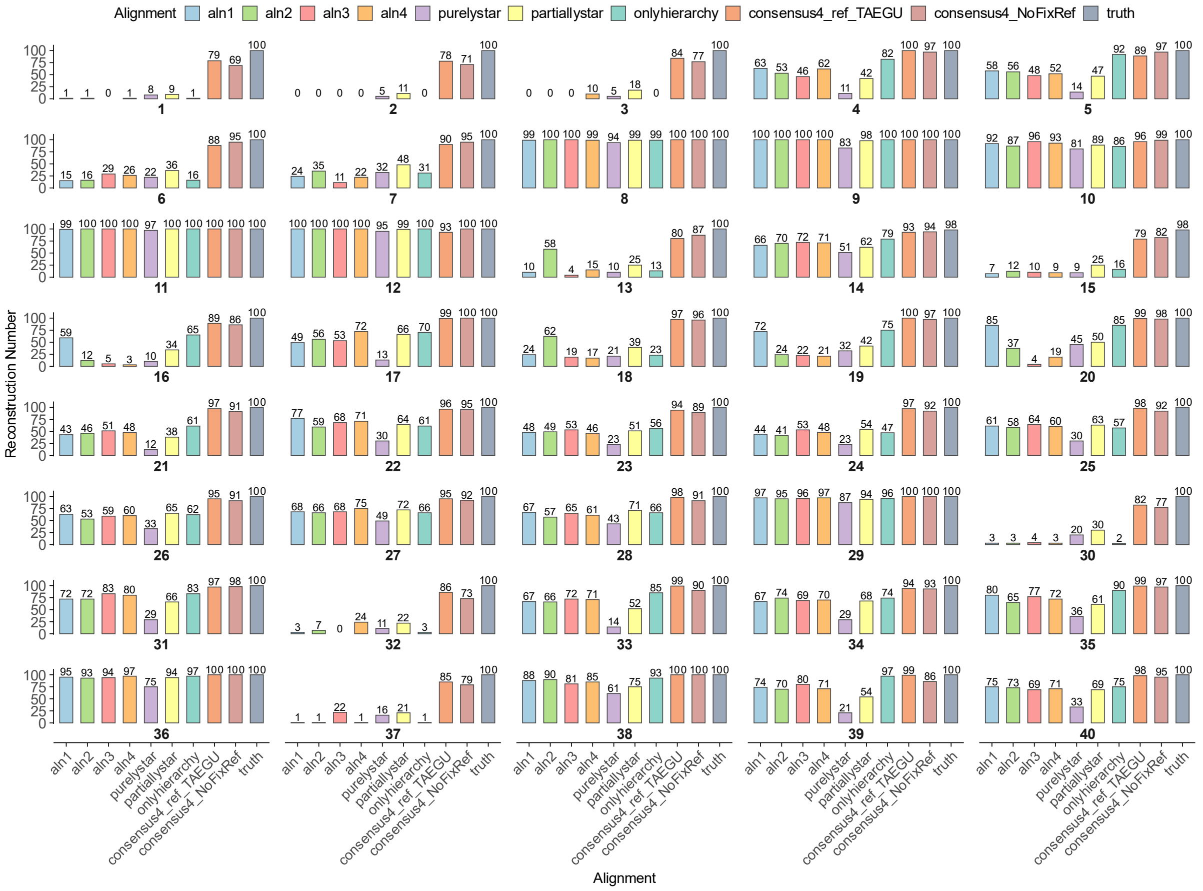


**Figure S37** Full split reconstruction results of 43-taxon simulated dataset using zebra finch as reference. “onlyhierarchy” refers to an alignment generated using the hierarchical pipeline with a single guide tree, without consensus construction. “consensus4_ref_TAEGU” refers to the hierarchical consensus alignment using zebra finch (TAEGU) as the fixed global reference, whereas “consensus4_NoFixRef” refers to the hierarchical consensus alignment without a fixed global reference.

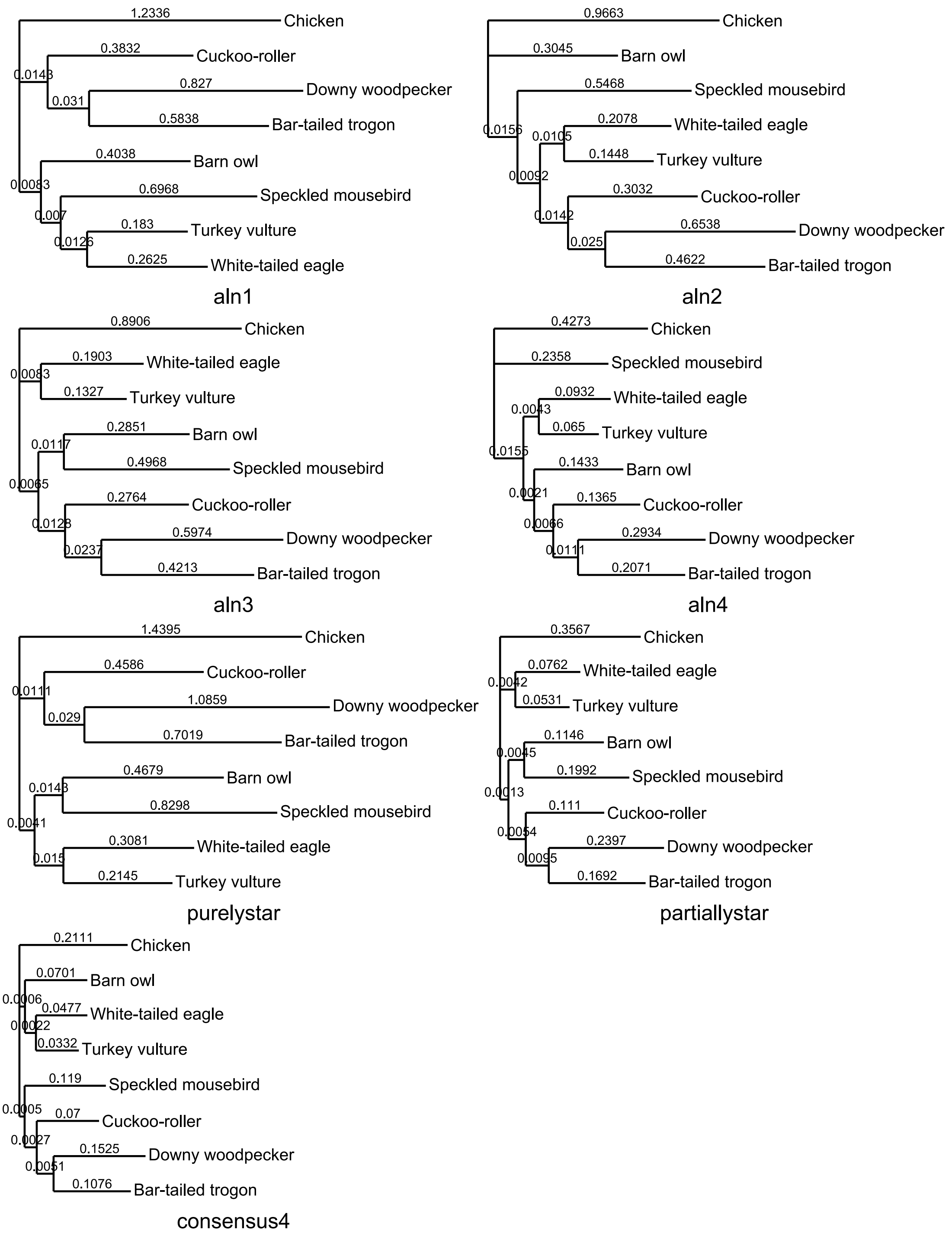


**Figure S38** Result trees inferred by IQ-TREE partition (10k) of the real 8-taxon experiment. Branch lengths were square-root transformed for visualization, whereas numerical labels indicate the original branch lengths estimated by IQ-TREE.

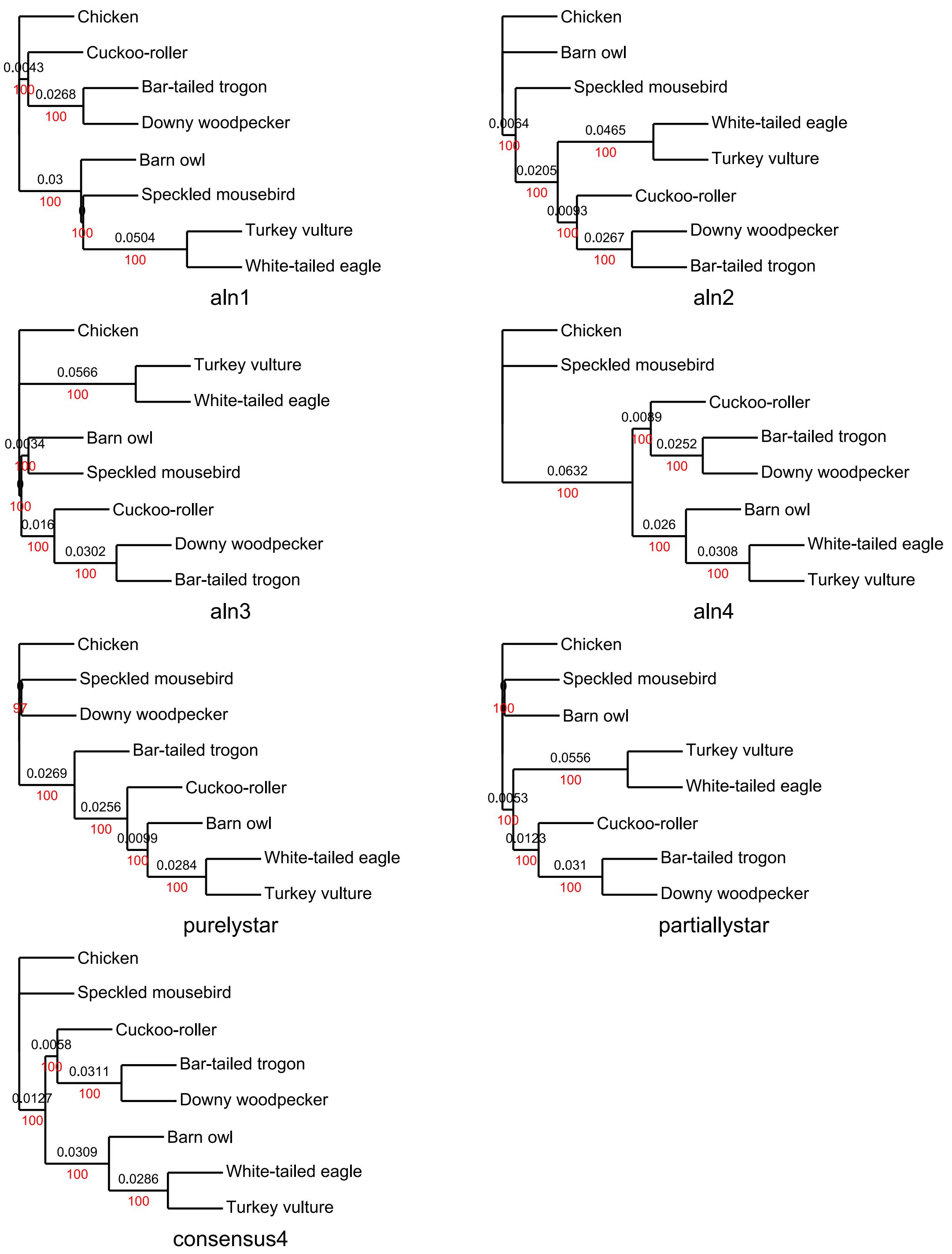


**Figure S39** Result trees inferred by CASTER-site of the real 8-taxon experiment. Internal branch lengths are drawn to scale and labeled with the original CASTER-site estimates, and terminal branch lengths were not provided by CASTER-site.

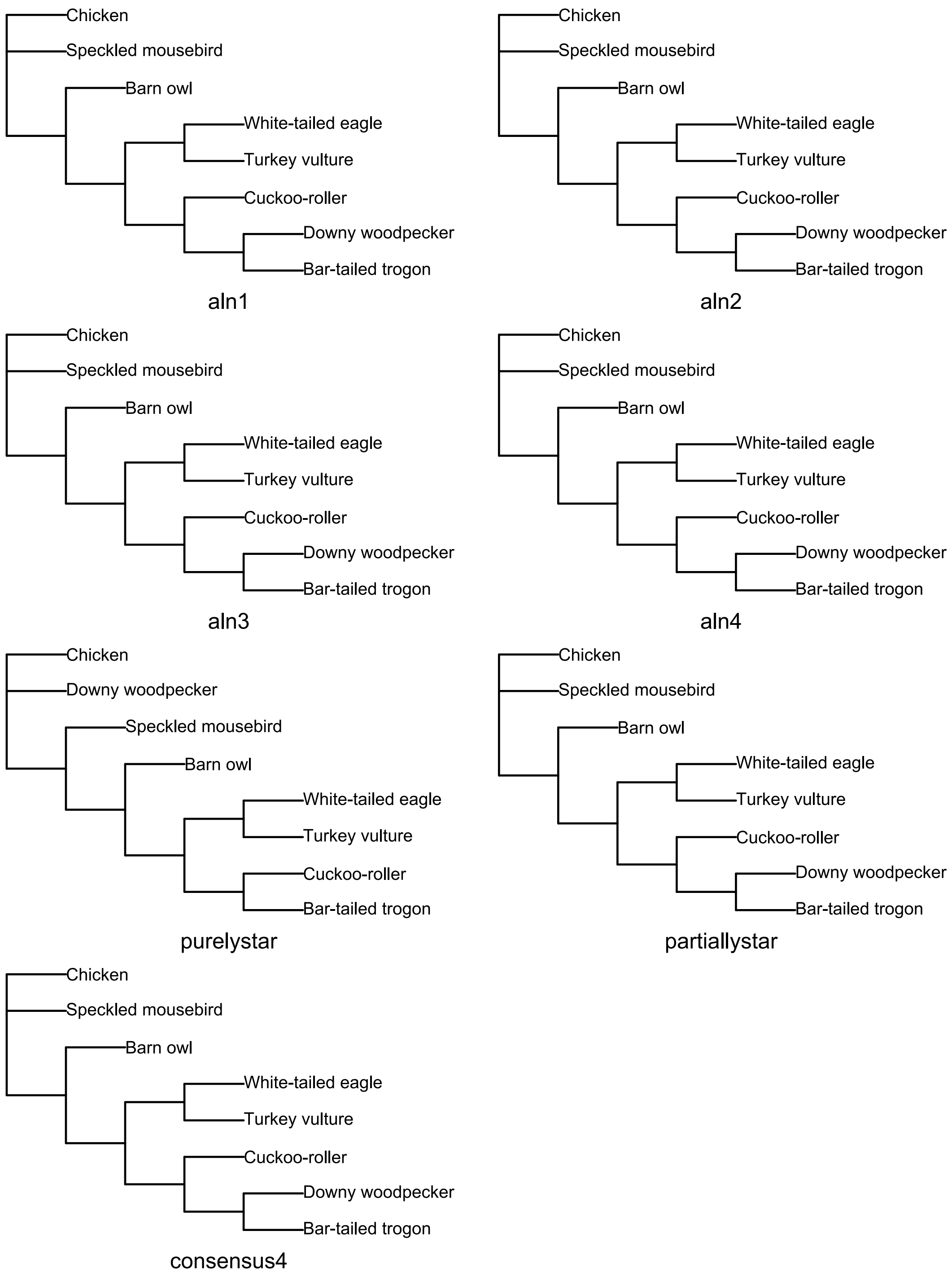


**Figure S40** Result trees inferred by maximum parsimony (PAUP) of the real 8-taxon experiment.

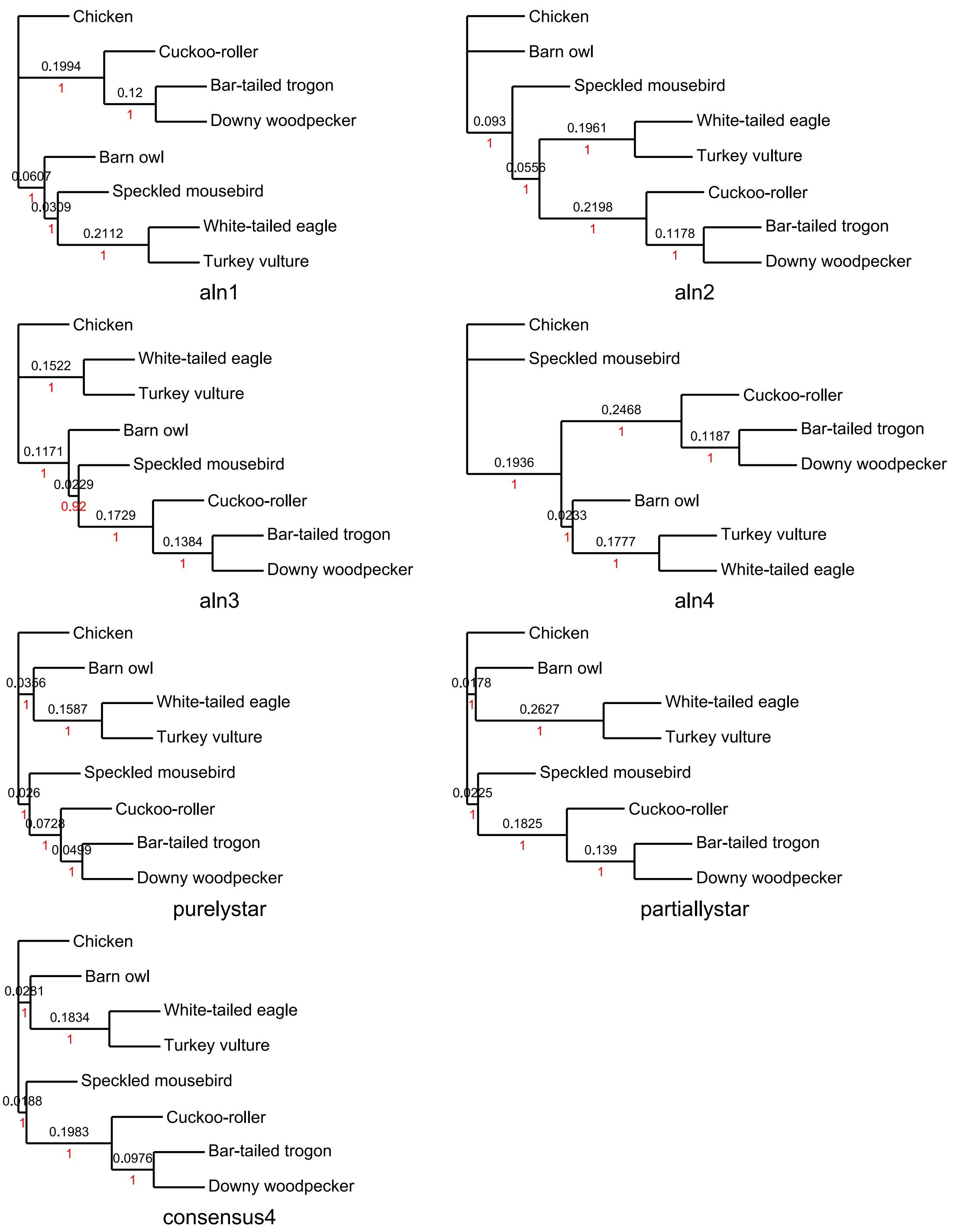


**Figure S41** Result trees inferred by loci-ASTRAL super tree method of the real 8-taxon experiment. Internal branch lengths are drawn to scale and labeled with the original ASTRAL estimates, and terminal branch lengths were not provided by ASTRAL.

| taxon | SeqType | SeqNumber | Length (bp) | TotalLength (bp) | Fraction_of_ MissingData |
|---|---|---|---|---|---|
| chicken | chromosome | 30 | 1,031,882,443 | 1,108,466,630 | 5.25% |
| | chromosome_r andom | 24 | 4,709,389 | | |
| | Un_random | 1 | 71,874,798 | | |
| zebra finch | chromosome | 34 | 1,021,453,031 | 1,235,794,146 | 0.84% |
| | chromosome_r andom | 33 | 39,115,800 | | |
| | Un | 1 | 175,225,315 | | |
| bar-tailed trogon | scaffolds | 46100 | 1,068,169,520 | 1,085,711,737 | 0.59% |
| | contigs | 104005 | 17,542,217 | | |
| cuckoo-roller | scaffolds | 44192 | 1,129,999,736 | 1,148,076,042 | 0.31% |
| | contigs | 104318 | 18,076,306 | | |
| seriema | scaffolds | 46631 | 1,129,974,933 | 1,145,705,463 | 0.52% |
| | contigs | 93214 | 15,730,530 | | |
| turkey vulture | scaffolds | 75632 | 1,109,559,534 | 1,179,513,848 | 1.21% |
| | contigs | 189975 | 69,954,314 | | |
| white-tailed eagle | scaffolds | 50606 | 1,135,974,242 | 1,144,733,688 | 0.78% |
| | contigs | 30616 | 8,759,446 | | |
| downy woodpecker | scaffolds | 5419 | 1,152,663,731 | 1,174,542,329 | 3.72% |
| | contigs | 80409 | 21,878,598 | | |
| speckled mousebird | scaffolds | 58808 | 1,072,904,594 | 1,086,303,309 | 0.66% |
| | contigs | 47009 | 13,398,715 | | |
| barn owl | scaffolds | 56508 | 1,121,074,466 | 1,138,815,065 | 1.02% |
| | contigs | 109584 | 17,740,599 | | |

**Table S1** Summary of genomes and assemblies used for whole-genome alignment

| chicken \| bar-tailed trogon \| cuckoo-roller \| turkey vulture | | | | | |
|---|---|---|---|---|---|
| TreeUsedToAlign | TreeTopoly | Alignment | FixedTreeTopology | Log-likelihood difference from the best score | Interior edge length |
| Tree1 | chicken & bar-tailed trogon | Alignment1 | Tree1 | 0 | 0.0056 |
| | | | Tree2 | 324963.3342 | 0.000045 |
| | | | Tree3 | 272209.5856 | 0.0016 |
| Tree2 | chicken & cuckoo-roller | Alignment2 | Tree1 | 401803.0193 | 0.00000073 |
| | | | Tree2 | 0 | 0.0052 |
| | | | Tree3 | 345006.7246 | 0.0017 |
| Tree3 | chicken & turkey vulture | Alignment3 | Tree1 | 469832.9205 | 0.00049 |
| | | | Tree2 | 472316.5757 | 0.000046 |
| | | | Tree3 | 0 | 0.0050 |

**Table S2** Log-likelihood differences from the best-scoring topology (best = 0) and optimized interior branch lengths for the three possible rooted topologies of three single-guide tree alignments of chicken, bar-tailed trogon, cuckoo-roller and turkey vulture.

| chicken \| seriema \| zebra finch \| turkey vulture | | | | | |
|---|---|---|---|---|---|
| TreeUsedToAlign | TreeTopoly | Alignment | FixedTree | Log-likelihood difference from the best score | Interior edge length |
| Tree1 | chicken & seriema | Alignment1 | Tree1 | 0 | 0.0057 |
| | | | Tree2 | 457352.43 | 0.00046 |
| | | | Tree3 | 449312.28 | 0.0013 |
| Tree2 | chicken & turkey vulture | Alignment2 | Tree1 | 277373.03 | 0.00062 |
| | | | Tree2 | 0 | 0.0041 |
| | | | Tree3 | 261665.59 | 0.0018 |
| Tree3 | chicken & zebra finch | Alignment3 | Tree1 | 1562633.77 | 2E-10 |
| | | | Tree2 | 1562079.54 | 0.00018 |
| | | | Tree3 | 0 | 0.016 |

**Table S3** Log-likelihood differences from the best-scoring topology (best = 0) and optimized interior branch lengths for the three possible rooted topologies of three single-guide tree alignments of chicken, seriema, zebra finch and turkey vulture.

<table>
<tr><th colspan="2">Alignment</th><th colspan="2">CPU time/hours</th><th colspan="2">Peak Memory/G</th><th>Average Nucleotides</th><th>Length</th><th>Locus Numbers</th></tr>
<tr><td colspan="2">aln1</td><td colspan="2">2771.76</td><td colspan="2">111.22</td><td>817,625,084</td><td>1,083,527,622</td><td>102,266</td></tr>
<tr><td colspan="2">aln2</td><td colspan="2">2618.45</td><td colspan="2">111.22</td><td>816,537,833</td><td>1,080,916,174</td><td>102,027</td></tr>
<tr><td colspan="2">aln3</td><td colspan="2">2674.05</td><td colspan="2">104.73</td><td>819,177,327</td><td>1,084,228,961</td><td>102,385</td></tr>
<tr><td colspan="2">aln4</td><td colspan="2">2779.59</td><td colspan="2">99.36</td><td>817,116,372</td><td>1,081,530,067</td><td>102,050</td></tr>
<tr><td colspan="2">purelystar</td><td colspan="2">3948.9</td><td colspan="2">439.05</td><td>816,538,091</td><td>1,070,119,704</td><td>100,054</td></tr>
<tr><td colspan="2">partiallystar</td><td colspan="2">2960</td><td colspan="2">110.45</td><td>818,611,570</td><td>1,083,778,114</td><td>102,289</td></tr>
<tr><td rowspan="7">consensus4</td><td>Cavitaves</td><td>606.41</td><td rowspan="7">4331.32</td><td>105.66</td><td rowspan="7">105.66</td><td rowspan="7">673,991,846</td><td rowspan="7">826,331,183</td><td rowspan="7">100,946</td></tr>
<tr><td>Accipitrimorphae</td><td>148.72</td><td>22.76</td></tr>
<tr><td>aln1 (5 taxa)</td><td>852.4</td><td>97.4</td></tr>
<tr><td>aln2 (5 taxa)</td><td>936.63</td><td>97.79</td></tr>
<tr><td>aln3 (5 taxa)</td><td>701.96</td><td>81.64</td></tr>
<tr><td>aln4 (5 taxa)</td><td>822.95</td><td>83.43</td></tr>
<tr><td>extract consensus+infer ancestor</td><td>262.25</td><td>62.8</td></tr>
</table>

**Table S4** Computational cost and alignment statistics for the real-data analyses.
